# Hypothalamic Representation of Aggressiveness across Mouse Strains

**DOI:** 10.64898/2026.06.10.731187

**Authors:** Xiuzhi Dai, Yifan Wang, Takashi Yamaguchi, Michael Genecin, Eyal Rozenfeld, Prakhar Dua, Bing Dai, Jing Cai, Dayu Lin

## Abstract

Aggression is an innate behavior conserved across species, serving as a critical means to compete for food, mating opportunities, and other essential resources. A central question in aggression research is the extent to which inter-individual variability in aggression is shaped by genetic factors. Here, we examine aggressive behaviors in naïve male mice across seven genetically defined strains and find large cross-strain differences. We find a tight correlation between aggressiveness and anxiety levels across strains, but not within the same strain, suggesting strong genetic control of both traits. Pharmacologically elevating anxiety in high-aggression strains reduces aggression, revealing a causal relationship between these behaviors. We further demonstrate that differences in the synaptic and cellular properties of neurons in the ventrolateral ventromedial hypothalamus (VMHvl) largely account for cross-strain variability in male aggression, and that chemogenetically increasing VMHvl excitability enhances attack behavior in a low-aggression strain. Together, these findings reveal the neuronal implementation of the genetic control of innate aggression level.

## Introduction

Aggression is an innate social behavior observed across a wide range of vertebrate and invertebrate species. It is an essential means of competing for resources, defending territory, and protecting oneself and one’s family. Given its critical role in survival, aggression is supported by developmentally hardwired circuits, ensuring its expression without learning^1^. Despite being innate, aggression differs drastically across individuals. In humans, some individuals actively seek the opportunity to fight, while others avoid any aggressive encounters^2,3^. Similar variability is observed in non-human primates^4^, mice^5–7^, rats^8,9^, and many other species^10–14^. Why does aggression have such a large cross-individual variability? Considerable evidence supports the important roles of experiential factors, such as winning, losing, and childhood adversity, in influencing aggression. For example, repeated winning increases the readiness to attack^15^, whereas defeat has the opposite effect^16,17^. Early life adversity often leads to increased aggression during adulthood^18–20^.

In addition to experiential factors, aggression level is also influenced by genetics. For example, mutation of monoamine oxidase A (MAOA), the enzyme needed to degrade monoamines, leads to hyper-aggression in males in both humans^21^ and mice^22^. Similarly, knocking out DAT, a dopamine transporter, leads to increased dopamine and exaggerated aggression^23^. Beyond specific gene mutations, twin studies estimate that approximately 50% of the variance in aggression can be explained by genetics^24^. In mice, it has long been recognized that wildtype animals with different genetic backgrounds, i.e., strain, show different aggression levels^25–30^. In fact, aggression researchers often take advantage of the different aggression levels across strains to stage fights with desired fighting outcomes^15,31,32^. Furthermore, researchers cross transgenic mice, e.g., Cre lines, with wildtype mice of high-aggression strains to increase the aggression level of the offspring^33^. These results suggest that aggression is determined collectively by “nature” and “nurture.”

Genetics presumably influences behaviors by altering the behavior-supporting neural circuits. To understand how genetically controlled aggression level is implemented neuronally, we first characterized aggressive behaviors of naïve males across seven commonly used laboratory mouse strains and examined their relationship with other emotional traits and neuroendocrine factors. We then examined the neural activation patterns in response to aggression-provoking cues across strains and found strain-specific responses of the ventrolateral part of the ventromedial hypothalamus (VMHvl), an indispensable region for driving attack^34,35^. Lastly, we examined VMHvl cell physiological properties across strains using in vitro patch-clamp recording and examined the causal relationship between VMHvl cell properties and aggression level.

## Results

### Different aggression levels across mouse strains

To investigate the genetic control of aggressive behaviors in wildtype animals, we performed inter-male resident-intruder tests (RI) using mice from seven commonly used laboratory strains, including CD-1 (CD1), Swiss Webster (SW), FVB, DBA/2 (DBA), C57BL/6 (C57), BALB/c (BC), and 129/SV-E (129) (**Fig. 1a**). All mice were purchased from Charles River, arrived around 6-8 weeks old and group-housed until 8 weeks old. They were then single-housed for two weeks before the RI test. At the time of the test, all subject animals were 10 weeks old. During the test, we introduced a group-housed male intruder (BC intruders for all strains except for BC; C57 intruders for BC) into the home cage of the test mouse for 10 minutes (**Fig. 1b**). The intruders were group-housed and previously defeated and never initiated attacks towards the resident test mice throughout the study. **Figure 1c** shows the investigation and attack events during the RI tests of three representative mice of each strain: the second-most, middle, and second-least aggressive mice.

**Figure 1.**
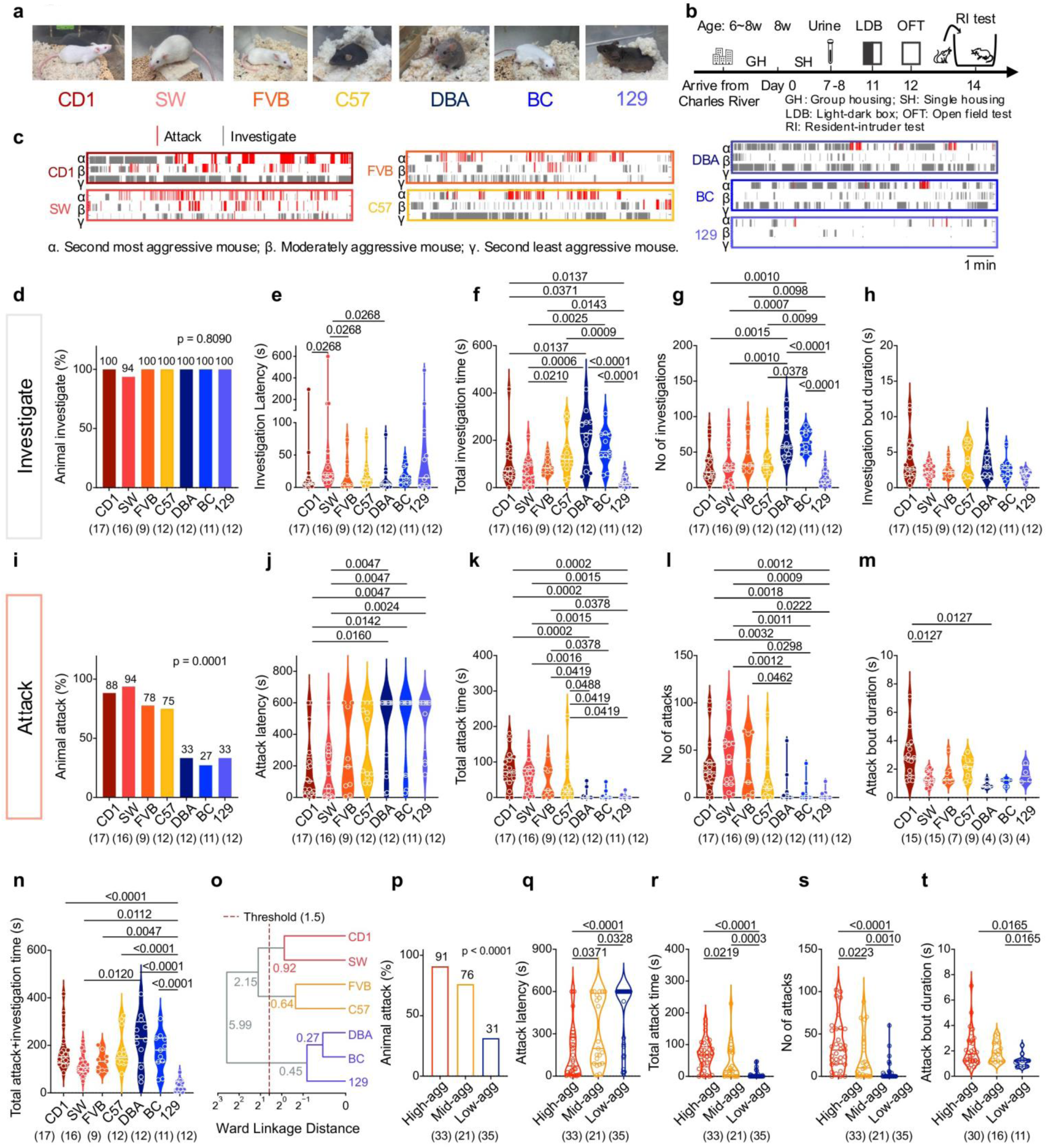
Male aggression levels of seven mouse strains. **(a)** Representative images of seven strains of mice used in the study. **(b)** Experimental timeline for urine collection, light-dark box test (LDB), open-field test (OF), and resident-intruder test (RI). All mice were purchased from Charles River company at the age of 6-8 weeks. Day 0 represents when the test mice were 8 weeks old. **(c)** Representative raster plots showing investigation (gray) and aggressive behaviors (red) of male mice of various strains during the 10-min RIT. Three mice per strain are shown. α: second most aggressive; β: moderately aggressive; γ: second least aggressive. Rank is based on attack duration. **(d)** Percentage of animals that investigated across strains. **(e–h)** Average **(e)** investigation latency, **(f)** total investigation duration, **(g)** number of investigation events, and **(h)** investigation duration per bout across individuals of all strains. **(i)** Percentage of animals that attacked across strains. **(j–m)** Average **(j)** attack latency, **(k)** total attack time, **(l)** number of attacks, and **(m)** attack duration per bout across individuals of all strains. **(n)** Average combined attack and investigation time across individuals of all strains. **(o)** Hierarchical clustering across strains. CD1 and SW are clustered as high-aggressive strains, FVB and C57 are clustered as middle-aggressive strains, and DBA, BC and 129 are clustered as low-aggressive strains. The numbers on the clustering tree indicate the Ward linkage distances (i.e., the increase in within-cluster variance) at each merge. **(p)** Percentage of animals that attacked in low-, mid-, and high-aggression groups. **(q–t)** Average **(q)** attack latency, **(r)** total attack time, **(s)** attack number, and **(t)** attack duration per bout of all animals in low-,mid- and high-aggression groups. Numbers in parentheses indicate animal numbers. Circles represent data of individual animals. Solid lines in (**e-h, j-m, n, q-t**) represent the median for each group, while dashed lines indicate quartiles. Color in **(d-n)** indicates strain identity while color in **(o-t)** indicate the aggression level. **(d, i, p)** Fisher’s exact test; **(e–h, j–n, q-t)** One-way ANOVA for normally distributed datasets or Kruskal–Wallis test for non-normally distributed datasets with FDR-corrected post hoc comparisons (Benjamini–Krieger–Yekutieli method). All statistical tests are two-tailed. Exact p- or q-value is shown if ≤ 0.05. Otherwise, p- or q-value is unspecified. See **Supplementary Table 1** for additional statistical details.

Upon intruder introduction, all animals quickly investigated the intruder with the exception of one SW mouse that initiated rapid and sustained attacks, leaving little opportunity for investigation (**Fig. 1d-e**). Across strains, DBA and BC mice investigated the intruder the most, spending on average 3-4 minutes investigating and initiating investigatory bouts more frequently compared to other strains (**Fig. 1f, g**). C57 mice investigated the intruder for a bit over 2 minutes, while CD1, SW, and FVB spent 1-2 minutes investigating (**Fig. 1f, g**). 129 is a clear outlier: all 129 animals investigated the intruder for less than 1 minute with an average investigation duration of approximately 30 seconds (**Fig. 1f, g**). The duration of individual investigation bouts was comparable across strains. (**Fig. 1h**)

After bounts of investigation, most animals attacked the intruder, although the probability of attacking varied markedly across the strains (**Fig. 1i**). CD1 and SW mice showed consistent aggression. 15/17 CD1 and 15/16 SW mice attacked the intruder (**Fig. 1i**). Most CD1 and SW mice initiated attack within 3 minutes and attacked for more than 60 seconds (**Fig. 1j, k**). SW attacked more frequently, while CD1 mice tended to have less frequent but longer attack bouts (**Fig. 1l, m**). In contrast, DBA, BC, and 129 showed minimal aggression. Only ∼30% of animals in these three strains attacked at all (**Fig. 1i**). For DBA, BC, and 129 animals that did attack, the attack duration was generally low, typically comprising several brief bouts (**Fig. 1c, k-m),** and the attack latency was variable (**Fig. 1j**). With regard to C57 and FVB, the majority of the animals (∼75%) attacked the intruder, and the attack duration averaged around 40s, although there was a high variability among individuals (**Fig. 1i-m**). Notably, the summation of attack duration and investigation duration, a measurement of overall social engagement, is relatively comparable across strains, with the obvious exception of 129, which is significantly lower than all other strains (**Fig. 1n**). These results suggest that the exceptionally low aggression of 129 mice may partly stem from its low social interest or high social anxiety.

Based on the percentage of animals that attacked the intruder, average attack latency and duration of each strain, we found that these seven strains of mice can be clustered into three distinct groups using the elbow method (**Supplementary Figure 1a**). Hierarchical clustering and k-means clustering further revealed high (CD1 and SW), moderate (C57 and FVB), and low (DBA, 129, and BC) aggression groups (**Figure 1o, Supplementary Figure 1b**). Animals in these groups differed significantly in their probability of attack, attack latency, duration, and frequency (**Fig. 1o-s**). Animals in high- and moderate-aggression strains exhibited longer attack bouts compared to those in low-aggression strains (**Fig. 1t**).

The body weight differed across these 7 strains (**Supplementary Fig. 2a**). CD1 and SW mice were similar in weight and heavier than all other strains. FVB mice were lighter than CD1 and SW but heavier than the remaining strains, whereas C57, BC, DBA, and 129 mice did not differ significantly from one another in body weight. Across strains, body weight is positively correlated with aggression level: the most aggressive strains, CD1 and SW, are also the heaviest (**Supplementary Fig. 2b**). However, within each strain, body weight is not correlated with aggression level (**Supplementary Fig. 2c, d**). In other words, heavier animals are not necessarily more aggressive than lighter animals of the same genetic background.

The body weight of BC and C57 adult intruders is lower than that of CD1 and SW residents but similar to or slightly higher than that of residents from other strains (**Supplementary Figure 2a**). Because CD1 and SW showed the highest level of aggression and were the only strains significantly heavier than the intruders, this raised the possibility that aggression level varies with the relative size of the intruder and resident. To test this, we compared the aggression level of C57, BC, and 129 males towards three different intruders: juvenile males, size-matched, group-housed, same-strain adult males, and our standard male intruders. The juvenile intruders were substantially smaller and lighter (mean ± standard error (SEM): 13.56 ± 0.54 g, n = 12) than all adult residents, regardless of strain (**Supplementary Fig. 2b**). However, we observed no significant difference in aggression toward the different types of intruders in the three tested strains (**Supplementary Fig. 2e-s**). For example, among 129 male residents, 4/12 (33%) attacked adult BC male intruders, 4/10 (40%) attacked adult 129 male intruders, and 2/10 (20%) attacked juvenile intruders (**Supplementary Fig. 2p**). These results indicate that in our study, aggression level is determined largely by intrinsic aggressiveness rather than by the size of the intruder.

Female aggression in mice varies with reproductive states: it is low during the virgin state and high during lactation^36^. Nevertheless, variation in aggression level across strains has also been reported^36,37^. To investigate the consistentcy of inter-strain differences in aggression between sexes, we performed RI tests using single-housed, virgin SW and C57 female residents exposed to juvenile BC male intruders (**Supplementary Fig. 3**). Consistent with the results in males, a higher proportion (6/10) of SW females attacked the intruder than C57 females (2/10) did (**Supplementary Fig. 3b)**. For SW females that initiated attack, the attack latency was relatively short (< 3 minutes), while the attack duration was variable, ranging from a few seconds to one minute (**Supplementary Fig. 3c-3f**). In contrast, the two aggressive C57 females only attacked briefly (<2s) (**Supplementary Fig. 3d**). These findings indicate that cross-strain differences in aggression are largely preserved between sexes.

**Figure 2.**
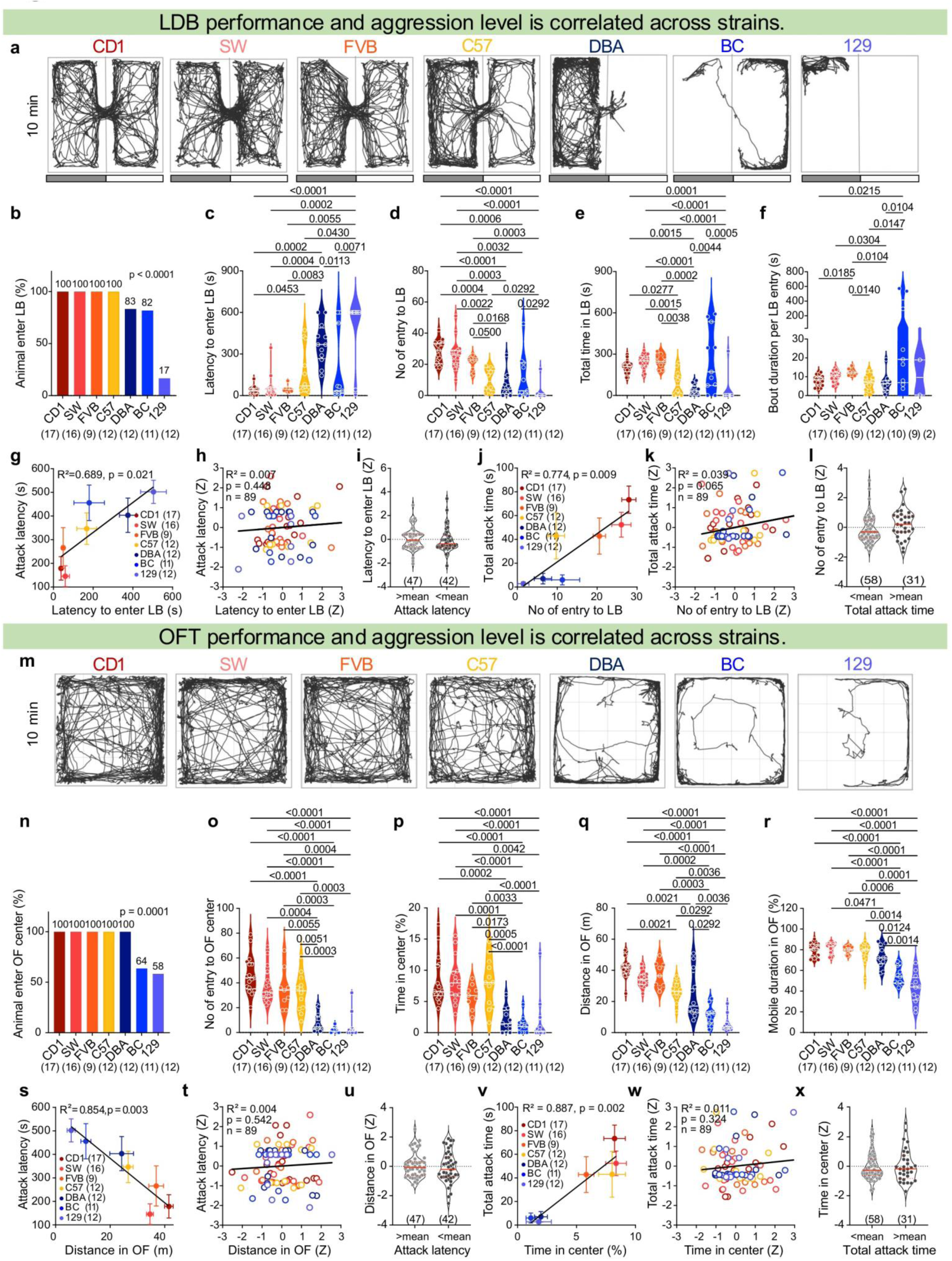
Anxiety and innate aggressiveness are negatively correlated across strains. **(a)** Tracking results from representative animals of various strains during the 10-minute light-dark box test (LDB). The representative animal is the middle-ranked animal of each strain based on latency to enter the light box. Gray and white bars indicate the dark and light sides, respectively. **(b)** Percentage of animals of each strain entering the light box. LB: light box. **(c–f)** Average **(c)** latency to enter the light box, **(d)** total number of light box entries, **(e)** total time spent in the light box, and **(f)** bout duration per light box entry of all animals in each strain. **(g)** Correlation between mean attack latency and latency to enter the light box across strains. **(h)** Correlation between strain-normalized attack latency and latency to enter the light box across individuals of all strains. **(i)** The light box entry latency between animals with attack latency below or above the strain mean. **(j)** Correlation between mean total attack time and total number of light box entries across strains. **(k)** Correlation between strain-normalized total attack time and total number of light box entries across individuals of all strains. **(l)** The light box entry number between animals with total attack time below or above the strain mean. **(m)** Tracking results from representative animals of various strains during the 10-minute open field test (OFT). The chosen animal is the middle-ranked animal of each strain based on total distance traveled in the open field. **(n)** Percentage of animals of each strain that enter the open field center. OF: open field. **(o–r)** Mean **(o)** total number of open field center entry, **(p)** percentage of time spent in the center of the open field, **(q)** distance traveled in the open field, and **(r)** mobile duration in the open field. **(s)** Correlation between mean attack latency and distance traveled in the open field across strains. **(t)** Correlations between strain-normalized attack latency and distance traveled in the open field across individuals of all strains. **(u)** Distance traveled in the open field between animals with attack latency above or below the strain mean. **(v)** Correlation between mean total attack time and percentage of time spent in the center. **(w)** Correlation between strain-normalized total attack time and percentage of time in the center across individuals of all strains. **(x)** Distance traveled in the open field between animals with total attack time below or above the strain mean. Numbers in parentheses indicate animal numbers. Color in (**b-h, j-k, n-t, and v-w**) indicates strain identity. Circles represent data of individual animals. Solid line in (**c-f, i, l, o-r, u, x**) represents the median for each group, while dashed lines indicate quartiles. Solid circles represent the strain mean, and error bars represent ± SEM in (**g, j, s, v**). **(b, n)** Fisher’s exact test; **(c-f, o-r)** One-way ANOVA for normally distributed datasets or Kruskal–Wallis test for non-normally distributed datasets with FDR-corrected post hoc comparisons (Benjamini–Krieger–Yekutieli method); **(i, l, u, x)** Unpaired t test for normally distributed datasets or Mann–Whitney U test for non-normally distributed datasets; **(g, h, j, k, s, t, v, w)** Linear regression and Pearson correlation with reported *R²* and *p*-values. All statistical tests are two-tailed. Exact p- or q-value is shown if ≤ 0.05. Otherwise, p- or q-value is unspecified. See **Supplementary Table 1** for additional statistical details.

**Figure 3.**
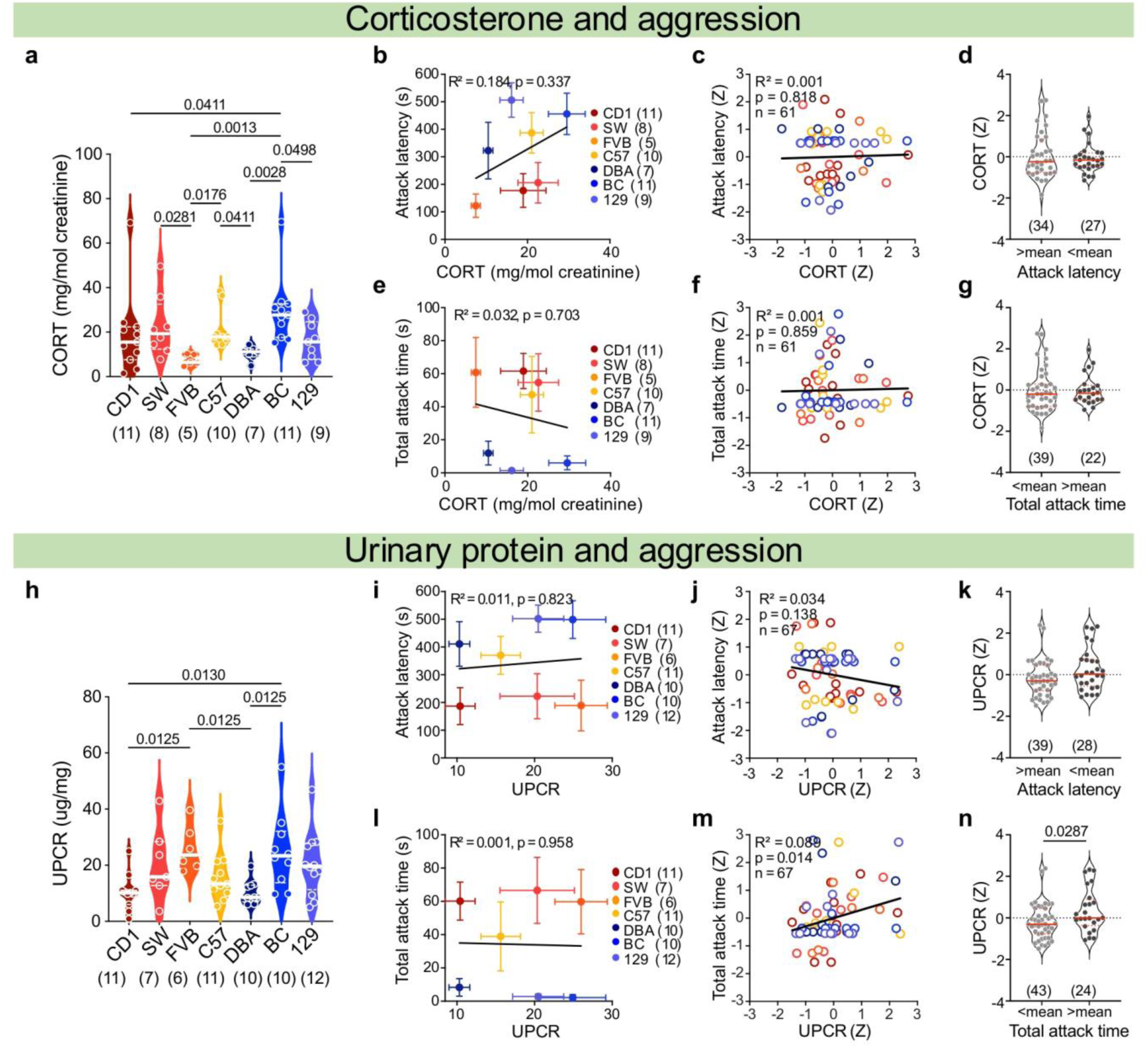
Relationship between aggression, corticosterone (CORT), and urinary protein levels. **(a)** Average urinary CORT concentration (normalized by creatinine) of various strains of male mice. **(b)** Correlation between CORT concentration and attack latency across strains. **(c)** Correlation between strain-normalized CORT concentration and attack latency across individuals. **(d)** CORT concentration of individuals with attack latency above and below the strain average. **(e)** Correlation between CORT concentration and total attack time across strains. **(f)** Correlation between strain-normalized CORT concentration and total attack time across individuals. **(g)** CORT concentration of individuals with total attack time below and above the strain average. **(h)** Normalized urinary protein concentrations of various strains of male mice. UPCR: urine protein-to-creatinine ratio. **(i)** Correlation between average urinary protein concentration and attack latency across strains. **(j)** Correlation between strain-normalized urinary protein concentration and attack latency across individuals. **(k)** Urinary protein concentration of individuals with attack latency above and below the strain average. **(l)** Correlation between average urinary protein concentration and total attack time across strains. **(m)** Correlation between strain-normalized urinary protein concentration and total attack time across individuals. **(n)** Urinary protein concentration of individuals with total attack time below and above the strain average. Numbers in parentheses indicate animal numbers. Color in (**a-c, e-f, h-j, and l-m**) indicates strain identity. Circles represent data of individual animals. Solid line in (**a, d, g, h, k, n**) represents the median for each group, while dashed lines indicate quartiles. Solid circles represent the strain mean, and error bars represent ± SEM in (**b, e, i, l**). **(a, h)** One-way ANOVA for normally distributed datasets or Kruskal–Wallis test for non-normally distributed datasets with FDR-corrected post hoc comparisons (Benjamini–Krieger–Yekutieli method); **(d, g, k, n)** Unpaired t test for normally distributed datasets or Mann–Whitney U test for non-normally distributed datasets; **(b, c, e, f, i, j, l, m)** Linear regression and Pearson correlation with reported *R²* and *p*-values. All statistical tests are two-tailed. Exact p-or q-value is shown if ≤ 0.05. Otherwise, p- or q-value is unspecified. See **Supplementary Table 1** for additional statistical details.

### Anxiety and aggression are negatively correlated across strains

Anxiety is an emotional trait linked to the genetic background of an individual^38^. Consistent with previous reports^39–41^, we found that anxiety levels differ dramatically among the seven strains of test mice, as reflected by their performance in the light-dark box (LDB) and open field test (OFT) (**Fig. 2a, m**). In the LDB test, nearly all CD1, SW, and FVB mice entered the light chamber within the first minute and spent over one-third of their time actively exploring it (**Fig. 2b-e**). C57 mice behaved more variably: some entered the light chamber quickly (<60s), while others did not enter for over 400s (**Fig. 2c**). Once entered, C57 mice moved along the chamber and spent on average 15% of time in the light chamber during the session (**Fig. 2a, e**). All DBA mice entered the light chamber with a relatively long latency (mean ± SEM: 375 ± 43 s) and often quickly retreated to the dark chamber without exploring the entire light chamber (**Fig. 2a, c**). 2/11 BC mice never entered the light chamber (**Fig. 2b**). For animals that entered, the latency was surprisingly short (<60s), and some animals spent over 80% of their time in the light chamber (**Fig. 2c, e**). However, a closer look at the behavior revealed that BC mice often stayed along the light chamber wall with little movement, likely reflecting a high anxiety level (**Fig. 2a, f**). Only 2/12 129 mice ever entered the light chamber, suggesting their high anxiety phenotype^42^ (**Fig. 2b**). Most 129 mice spent the entire time in one corner of the dark chamber (**Fig. 2a, b, d, e**).

The animals’ performance in the OFT is generally consistent with theirs in LDB (**Fig. 2m-r**). CD1, SW, and FVB mice moved in the arena extensively, repeatedly passing the center zone (**Fig. 2m-r**). In comparison, C57 mice traveled a bit less but also entered the center zone many times (**Fig. 2o**). DBA and BC moved along the edges of the arena but rarely entered the center zone, while 129 mice spent most time hugging the wall (**Fig. 2m-r**). Overall, DBA, BC, and 129 mice spent significantly less time in the center zone compared to other strains, suggesting their higher anxiety level (**Fig. 2p**).

For 5 strains, including CD1, SW, C57, BC and 129, we also assessed their anxiety level using elevated plus maze (EPM) (**Supplementary Fig. 4**). Consistent with the LDB and OFT results, CD1 and SW mice showed the lowest anxiety level: all animals entered the open arms almost immediately and repeatedly, and all but one animal walked all the way to the edge of the open arm (**Supplementary Fig. 4a-g)**. Similarly, all C57 males entered the open arm, although with a slightly longer latency than CD1, and spent on average approximately one minute in the open arm (**Supplementary Fig. 4a-g)**. While 9/10 BC entered the open arm, only 3/10 reached the edge, and the entry latency tends to be longer (**Supplementary Fig. 4a-g)**. Lastly, for 129 mice, 4/10 entered the open arm and only one walked to the far end (**Supplementary Fig. 4a-g)**.

The strain difference in anxiety was also observed in virgin female mice (**Supplementary Fig. 3g-x**). Compared to C57 females, SW females exhibited more frequent and prolonged visits to the light compartment in the LDB, as well as increased entries into and time spent in the open arms of the EPM (**Supplementary Fig. 3g-r**). In contrast, performance in the OFT, including total distance traveled, center entries, and percentage of time in center did not differ significantly between SW and C57 females (**Supplementary Fig. 3s-x**). This pattern mirrors observations in males, where SW and C57 differ in LDB and EPM but not OFT (**Fig. 2 and Supplementary Fig. 4**). This likely reflects the lower sensitivity of the OFT as an anxiety assay, as its readouts are strongly influenced by locomotor and exploratory drives^43^.

Anxiety has been shown to influence aggression, although the direction of influence remains controversial^44–48^. We next examined the relationship between anxiety- and aggression-related parameters across strains and found a clear relationship (**Fig. 2g, j, s, v**). For example, the latency to attack an intruder is significantly positively correlated with the latency to enter the light box: the faster the animal attacks, the sooner it enters the light box, i.e., less anxious (**Fig. 2g**). Similarly, the attack duration is significantly positively correlated with percentage of time in center (**Fig. 2v).** Indeed, across LDB, OFT, and EPM, anxiety-related parameters are significantly correlated with male aggression-related parameters consistently (**Fig. 2g, j, s, v, Supplementary Fig. 4h, k**). Although the limited number of female strains examined in the study precludes correlation analysis, the more aggressive SW females showed lower anxiety than less aggressive C57 females, supporting an inverse relationship between anxiety and aggression in both sexes (**Supplementary Fig. 3y-dd**).

The levels of anxiety and aggression also varied among individuals within a strain, likely reflecting individual-specific life experiences. We therefore asked whether these two traits remain correlated among animals with the same genetic background. Correlation analyses of anxiety-and aggression-related parameters within each of the seven strains failed to reveal a consistent relationship (**Supplementary Figs. 5–7**). Although a statistically significant correlation between certain pairs of parameters in certain strains was detected in a couple of cases, e.g., latency to attack and latency to enter the light box of CD1 mice, the correlation could be either positive or negative (**Supplementary Figs. 5a1, 5a3, 5a4, 5b2, 5f4, 5h4, 6a4, 6b3, 6c3, 6g4, 6h2, 7b1**).

To further assess the relationship between anxiety and aggression independently of genetic background, we z-scored each animal’s aggression- and anxiety-related parameters relative to the mean and standard deviation of its own strain. These normalized values, therefore, reflect each animal’s relative aggression and anxiety levels compared to others with the same genetic background. We then examined the relationship between strain-normalized aggression and anxiety across individuals from all seven strains and found no significant correlation for any pair of parameters (**Fig. 2h, k, t, w**). Consistent with this, when animals were divided into relatively high- and low-aggression groups based on whether their aggression level was above or below the strain mean, the two groups did not differ in strain-normalized anxiety levels (**Fig. 2i, l, u, x**). These results suggest that anxiety and aggression are influenced by overlapping genetic factors, whereas the within-strain variation captured in our study, whether experiential or genetic (e.g. epigenetic or spontaneous mutation), is not sufficient to produce a consistent correlation between the two traits.

Because performance in anxiety assays is largely based on movement, we next sought to separate the contributions of anxiety-related behavior and general locomotion to strain differences in aggression. We examined animal’s locomotion in its home cage, which reflects animal’s spontanous mobility in a low-anxiety state (**Supplementary Fig. 8a**). Although animals of some strains (e.g. DBA and CD1) showed higher spontaneous homecage locomotion than other strains (e.g. 129 and FVB), there is no significant correlation between home cage locomotion and aggression level (**Supplementary Fig. 8b-h**). This contrasts with the significant correlation between aggression and locomotion-dependent anxiety measures, such as distance travelled in the OFT and the number of light-box entries in the LDB, suggesting that these correlations do not simply reflect strain differences in locomotion (**Fig. 2j, s**). Furthermore, in a combined regression model (Aggression = β₀ + β₁Anxiety + β₂Mobility + ε), when anxiety was quantified as percentage of time in OFT center and mobility as home-cage locomotion, the anxiety measure remained a significant predictor of aggression (β₁ = 0.875, p = 0.008) after controlling for mobility. In contrast, locomotor activity did not (β₂ = 0.142, p = 0.476). A similar pattern was observed when anxiety was quantified as light-box entry number in the LDB (β₁ = 0.865, p = 0.038; β₂ = 0.031, p = 0.917). The relationship between anxiety and aggression independent of mobility can be visualized by plotting the residual aggression after regressing out home-cage locomotion against anxiety-related measures. In this analysis, residual aggression remained significantly correlated with OFT percentage of time in center and LDB light-box entry number (**Supplementary Fig. 8i, j**). These results suggest that the association between anxiety and aggression cannot be explained simply by differences in general locomotor activity across strains.

To functionally investigate the link between anxiety and aggression, we i.p. injected Yohimbine (YO), an α2-adrenergic receptor antagonist and a potent anxiogenic agent^49^, into CD1 mice to examine its potential influence on aggression (**Supplementary Fig. 9a**). Consistent with the anxiogenic effect of YO, YO-injected CD1 mice significantly altered their behaviors in the LDB (**Supplementary Fig. 9b**). All mice spent significantly less time in the light chamber after YO injection compared to saline injection (**Supplementary Fig. 9c-f**). Importantly, the maximal movement velocity did not change, suggesting locomotion was not compromised (**Supplementary Fig. 9g**). Strikingly, aggression was nearly abolished after YO injection (**Supplementary Fig. 9h-m**). While saline-injected CD1 residents attacked the BC intruders quickly and repeatedly, only 3/5 males attacked the intruder briefly (<2s) after YO injection (**Supplementary Fig. 9i-s)**. Together, these results suggest anxiety and aggression are tightly and inversely linked to each other.

### Relationship between aggression and corticosterone

Corticosterone (CORT) is widely known as the “stress hormone” as it is released under stressful situations through activation of the hypothalamic-pituitary-adrenal (HPA) axis. Given the tight correlation between baseline anxiety and aggression, we asked whether CORT is predictive of aggression level. To measure CORT, we collected urine samples of unstressed mice one week before the RI test and performed ELISA (**Fig. 1b**). We found that CORT levels differed significantly across strains: FVB and DBA mice had the lowest CORT levels; BC mice had the highest level; CD1, SW, C57, and 129 mice had intermediate levels (**Fig. 3a**). However, these differences in CORT did not align with the observed strain differences in aggression. We found no correlation between CORT level and attack latency or attack duration across strains (**Fig. 3b, e**). Indeed, DBA, 129 and BC, the three low-aggression strains, showed the second lowest, intermediate and highest CORT levels (**Fig. 3a**). Furthermore, no strain showed a significant correlation between individual CORT levels and aggression (**Supplementary Fig. 10a1-g1, a2-g2**). When combining all the strains, we did not find a correlation between strain-normalized CORT level and aggression level (**Fig. 3c, f**). In other words, the relative CORT level of an animal among all animals of the same strain does not predict the relative aggression level of the animal within the strain. High-aggression and low-aggression individuals within strains had comparable CORT levels **(Fig. 3d, g)**.

Given that anxiety and aggression are tightly linked, the lack of correlation between CORT and aggression suggests that baseline CORT may not be a good indicator of anxiety levels in unstressed animals. Consistent with this prediction, we found no significant correlation between CORT levels and LBD or OFT performance across strains or individuals (**Supplementary Fig. 11**).

### Relationship between aggression and Major Urinary Protein

Major urinary proteins (MUPs) are a family of proteins secreted by the liver. They bind pheromones and are important for individual recognition and social communication^50^. In our recent study focusing on C57 mice, we found that the MUP level is higher in future aggressive mice than in non-aggressive mice^15^. Other studies have shown that the MUP level is higher in dominant than in subordinate CD1 mice^50^. Thus, we asked whether MUP concentration can predict the aggression level of animals across strains and individuals. As MUP constitutes 90% of all urinary proteins^51^, we measured the total urinary protein level normalized by the creatinine level as an approximation of MUP concentration. As in the case of CORT, urinary protein was measured in urine collected from naïve animals one week before RI tests (**Fig. 1b**).

MUP levels varied significantly among strains: DBA and CD1 show the low MUP concentration, whereas FVB, BC and 129 mice show high levels (**Fig. 3h**). Across strains, MUP level and attack duration or latency were uncorrelated (**Fig. 3i, l**). However, correlational analysis between MUP and aggression across individuals of the same strain revealed a significant positive relationship or a trend in this direction in the majority of strains (**Supplementary Fig. 10a3-g3, a4-g4**). When combining all strains, we found a positive correlation between strain-normalized aggression and MUP levels across individuals (**Fig. 3j, m**). High-aggression (> strain average) animals showed higher MUP levels than low-aggression (< strain average) animals of the same strain (**Fig. 3k, n**). Thus, while the MUP level in each strain is likely controlled by strain-specific metabolism unrelated to aggression, the MUP level is predictive of future aggression levels in naïve animals with the same genetic background.

### Differential activation of the aggression circuit across strains

Aggression is generated and modulated by several interconnected nuclei^1,52^, including posterodorsal medial amygdala (MeApd)^53,54^, posteroventral MeA (MeApv)^55^, principal nucleus of the bed nucleus of the stria terminalis (BNSTpr)^56,57^, posterior amygdala (PA)^58,59^, posterolateral cortical amygdala (CoApl)^60^, dorsal and ventral parts of lateral septum (LSd and LSv)^61–64^, medial preoptic area (MPOA)^65^, anterior hypothalamus (AHN)^66,67^, ventrolateral part of the ventromedial hypothalamus (VMHvl)^35,68,69^, PMv^70–72^, and periaqueductal gray (PAG)^73^. The difference in aggressive behaviors across strains is presumably due to the differential responses of the aggression circuit to the male intruder. Based on this rationale, we evaluated the responses of the aggression circuit to a cupped male mouse in low (129), middle (C57), and high (SW) aggression strains using c-Fos, an immediate early gene across the limbic system (**Supplementary Table 2**). We used a cupped animal as the stimulus to minimize variability arising from differences in aggressive interactions and the rewarding effects of winning^32^. In control C57 mice, we introduced an empty cup into their home cage (**Fig. 4a**).

**Figure 4.**
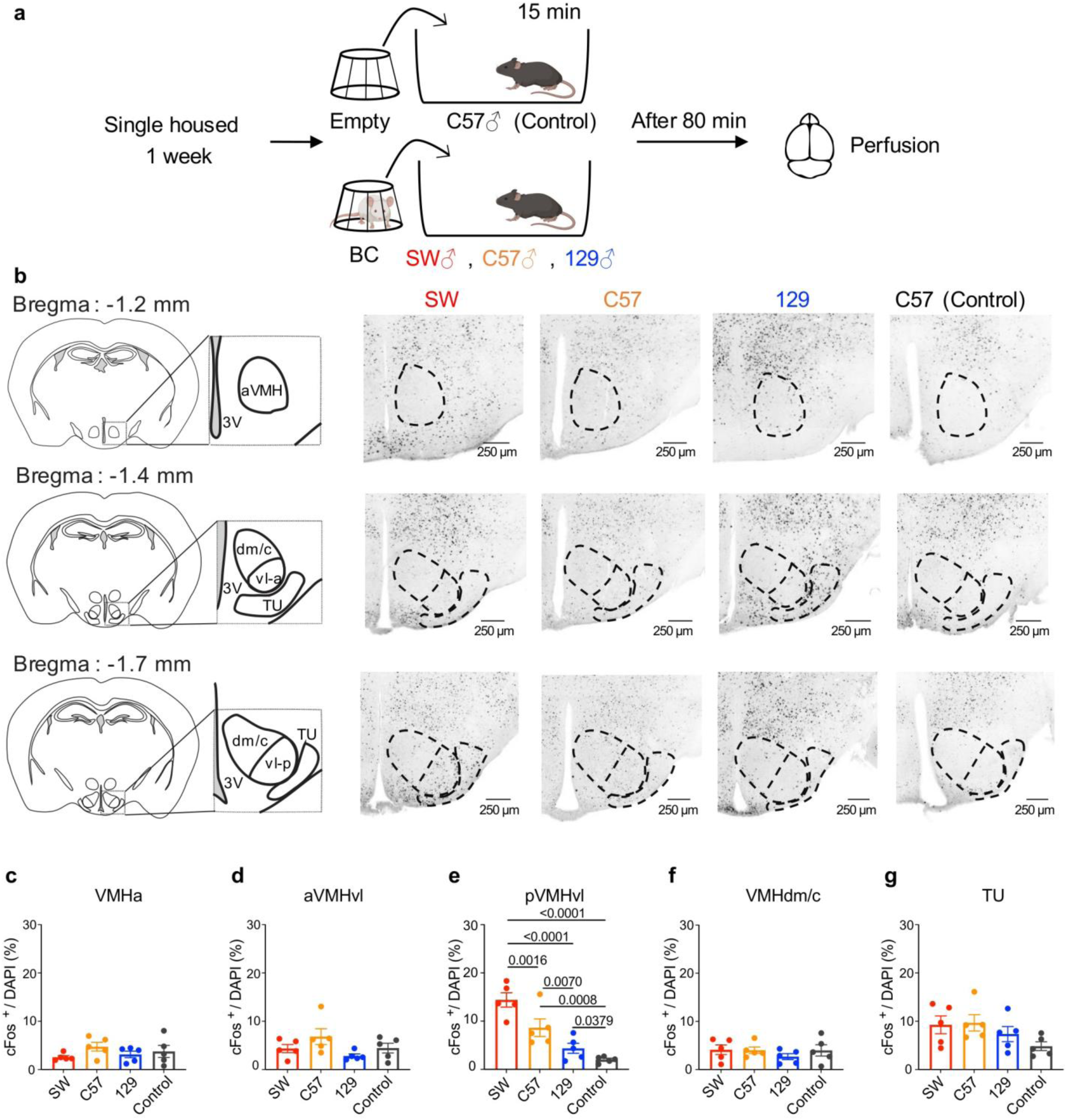
Male pVMHvl responses to male intruders vary across strains. **(a)** Experimental timeline for c-Fos induction. Male mice were single-housed for one week, exposed to a 15-minute cupped male intruder or an empty cup, and perfused 80 minutes later. **(b)** Left: mouse brain atlas showing different VMH subdivisions and tuberal nucleus (TU). Right: coronal brain sections showing intruder-induced c-Fos expression at the Bregma levels -1.2, -1.4 and -1.7mm of representative SW, C57, and 129 male mice and empty cup-induced c-Fos expression of a C57 male mouse. c-Fos positive cells appear black. **(c–g)** Quantification of c-Fos positive cells in **(c)** anterior VMH (aVMH), **(d)** anterior VMHvl (aVMHvl), **(e)** posterior VMHvl (pVMHvl), **(f)** dorsomedial/central VMH (VMHdm/c), and **(g)** TU. n = 5 male mice per group. Circles represent data of individual animals. Bar and error bars represent mean ± SEM. One-way ANOVA for normally distributed datasets or Kruskal–Wallis test for non-normally distributed datasets with FDR-corrected post hoc comparisons (Benjamini–Krieger–Yekutieli method). All statistical tests are two-tailed. Exact p- or q-value is shown if ≤ 0.05. Otherwise, p- or q-value is unspecified. See **Supplementary Table 1** for additional statistical details.

VMHvl showed the largest inter-strain differences in c-Fos expression (**Fig. 4b**). Specifically, posterior VMHvl (pVMHvl) expressed c-Fos differentially across strains: highest in SW mice, followed by C57 mice, and the lowest in 129 mice. In contrast, c-Fos level in the anterior VMHvl (aVMHvl) was similar across strains and unelevated in comparison to control mice (**Fig. 4b, d, e**). This is consistent with previous studies showing that aVMHvl is critical for social defense, while the pVMHvl is for aggression^31,74^. Tuberal nucleus (TU), a region lateral to pVMHvl, provides GABAergic input to the pVMHvl cells^75^. A trend of increase in TU c-Fos was observed in all strains with no clear strain difference (**Fig. 4g**). Other VMHvl neighboring regions, including the anterior VMH (aVMH) and the dorsomedial and central parts of the VMH (VMHdm/c), showed no increase in c-Fos expression regardless of the strain (**Fig. 4c and f**).

Beyond the VMHvl, c-Fos also showed a significant increase in the MeApd and PMv after exposure to a cupped male mouse, although there is no strain difference (**Supplementary Fig. 12a6-c6, a8-c8**). None of the other analyzed regions showed a significant increase in c-Fos expression after intruder exposure compared with control animals (**Supplementary Fig. 12**). DAPI quantification showed no significant difference in the number of cells across strains in the analyzed regions (**Supplementary Fig. 13**).

We further asked whether the VMHvl contributes to strain-dependent differences in aggression in virgin females by examining VMHvl c-Fos expression after SW and C57 females were exposed to a cupped juvenile male mouse for 15 minutes (**Supplementary Fig. 14a**). For analysis, we subdivided the VMHvl into medial (VMHvlm) and lateral (VMHvll) regions, as VMHvlm is preferentially implicated in female aggression^36,37^. In contrast to males, however, we did not observe significant differences in c-Fos expression level in the posterior VMHvlm between SW and C57 females (**Supplementary Fig. 14b, f)**. Notably, both strains exhibited low levels of c-Fos expression following exposure to cupped juveniles, comparable to male 129 mice and substantially lower than male C57 and SW mice (**Fig. 3e, Supplementary Fig. 14f**). Other VMHvl regions also did not show a significant difference in c-Fos expression between SW and C57 (**Supplementary Fig. 14c, d, e, g, h**). These results suggest that differences in VMHvl activity may contribute more to aggression variability across strains in males than in females.

### Physiological properties of VMHvl cells differ across strains

We wonder whether the strain differences in pVMHvl cell responses to male intruders can be explained by the physiological properties of the cells. To test this possibility, we performed whole-cell patch-clamp recordings of pVMHvl cells in naïve, single-housed male mice of the SW, C57, and 129 strains. To identify the pVMHvl on the brain slice, we injected red retrobeads into the MPOA, a region known to receive dense projections from the pVMHvl^74,76^ (**Fig. 5a**). One week after retrobeads injection, we recorded cells in the pVMHvl, identified based on both anatomical landmarks and fluorescent labeling (**Fig.5b**).

**Figure 5.**
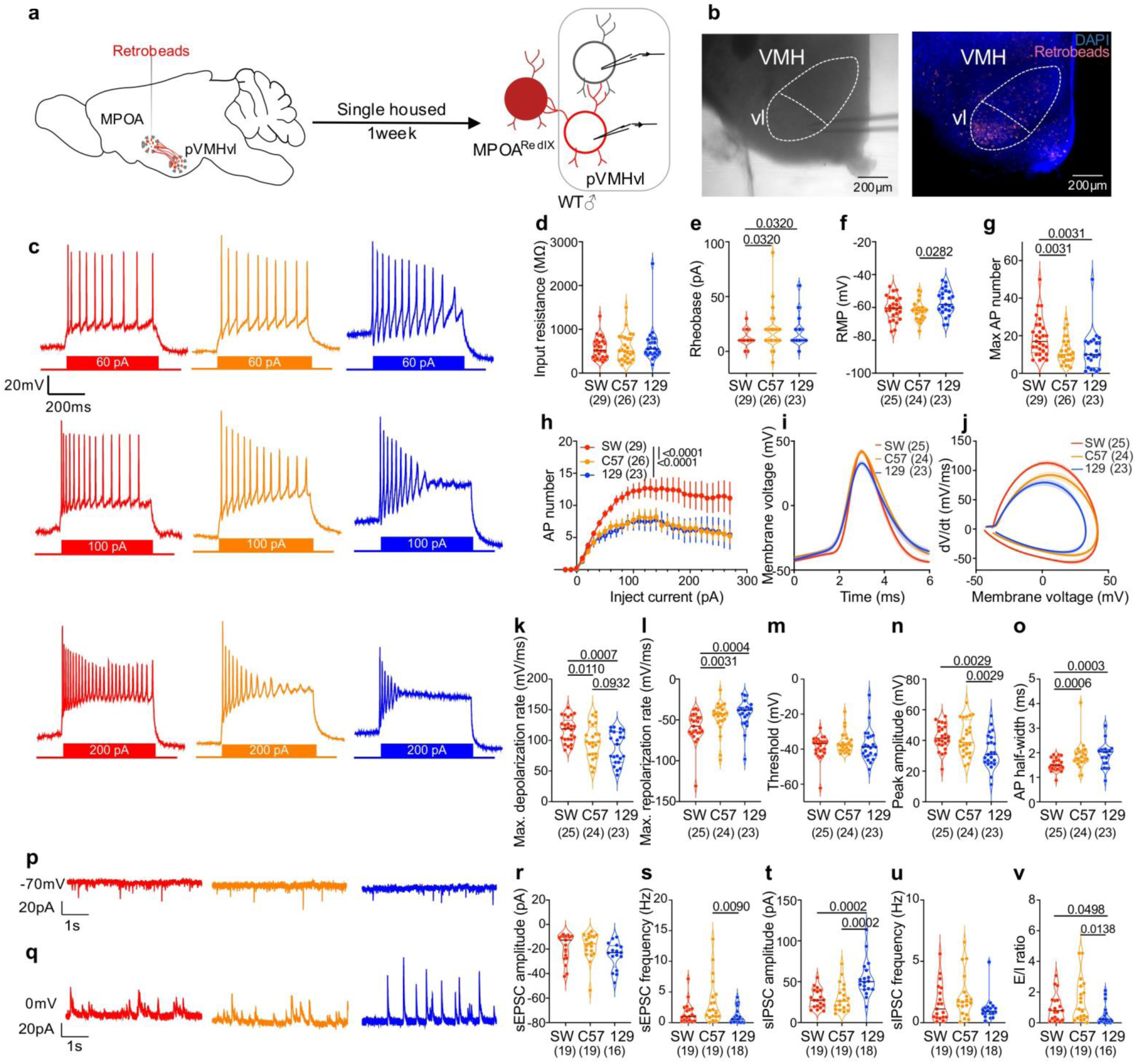
Physiological properties of VMHvl cells differ across strains in male mice. **(a)** Experimental design for patch-clamp recordings. Retrobeads were injected into the medial preoptic area (MPOA) to label neurons projecting to the posterior ventrolateral ventromedial hypothalamus (pVMHvl). Following post-surgical recovery and one week of single housing, brain slices were prepared, and both retrobead-positive and - negative cells in the area containing dense retrobead labeling were recorded. **(b)** Representative image showing the patched cell location under Differential Interference Contrast (DIC) microscopy (left) and fluorescence microscope (right). **(c)** Representative recording traces from pVMHvl cells in SW (red), C57 (orange), and 129 (blue) mice in response to 60, 100, and 200 pA current injections. **(d-f)** Intrinsic properties of pVMHvl cells across strains, including **(d)** input resistance, **(e)** rheobase and **(f)** resting membrane potential (RMP). **(g)** Maximum action potential (AP) number across SW, C57, and 129 pVMHvl cells. **(h)** Spike counts across –20 pA to 270 pA current steps, each for 500 ms. **(i)** Average waveform of the first action potential across different current steps in pVMHvl neurons from SW, C57, and 129 male mice. **(j)** Phase plots (dV/dt vs. V) for first spikes across different current steps in pVMHvl neurons from SW, C57, and 129 male mice. **(k-o)** First spike properties, including **(k)** maximum depolarization rate, **(l)** maximum repolarization rate, **(m)** spike threshold, **(n)** spike peak amplitude, and **(o)** action potential half-width. **(p-q)** Representative traces of **(p)** sEPSCs and **(q)** sIPSCs. **(r-v)** Quantification of synaptic activities, including **(r)** sEPSC amplitude, **(s)** sEPSC frequency, **(t)** sIPSC amplitude, **(u)** sIPSC frequency, and **(v)** excitation/inhibition (E/I) ratio. Numbers in parentheses indicate cell numbers. Cells in **(d-o)** were from 5 SW, 5 C57, and 6 129 mice, while cells in **(r-v)** were from 6 SW, 6 C57, and 6 129 mice. Circles represent individual cells. Solid lines and shades in **(i, j)** represent mean ± SEM. Error bars in **(h)** represent ± SEM. Solid line in (**d-g, k-o, r-v**) represents the median for each group, while dashed lines indicate quartiles. **(d-g, k-o, r-v**) One-way ANOVA for normally distributed datasets or Kruskal–Wallis test for non-normally distributed datasets with FDR-corrected post hoc comparisons (Benjamini–Krieger–Yekutieli method); **(h)** Two-way ANOVA with FDR correction. All statistical tests are two-tailed. Exact p- or q-value is shown if ≤ 0.05. Otherwise, p- or q-value is unspecified. See **Supplementary Table 1** for additional statistical details.

pVMHvl cells in SW, C57, and 129 mice did not differ in their input resistance, although cells in 129 showed more depolarized resting membrane potential (RMP) and cells in SW mice showed lower rheobase (**Fig. 5d-f**). Frequency-current (F-I) curves revealed significant differences in the intrinsic excitability of cells across mouse strains (**Fig. 5g, h**). SW pVMHvl cells could reach a significantly higher maximum firing rate in comparison to cells in C57 and 129, while the maximum firing rate of pVMHvl cells in C57 and 129 mice was comparable (**Fig. 5g, h**). We further examined the spike waveforms using phase plots (**Fig. 5i, j**), and found that the maximum depolarization rate of pVMHvl cells differed significantly across strains: the highest in SW, moderate in C57, and the lowest in 129, suggesting potential sodium current differences (**Fig. 5k**). We also found the maximum repolarization rate was higher in SW than C57 and 129, suggesting different potassium current levels (**Fig. 5l**). The threshold voltage to fire action potentials did not differ across strains, while the peak voltage of action potentials was lower in 129 mice than in C57 and SW mice (**Fig. 5m, n**). The half-width of spikes was narrower in SW mice than in C57 and 129 mice, consistent with the higher firing rates of cells in SW mice (**Fig. 5o**).

We next compared spontaneous excitatory and inhibitory postsynaptic currents (sEPSCs and sIPSCs) in pVMHvl cells across strains using voltage-clamp recordings (**Fig. 5p, q**). Interestingly, we observed that sIPSC amplitudes were higher in 129 mice than in SW and C57 mice, whereas sEPSC frequencies in 129 mice tended to be lower (**Fig. 5r-v**). 2/18 pVMHvl cells in 129 mice received no detectable spontaneous excitatory inputs during the recorded period (40s in total), while no cells in C57 and SW mice exhibited the same phenotype (**Fig. 5r, s**). Overall, the excitation–inhibition (E/I) ratio of pVMHvl cells in 129 mice was significantly lower than that in C57 and SW mice (**Fig. 5v**). These results suggest that pVMHvl cells in 129 mice receive lower excitatory drive and higher tonic inhibition than C57 and SW mice, consistent with overall dampened responses of pVMHvl cells to aggression-provoking cues and minimal aggressive behaviors in 129 males.

We further compared the intrinsic and synaptic properties of VMHvlm neurons between SW and C57 females (**Supplementary Fig. 15**). To label VMHvlm neurons, we injected CTB-Alexa 488 retrograde tracer into the MPOA, which labels both pVMHvlm and pVMHvll populations, and red retrobeads into the anteroventral periventricular nucleus (AVPV), which preferentially labels pVMHvlm neurons^37,77^ (**Supplementary Fig. 15a**). During patch-clamp recordings, we targeted both CTB-Alexa 488 labeled and unlabeled cells within the pVMHvlm region (green positive and red negative) (**Supplementary Fig. 15a, b**). Consistent with the c-Fos results, we observed no significant differences in intrinsic properties of the cells between SW and C57 pVMHvlm neurons, including input resistance, resting membrane potential, rheobase, F–I relationships, and maximal firing rates (**Supplementary Fig. 15c, e-i**). Interestingly, however, spike waveform differed between strains: C57 neurons exhibited faster depolarization and repolarization rate and consequently narrower spike widths, while the peak spike amplitude and spiking threshold were similar between the two strains (**Supplementary Fig. 15j-p**). Because spike width can influence presynaptic Ca²⁺ entry, the difference in spike width suggests that pVMHvlm neurons in SW females may exert stronger downstream synaptic drive despite similar firing rates. We also examined synaptic inputs onto pVMHvlm neurons using voltage-clamp recording and detected no significant differences in sEPSC or sIPSC frequency or amplitude, as well as E/I ratio between SW and C57 females (**Supplementary Fig. 15d, q-u**).

When comparing between sexes, pVMHvlm neurons in females exhibited lower intrinsic excitability than males, particularly in SW mice, in which male pVMHvl neurons are highly excitable (**Supplementary Fig. 16a-j, p-y**). With regard to synaptic inputs, regardless of strains, female pVMHvlm cells generally received lower-amplitude sEPSC and sIPSC than male pVMHvl cells and the E/I ratio also tended to be lower in females (**Supplementary Fig. 16k-o, z-dd**). Together, these findings suggest that strain- and sex-dependent tuning of pVMHvl excitability and synaptic inputs could contribute to variability in aggressive behaviors across individuals.

### Chemogenetical activation of pVMHvl cells increases aggression in low-aggression strain

To address whether the difference in pVMHvl cell excitability is causally linked to the different aggression levels across strains, we chemogenetically activated pVMHvl neurons by injecting a retrograde virus (AAVrg-Cre) into the MPOA and a Cre-dependent AAV expressing hM3Dq-mCherry into the VMHvl of 129 male mice (**Fig. 6a, b**). Notably, activation of hM3Dq engages Gq signaling, leading to PLC activation, increased intracellular Ca²⁺, and suppression of K⁺ conductance (e.g., M-current), thereby enhancing neuronal excitability and approximating the physiological properties of pVMHvl neurons observed in more aggressive strains^78^. Histology analysis showed that 86.6% of retrogradely labeled mCherry+ cells express Esr1, a molecular marker of the aggression-relevant population in the VMHvl (**Fig. 6c, d**)^68^.

**Figure 6.**
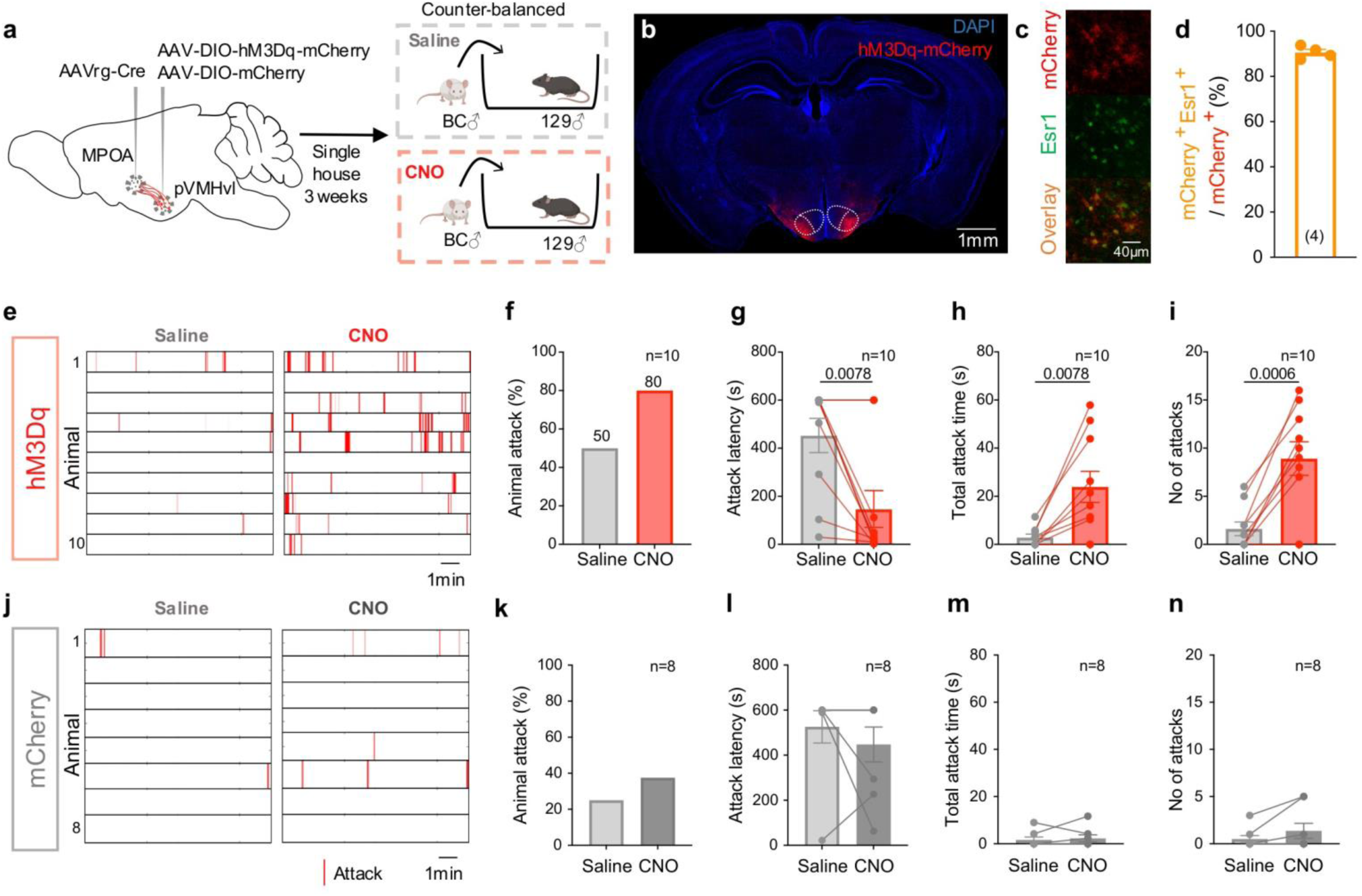
Chemogenetic activation of MPOA-projecting pVMHvl neurons promotes aggression in 129 male mice. **(a)** Experimental design. AAVrg-Cre was injected into the medial preoptic area (MPOA) and AAV-DIO-hM3Dq-mCherry (or AAV-DIO-mCherry control) was injected into the pVMHvl of 129 male mice. After 3 weeks of recovery, mice were single-housed and tested in a resident–intruder assay after saline or CNO injections in a counterbalanced order. **(b)** A representative image showing hM3Dq-mCherry expression in the pVMHvl. **(c)** An images showing retrogradely labeled mCherry^+^ cells (red) and estrogen receptor alpha (Esr1) expression (green) in pVMHvl. **(d)** Quantification of mCherry⁺ cells that co-express Esr1 in the pVMHvl. **(e)** Raster plots showing attack events under saline and CNO conditions of all hM3Dq-expressing mice. **(f–i)** Quantification of aggression in hM3Dq mice, including **(f)** percentage of animals that attacked, **(g)** attack latency, **(h)** total attack duration, and **(i)** attack frequency. n = 10 animals. Lines connect paired measurements. **(j)** Raster plots showing attack events under saline and CNO conditions of all mCherry control mice. **(k–n)** Quantification of aggression in control mice, including **(k)** percentage of animals that attacked, **(l)** attack latency, **(m)** total attack duration, and **(n)** attack frequency. n = 8 animals. Lines connect paired measurements. Circles represent data of individual animals. **(f-i, k-n)** Bar and error bars represent mean ± SEM. **(g-i, l-n)** Paired t test for normally distributed datasets or Wilcoxon test for non-normally distributed datasets. All statistical tests are two-tailed. Exact p is shown if ≤ 0.05. Otherwise, p value is unspecified. See **Supplementary Table 1** for additional statistical details.

Three weeks after virus injection, we i.p. injected saline and CNO on separate days in a counter-balanced order. To avoid the winner effect^15^, we waited for over a week for the second RI test if the animal attacked the intruder during the first test. We observed a significant increase in aggression in 129 mice following CNO administration compared to saline injection (**Fig. 6e-i**). After saline injection, 5/10 mice attacked a BC male intruder, whereas 8/10 mice did so following CNO injection(**Fig. 6e, f**). CNO-treated mice also exhibited significantly shorter attack latency, more frequent attacks, and longer total attack duration (**Fig. 6g-i**). In contrast, no differences in aggression were observed in mCherry control animals following saline or CNO injection (**Fig. 6j-n**). These results, along with previous chemogenetic activation experiments in C57 mice^79^, demonstrate that the excitability of pVMHvl neurons contributes causally to the aggression level of the animals.

## Discussion

Here, we demonstrated that the aggression levels of naïve male mice differ widely depending on the animals’ genetic background. The strain-specific aggression level aligns with the strain-specific pVMHvl cell responses to aggression-provoking cues, which can be attributed to the cell’s intrinsic properties and synaptic inputs. Furthermore, we found an inverse relationship between innate aggression and baseline anxiety level across strains and a positive correlation between aggression and urinary protein levels across individuals of the same genetic background.

### Neural mechanisms determining the innate aggressiveness

In 1942, two studies examined inter-male aggression across three mouse strains, including C57, C3H, and C albino, demonstrating for the first time the potential genetic influences on aggressive behaviors^29,30^. Since then, multiple rodent studies using over 10 different strains confirmed that an animal’s aggressiveness is tightly linked to its genetic background^25–28,80^. Our behavioral results are broadly consistent with these earlier studies, showing that certain mouse strains, such as CD1, are highly aggressive^80^, whereas others, such as DBA, exhibit minimal aggression^26^, suggesting that strain-dependent differences in baseline aggression are reproducible across cohorts and across studies. One exception is BALB/c strain, which reportedly showed high aggression in earlier studies^30,81^, while we found that the aggression level of BALB/c male mice is low. This discrepancy may be due to the significant differences in aggression levels across BALB/c sub-strains^82,83^. Our BALB/c mice, purchased from Charles River, have been bred separately from those available from the Jackson Laboratory since 1932.

It is worth noting that although CD1 and SW are outbred mice, and the other five strains are inbred, we did not find higher individual variability in CD1 and SW mice than in the inbred strains. This is consistent with a previous study comparing various social behaviors between CD1 and C57BL/6J^80^. The low variability across individuals in outbred strains likely reflects the fact that these strains were established from a small number of founder mice captured in the same region. These founder mice likely had relatively similar aggression levels, as aggression is under the pressure of sexual selection, which is strongly influenced by resource availability in the habitat^84^.

After recognizing a clear link between aggression and genetics decades ago, follow-up research focused mainly on identifying genes that can influence aggression. By comparing the genomic landscapes of high- and low-aggression strains, several quantitative trait loci (QTL) that may affect aggressiveness have been reported^26,27,81^, although pinpointing the genes causally linked to aggression has been challenging. Meanwhile, numerous knockout studies revealed dozens of genes that could alter aggression^85^, including MAOA, the famous “warrior gene”^86^. These genetic studies clearly demonstrate that aggression is a complex behavior modulated by many genetic factors.

Genes must modulate behaviors through the neural circuit driving the behavior. How the genetic variability across strains translates to the neural circuit difference to influence aggression has been largely unexplored. Here, our c-Fos mapping revealed that responses of the pVMHvl to an intruder, but not responses of any other analyzed brain regions, correlate with the strain’s aggression level. pVMHvl is a key node for inter-male aggression^34,35^. In male mice, pVMHvl is activated by aggression-provoking cues, e.g., pheromones from male conspecifics, and is the first to increase activity among aggression-related brain regions^87^. When pVMHvl is artificially activated, it can drive the animal to attack within seconds^35^. Thus, pVMHvl is a critical site for detecting aggression-provoking social cues and authorizing attacks. As the pVMHvl responses to aggression-provoking cues increase, the probability of attack initiation increases.

What are the underlying mechanisms that cause the pVMHvl cells to respond differently across strains? The answers appear to lie in both cellular and synaptic variations. Between C57 and SW animals, we found that the excitability of pVMHvl cells in SW males is significantly higher than that in C57, explaining the higher intruder responses in SW than in C57 mice. Between C57 and 129, the pVMHvl cell excitability does not differ. However, pVMHvl cells in 129 mice receive less frequent excitatory synaptic inputs and stronger inhibitory inputs, likely contributing to the muted responses of 129 mice to intruder cues.

Intriguingly, our recent study revealed that pVMHvl cells in C57 male mice increase excitability and responses to male intruders after repeated winning (≥ 5 days)^15^. We speculate that the genetic background of the animal could determine the initial set point of the aggression circuit. In strains of mice with a low starting point, the winning experience could enhance the aggression circuit by making it more “high aggression strain” like. These data suggest that genetic and experiential factors may utilize the same synaptic and cellular mechanisms to modify the input-output relationship of the behavioral circuit, thereby altering the behavioral output. The convergence of “nature” and “nurture” influences at the neural level is likely not specific to the aggression circuit. In our recent study, we found that the different tendencies of C57 and SW virgin female mice in expressing infanticide can be attributed to the difference in the excitability of BNSTp^Esr1^ cells, a key population that drives pup-attack^88^. During motherhood, BNSTp^Esr1^ cell excitability in SW females decreases to suppress infanticide and enable maternal care^88^.

### Relationship between anxiety and aggression

In our study, we identified an inverse relationship between innate aggression and anxiety in naïve male mice. High-aggression strains, such as CD1 and SW, exhibited the lowest anxiety-like behavior in the open field and light–dark box tests, whereas low-aggression strains, such as BALB/c, 129, and DBA, exhibited the highest anxiety. This finding is consistent with our recent study showing that repeated winning increases aggression while reducing anxiety-like behavior, as measured in the light–dark box and novelty-suppressed feeding assays^15^. Our results in females further support an inverse relationship between anxiety and aggression: naïve SW females, which are more aggressive than C57 females, also displayed lower anxiety. Previous studies similarly showed that during lactation, female aggression increases while anxiety decreases^89^.

Our findings are also consistent with several earlier studies in high- and low-aggression strains of mice^44,45^ and rats^46^. For example, the Turku aggressive (TA) and non-aggressive (TNA) lines are SW-derived mouse lines selectively bred in Turku, Finland, for high and low levels of aggression^90^. In multiple anxiety-related tests, aggressive TA mice were found to be less anxious and more active than the non-aggressive TNA line^45^. However, other studies have reported that aggression and anxiety are positively correlated^47^ or uncorrelated^48^. For example, NC900 and NC100 are ICR-based mouse lines, closely related to CD1, that were selectively bred in North Carolina for high and low aggression^91^. In both the OF and LDB tests, the high-aggression NC900 line displayed higher anxiety than the low-aggression NC100 line^92^. In another study, rats were selectively bred for extremely high- or low-anxiety phenotypes, and aggression changed concurrently despite not being under direct selection. Intriguingly, both the high-anxiety and low-anxiety lines were more aggressive than unselected controls^93^. Together, these findings indicate that anxiety and aggression are closely linked, although the nature of their relationship is complex and may depend on genetic background and selection history.

What mechanisms underlie the co-variation between anxiety and aggression? Because these traits are correlated across strains, they are likely influenced, at least in part, by shared genetic factors. Indeed, both aggression and anxiety are regulated by monoaminergic signaling, including dopamine, norepinephrine, and serotonin. Genes that affect these systems, such as Drd2, Maoa, Comt, and Slc6a4 (5-HTT), have been shown to alter both anxiety-like and aggressive behaviors^94,95^. Thus, neural circuits controlling aggression and anxiety may be shaped by common neuromodulatory systems. Consistent with this idea, we found that pharmacologically increasing anxiety by antagonizing α2-adrenergic receptors virtually abolished aggression in highly aggressive SW mice.

Additionally, anxiety-promoting regions may directly and tonically suppress the aggression circuit. Several brain regions implicated in anxiety and fear, including the LSv and AHN, are predominantly GABAergic^96–98^ and send dense projections to key aggression-promoting nodes such as the VMHvl^76^. Inhibiting LS elevates aggression, while optogenetic activation of the LS to VMHvl pathway is sufficient to terminate attack acutely^76^. In high-anxiety animals, elevated baseline activity in these regions may impose sustained inhibitory tone onto aggression-related circuits. Consistent with this possibility, VMHvl neurons in 129 male mice exhibit higher sIPSC amplitudes than those in SW and C57 males.

It is worth noting that the observed difference in VMHvl excitability between SW and C57 mice is unlikely to account for the differences in anxiety between the two strains. In our recent study, experimentally increasing VMHvl excitability robustly elevated aggression without altering anxiety-like behavior^15^.

Taken together, these findings suggest that aggression and anxiety covary through at least two mechanisms: (1) shared genetic influences—particularly those regulating neuromodulatory systems—that shape the function of both anxiety- and aggression-related circuits; and (2) strong inhibitory projections from anxiety-promoting regions onto the aggression circuit, enabling elevated anxiety states to suppress aggressive behavior.

### Relationship between CORT, MUP and aggression

Although anxiety and aggression are highly correlated across strains, we did not find a clear relationship between baseline CORT level and aggression or anxiety-like behaviors. CORT level varies across strains but does not track the strain’s aggression or anxiety levels in any obvious way. This result is consistent with twin studies in humans, which have shown that hair cortisol concentration has a high heritability (72%), indicating that it is genetically determined. However, it has no apparent correlation with hypothalamic-pituitary-adrenal (HPA) axis activity, responses to stress, or depressive symptoms^99^. Indeed, although CORT is best known as a stress hormone, its primary function is to control metabolic rate by promoting gluconeogenesis in the liver. Numerous studies have shown that CORT levels alter with factors expected to affect energy expenditure, such as seasons^100^, time of day^101^, and brood size^102^. One study in zebrafish measured CORT levels when the animals increased energy expenditure in response to various stressors and non-stressors, and concluded that CORT is better explained by variation in metabolic rates than by stress^103^. Other studies have examined the acute and chronic effects of exercise on CORT and found that both induce an increase in CORT^104^. Thus, while CORT clearly increases during acute stress to prepare the body for energy-demanding motor responses, it is not a specific stress indicator, as its level could be affected by many stress-unrelated factors. This may explain the poor correlation between the baseline CORT level and anxiety or aggression.

Despite the absence of a simple linear relationship between CORT and aggression in our study, CORT is still likely to modulate aggressive behavior. Previous studies have shown that chronic CORT elevation suppresses aggression across species^105–108^ and in group-living animals, less aggressive subordinate individuals consistently exhibit higher baseline CORT levels than dominant, more aggressive animals^109–112^. Consistent with this, we found that strains with high baseline CORT, including BC and 129, display low levels of aggression. However, DBA mice, which exhibit low baseline CORT, are also minimally aggressive. Together, these findings suggest that while elevated CORT may constrain aggression, low baseline CORT does not guarantee high aggression.

Notably, CORT is dynamically regulated and increases acutely under challenging conditions. Prior studies in humans and rats have reported acute CORT elevations during aggressive encounters, and artificial CORT injection can facilitate attack initiation^113–115^. However, our previous study in mice found minimal activation of PVN CRH neurons during aggression^116^, highlighting potential species differences. Nevertheless, it remains possible that acute CORT responses to stressors, which were not measured here, are better correlated with aggression.

We found that MUP levels are uncorrelated with aggression across strains. This is not unexpected, as MUPs are synthesized by hepatocytes in the liver, and the genetic determinants of hepatic transcriptional activity are unlikely to substantially overlap with those governing neural circuits of aggression. Thus, although MUP levels vary robustly across strains, this variation appears independent of aggression. Interestingly, however, within the same strain, MUP levels positively correlated with aggression across individuals. This finding is consistent with our recent study in C57 male mice, showing that MUP levels not only predict future aggression in naïve animals but also increase in repeated winners that exhibit heightened aggression^15^. The positive relationship between MUP levels and individual aggressiveness likely reflects shared endocrine regulation. In particular, MUP synthesis is influenced by growth hormone and testosterone, both of which also strongly modulate aggressive behavior. For example, castration reduces aggression, whereas testosterone replacement restores it^117^, and growth hormone–releasing hormone (GHRH) knockout mice exhibit reduced aggression that is rescued by growth hormone administration^118^. Together, these findings suggest that, when genetic background is controlled, MUP levels can track individual variation in aggression, potentially through their shared neuroendocrine modulators.

Altogether, our study provides new insights into the neural implementation of genetic control of aggression, highlighting the critical role of intrinsic cell properties in determining behavioral traits in naïve animals. Our study also supports a tight link between anxiety and aggression, likely reflecting interactions of the underlying circuits or their shared modulatory mechanisms.

## Methods

### Mice

All procedures were approved by the NYULMC Institutional Animal Care and Use Committee (IACUC) in compliance with the National Institutes of Health (NIH) Guidelines for the Care and Use of Laboratory Animals. Mice were housed under a 12-hour light-dark cycle (dark cycle: 10 am to 10 pm), with food and water available *ad libitum*. Room temperature was maintained between 20–22 °C and humidity between 30–70%, with a daily average of approximately 45%. Experimental animals included adult wildtype CD-1 (Strain No. 022), Swiss Webster (Strain No. 551), FVB (Strain No. 559 or 021), C57BL/6 (Strain No. 027), DBA/2 (Strain No. 026), 129/SV-E (Strain No. 476), and BALB/c (Strain No. 028) male mice, all purchased directly from Charles River. They were 6-8 weeks old when they arrived at NYULMC. Upon arrival, the test mice were group housed until 8 weeks old, and single housed afterward. Intruder mice for the resident-intruder (RI) tests were group-housed adult male BALB/c, C57BL/6, and 129 mice (>8 weeks old, ∼25 g), as well as BALB/c juvenile male mice (4-5 weeks, ∼15 g). BALB/c and C57BL/6 intruders were originally sourced from Charles River and then bred in-house. 129 intruder mice were purchased from Charles River. For female experiments, female SW (Strain No. 024) and C57/BL6N (Strain No. 027) mice were purchased from Charles River. They were 6 weeks old when they arrived at NYULMC and were single-housed at 7 weeks. The behavioral tests were conducted after one week of single housing. All behavioral experiments were conducted during the animals’ dark cycle.

Each experiment was conducted using 1–6 cohorts of animals, and data from different cohorts were combined for the final analysis. Details regarding cohort numbers are provided in **Supplementary Table 1.**

### Resident Intruder Test

For the male resident intruder test, the test mice were single-housed resident mice. Intruder mice were randomly selected from a group of 20–30 mice housed in groups of 4–5 per cage. We used group-housed BC adult males as intruders for all strains except BC, for which group-housed C57 males were used. Group-housed BC intruders were chosen as our previous studies^15,35,58,61,63,73,74,87^ showed that they rarely initiate attacks as intruders, thereby allowing the resident to fully control the occurrence and intensity of aggressive interactions. To further reinforce this submissive phenotype, we pre-defeated BC and C57 intruders by introducing them to a separate cohort of single-housed SW or CD1 resident cages for at least 3 times, 10 minutes each time. After the last defeat, the intruder rested for more than 2 days before being used in the experiments. For resident intruder test towards the same-strain adult male or juvenile intruders, the intruder mice were group-housed but not pre-defeated. For the female resident-intruder test, a juvenile male BALB/c mouse was introduced into the test mouse’s home cage for 10 min. The juvenile intruders were not pre-defeated.

### Light Dark Box

The light-dark test was performed using a previously reported method with minor modifications^119^. The light-dark box consists of a starting dark box (W × L × H: 40 cm × 20 cm × 35 cm) and a light box (W × L × H: 40 cm × 20 cm × 35 cm), separated by a divider with an opening door (Stoelting, #63101). Mice were placed into the dark box at the start and allowed to move freely for 10 minutes. The latency to enter the light box, the number of entries to the light box, the time spent in the light box, and the duration of each entry were recorded and analyzed using the ANY-maze software (Stoelting Co.). The illumination at the center of the light box was approximately 100 lux.

### Open Field Test

The open field test was performed in a square arena (40 cm × 40 cm × 35 cm) (Stoelting, #60101) following the previously described method^120^. Each animal was introduced into the center of the open field illuminated at approximately 150 lux. The center area was defined as the inner four squares when the open field arena was divided into 16 equal squares.

### Elevated Plus Maze Test

The elevated plus maze test was performed in a customized mouse elevated plus maze, which consists of four arms (two open without walls and two enclosed by 15 cm high walls), 40 cm long and 5 cm wide. Each arm of the maze is attached to sturdy metal legs such that it is elevated 30 cm off the table. Animals were placed at the junction of the open and closed arms and allowed to freely explore the apparatus for 5 minutes.

### Home cage locomotion test

We measured the animal’s locomotion in its home cage without a lid for 3 minutes before the RI test. Animals were pre-habituated to the recording room for more than 30 minutes before any behavioral test. The cage lid, water bottle, and food were removed at least 5 min before the intruder was introduced. Trajectories and total distance traveled during the pre-intruder habitation period were analyzed using DeepLabCut v3.0^121^ and customized Python v3.11 script. The body centroid was tracked, pixel coordinates were converted to centimeters using a 30-cm in-frame calibration, low-confidence frames (likelihood ≤ 0.75) and implausible jumps (adjacent-frame jumps > 10 cm) were interpolated, and the trajectory was lightly smoothed with a centered 5-frame rolling mean. Home-cage locomotion (m/min) was computed as the mean instantaneous speed across the entire recording period, including stationary frames.

### Behavioral recordings and analysis

Animal behaviors were recorded from a top-down perspective using a Basler acA640-120um camera and commercial video acquisition software (Norpix, StreamPix 8) in a semi-dark room illuminated with infrared light, with the frame rate at 25 frames per second. For the RI tests, attack was manually annotated frame-by-frame by an experienced observer. “Attack” was characterized by a sequence of high-intensity actions directed toward the intruder, including pushing, lunging, biting, tumbling, and episodes of rapid locomotion between these movements. Non-contact chasing was not considered as “attack”. An attack bout starts with the resident lunging towards the intruder, often followed by biting on the intruder’s back and tumbling, and ends when the two animals separate, either because the resident walks away or the intruder escapes. Total attack duration, attack bout duration and frequency, and latency to attack were analyzed using custom MATLAB code (The MathWorks, Inc., 2023) based on the frame-by-frame manual annotation. The duration of attack bout across all animals ranged from 0.2 to 24.28 s, with a mean duration of 1.73 s.

For the LDB and OF tests, animal body centers were tracked using ANY-maze software. In LDB, entry into the light box was defined as the tracked body center entering the light box area. Immobility was defined as the absence of detectable movement of the body center for ≥ 2s, even if minor movements such as grooming were present. Total duration spent in the light box, number of light box entries, and latency to the first entry were analyzed. Bout duration was calculated by dividing the total time spent in the light compartment by the number of entries. Tracking data were also used to generate the movement traces.

For the OF test, the open-field arena was divided into 16 equally sized squares (10 × 10 cm each), with the central area defined as the innermost four squares and the periperial area as the remaining 12 squres. Because the animal was initially placed in the center area, center entries were counted only after the animal first exited the center and subsequently re-entered it. Center entry was determined by the presence of the body center in the center area. Immobility was defined as the animal remaining stationary, with no detectable movement (the minimal velocity is 1cm/s) of the body point for ≥ 2s, even if minor movements such as grooming were present. Mobile time in the arena was calculated as the total duration in the arena minus the immobile duration. The total distance traveled was calculated as the accumulated displacement of the tracked body center over the test duration, calculated using a smoothed path that corrects for small tracking jitters. The percentage of time in center was calculated as 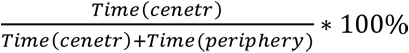. All behavioral metrics were obtained using ANY-maze software and customized MATLAB or Python codes.

For the strain-normalized values, we first calculated the strain mean (µ) and standard deviation (σ) using values from all animals of the same strain. We then calculated the strain normalized value (Z) for each animal (x) as: 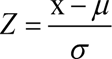.

Clustering analyses were performed on strain-level behavioral data using three aggression-related features: percentage of animals exhibiting attacks, attack latency, and total attack duration. Variables were z-scored before analysis. The optimal number of clusters was determined using the elbow method based on the within-cluster sum of squares, which indicated k = 3. K-means clustering was subsequently applied to group strains, and hierarchical clustering based on Euclidean distance and agglomerative linkage was performed to visualize the resulting cluster structure.

Partial regression analyses were conducted at the strain level (n = 7) to determine whether anxiety-related behavioral measures predicted aggression while controlling for locomotor activity. All variables were z-scored prior to analysis. Anxiety-related measures (LDB entry number and percentage of time in center) and the aggression-related measure (total attack duration) were each separately regressed against home-cage locomotor velocity, and the resulting residuals were extracted and plotted with a least-squares regression line. Pearson correlations between residuals were used to quantify partial correlations. Statistical significance was assessed using a multiple linear regression model that included both anxiety-related measures and home-cage locomotion as predictors. Regression coefficients were estimated using ordinary least-squares fitting, and significance for each predictor was determined using two-tailed t-tests on the corresponding regression coefficients based on their estimated standard errors within the full model.

### Measure CORT and urinary protein

To measure corticosterone and urinary protein levels, we collected urine from test animals between 11:00 am and 12:00 pm, which is one to two hours after light off. During urine collection, each animal was placed in a clean cage without any bedding material for 10 minutes. If the mouse did not urinate on day 1, we repeated the same procedure on the following day. Mice that failed to urinate on both days were excluded.

After urine collection, the samples were stored at -80 ℃ until use. The total protein and creatinine levels were measured using QuantiChrom Protein Creatinine Ratio Assay Kit (BioAssay Systems, #DPCR-100). The urinary protein-to-creatinine ratio was used as an approximation of MUP level, given that MUP constitutes approximately 90% of all urinary proteins^51^. Using the same urine sample, we measured the corticosterone concentration using an ELISA Kit (Arbor Assays™, K014) according to the manufacturer’s instructions. Briefly, samples were loaded into the pre-coated plate with antibody and corticosterone-peroxidase conjugate. After incubation for 1 hour on a high-speed shaker (600 rpm), the plate was washed four times, and the substrate was added. Following a 30-minute incubation without shaking, the stop solution was added, and optical absorbance was subsequently measured. Corticosterone concentrations were quantified using a standard curve generated from samples of known concentrations and were normalized to creatinine levels measured in the same urine samples to control for variation in urine dilution.

### c-Fos analysis

For c-Fos induction, all animals were directly purchased from Charles River at the age of 8 weeks. After one week of single housing, male animals were exposed to a pre-defeated group-housed Balb/c male mouse placed under a mental pencil cup for 15 minutes. Female mice were exposed to a cupped juvenile Balb/c male intruder for 15 minutes. Control animals were presented with an empty cup for 15 minutes.

Eighty minutes after the interactions, the animals were deeply anesthetized and perfused with Phosphate-Buffered Saline (PBS), followed by 4% paraformaldehyde (PFA). After perfusion, brains were harvested, post-fixed in 4% PFA overnight at 4°C, and then cryoprotected in 30% (w/v) sucrose overnight until sunk at 4°C. The brains were then embedded in OCT compound and sectioned into 50-μm-thick slices using a CM1900 cryostat (Leica). Immunostaining for cFos was performed using a guinea pig anti-cFos primary antibody (1:1000, Synaptic Systems #226 308), followed by an Alexa Fluor 647-conjugated goat anti-guinea pig secondary antibody (1:1000, Invitrogen A21450). Nuclear staining was achieved by including DAPI (1:20,000, Thermo Fisher, D1306) in the secondary antibody staining solution. Fluorescence images were acquired using an Olympus VS120 virtual slide scanner with identical imaging settings across all animals.

For quantification, the following brain regions were manually outlined bilaterally based on DAPI staining, using anatomical landmarks defined in the Paxinos and Franklin Mouse Brain Atlas and the Allen Brain Atlas: LSd (bregma AP: 0.75 to -0.05 mm), LSv (0.75 to -0.05 mm), BNSTpr (−0.1 to −0.6 mm), MPOA (0.1 to −0.5 mm), AHN (−0.8 to −1.1 mm), PVN (−0.8 to −1.1 mm), DMH (−1.4 to −2.0 mm), aVMH (−1.1 to −1.3 mm), aVMHvl (−1.3 to −1.6 mm), pVMHvl (−1.6 to −1.9 mm), VMHdm/c (−1.3 to −1.9 mm), TU (−1.4 to −1.8 mm), MeApd (−1.4 to −1.8 mm), MeApv (−1.4 to −1.8 mm), COApl (−1.5 to −2.0 mm), PMd (−2.4 to −2.6 mm), PMv (−2.4 to −2.6 mm), PA (−2.2 to −2.8 mm), PAGd, PAGdl, PAGl, and PAGv (−3.55 to −4.2 m). The number of sections stained and quantified for each region is listed in **Supplementary Table 2**. For 3 males of each strain, we quantified every 4^th^ section, and for 2 males of each strain and all females, we quantified every other section. Five sections were excluded due to poor sectioning or staining quality. cFos-positive cells and DAPI were counted using QuPath (v0.5.0) open-source software. Automatic segmentation was performed using the following parameters across all images: requested pixel size = 2 μm, background radius = 8 μm, median filter radius = 0 μm, sigma = 1.5 μm, intensity threshold = 50, and cell expansion = 5 μm. All images were analyzed under identical contrast settings to ensure consistency in fluorescence signal detection.

### Stereotaxic surgery

During the surgery, mice were anesthetized with 1-1.5% isoflurane and placed in a stereotactic frame (Kopf Instruments, Model 1900). Stereotaxic injection coordinates were based on the Paxinos and Franklin mouse brain atlas.

For visualizing pVMHvl in the male patch clamp experiment, we injected 100 nL/side of red retrobeads (Lumafluor Inc, Red Retrobeads™ IX) into the MPOA bilaterally (AP: -0.30 mm, ML: ±0.40 mm, DV: -5.00 mm) of 7-week-old wildtype mice through a glass capillary using a nanoinjector (World Precision Instruments, Nanoliter 2000) at a speed of 20 nL/min. Retrobeads were diluted 1:3 with ddH_2_O according to the manufacturer’s protocol before injections.

For visualizing pVMHvlm in female patch clamp experiment, we injected 100 nL/side of CTB Alexa 488 (Thermofisher Scientific, C34775) into the MPOA (AP: 0.31 mm, ML: ±0.30 mm, DV: 4.95 mm) and red retrobeads (Lumafluor Inc, Red Retrobeads™ IX) into AVPV bilaterally (AP: - 0.60 mm, ML: ±0.15 mm, DV: 4.90 mm) through a glass capillary using a nanoinjector (World Precision Instruments, Nanoliter 2000) at a speed of 20 nL/min in 10-12 weeks old wildtype female mice.

To chemogenetically activate VMHvl neurons in 129 male mice, we bilaterally injected 300 nL of a Cre-dependent hM3Dq virus (AAV2-hSyn-DIO-hM3D(Gq)-mCherry; Addgene #44361; titer: 8.7 × 10¹² vg/mL) or a control virus (AAV2-hSyn-DIO-mCherry; Addgene #50459; titer: 4.4 × 10¹² vg/mL) into the VMHvl (AP −1.58 mm, ML ±0.775 mm, DV −5.65 mm). To restrict expression to VMHvl neurons projecting to the MPOA, 200 nL of RetroAAV-EF1a-Cre (Addgene #55636) was injected into the MPOA (AP −0.30 mm, ML ±0.40 mm, DV −5.00 mm) of 7-week-old wild-type male 129 mice.

### Chemogenetic activation

After surgery, animals were single-housed for three weeks. On the day of testing, the animals were i.p. injected with either saline or (1 mg/kg) CNO. 6/10 animals received saline injection on the first day, while the remaining 4 mice were injected with CNO first. 30 minutes after injection (saline or CNO), a non-aggressive juvenile BALB/c male was introduced into the test male’s home cage for 10 minutes.

Following the experiment, mice were transcardially perfused with PBS followed by 4% PFA. Brains were post-fixed (3–4 h, 4 °C), cryoprotected in 20% sucrose for 3-4 days, embedded in OCT, and sectioned coronally at 50 μm using a cryostat. Free-floating sections were blocked in PBS-T (0.3% Triton X-100) with 10% normal donkey serum and incubated with primary antibody against ESR1 (ESR1; rabbit anti-ESR1, 1:1,000, Invitrogen, PA1-309) at 4 °C overnight. The sections were then washed with 0.3% PBST X 3 times and incubated with fluorescent secondary antibodies (donkey anti-rabbit Alexa Fluor-conjugated, 1:1,000; Jackson ImmunoResearch) and DAPI (1:1,000; Thermo Fisher Scientific) for 2 h at room temperature. Sections were mounted onto APS adhesive slides (MATSUNAMI GLASS SUAPS19), coverslipped with glass coverslips, and imaged using either a confocal microscope (Zeiss LSM series) or a slide-scanning system (Olympus VS120) to verify viral expression and injection sites.

To quantify the percentage of mCherry-positive cells that were also ESR1-positive, confocal z-stacks were maximum-intensity projected in ImageJ and imported into QuPath. mCherry-positive cells were identified based on the mCherry and DAPI channels, while the ESR1 channel was initially hidden to minimize annotation bias. The ESR1 channel was then displayed to classify each mCherry-positive cell as ESR1-positive or ESR1-negative. Two sections per animal were quantified, and the resulting percentages were averaged to obtain one value per animal.

### Pharmacological administration

For the pharmacological manipulation experiment, single-housed adult male CD1 mice (12–13 weeks old) received intraperitoneal injections of yohimbine (YO, 2 mg/kg; Thermo Scientific, 141030050) or saline. All mice were first injected with saline and with YO 3 hours after the first test. Thirty minutes after each injection, a non-aggressive adult BALB/c male was introduced into the test male’s home cage for 10 min to assess aggression in a resident-intruder test. To control for potential effects of repeated RI testing on baseline aggression, a saline–saline group was included and tested two weeks after the saline–YO condition. One week after the last resident-intruder test, the same animals were tested in the LDB test after saline and YO injections on separate days.

### Patch clamp slice recording

Following retrobeads injection, mice were group-housed for 1 week to recover from surgery, then single-housed for an additional week before slice electrophysiology. For in vitro whole-cell patch-clamp recordings, mice were anesthetized with isoflurane and perfused with 15 mL oxygenated ice-cold cutting solution containing (in mM) 110 choline chloride, 25 NaHCO_3_, 2.5 KCl, 7 MgCl_2_, 0.5 CaCl_2_, 1.25 NaH_2_PO_4_, 25 glucose, 11.6 ascorbic acid, and 3.1 pyruvic acid. The coronal pVMHvl brain sections (275 µM in thickness) were cut using the Leica VT1200s vibratome and collected into the oxygenated artificial cerebrospinal fluid (ACSF) solution containing (in mM) 125 NaCl, 2.5 KCl, 1.25 NaH_2_PO_4_, 25 NaHCO_3_, 1 MgCl_2_, 2 CaCl_2,_ and 11 glucose at 32 ℃. Then, the sections were transferred to room temperature to incubate for at least 30 minutes until use. During recordings, slices were perfused with oxygenated ACSF at a constant flow rate in a recording chamber. Whole-cell patch-clamp recordings were performed using a MultiClamp 700B amplifier (Molecular Devices), digitized at 20 kHz with a Digidata 1550B interface (Molecular Devices), and acquired using Clampex 11.0 software (Axon Instruments). Data were analyzed using Clampfit (Molecular Devices) and MATLAB R2023a. Only cells with input resistance > 200 MΩ, access resistance (Ra) < 30 MΩ, and leak current within ±30 pA were included for analysis.

For current-clamp recordings, the internal pipette solution contained (in mM): 145 K-gluconate, 2 MgCl₂, 2 Na₂ATP, 10 HEPES, and 0.2 EGTA (286 mOsm, pH 7.2). Cells were subjected to a series of 500-ms current injections ranging from −20 pA to 270 pA in 10 pA increments. Prior to analysis, raw electrophysiological traces were low-pass filtered using a second-order Butterworth filter with a 1 kHz cutoff frequency to reduce high-frequency noise. Spikes were detected using custom scripts in MATLAB R2023a with a threshold of 10 mV. The rheobase was defined as the minimum current required to evoke an action potential. The resting membrane potential was calculated as the average membrane potential during the 400-ms baseline period of sweeps without spontaneous spiking. Input resistance was obtained from the membrane test window following break-in (Clampex 11.0). The Input-Frequency (IF) curve was constructed by counting action potentials at each current step, and the maximum spike number per cell was defined as the highest spike count observed across all steps.

For each cell, an average spike waveform was generated by aligning and averaging the first spike (detection threshold: 10 mV) across all depolarizing sweeps within a ±3 ms window around the spike peak; sweeps without spikes were excluded. Strain-level spike waveforms were then generated by averaging across cells of the strain, with the SEM shown as a shaded region. Phase plots were constructed by calculating the first derivative (dV/dt) of the strain-averaged waveform and plotting dV/dt against membrane potential (V), with SEM shading indicating across-cell variability. The spike threshold was estimated using a second derivative phase plot method^122^, defined as the membrane potential at which d²V/dt² first exceeded 5% of its peak during the rising phase, which captures the rapid acceleration of depolarization that marks the onset of spike initiation. The maximum rate of rise was calculated as the maximum dV/dt within the spike window, and the spike peak amplitude was defined as the maximum voltage within the spike window. The rate of repolarization was determined as the most negative dV/dt following the spike peak, representing the steepest repolarization slope. AP half-width was defined as the spike duration at half-maximal amplitude (i.e., the midpoint between spike threshold and peak amplitude) and was calculated as the time difference between the rising and falling phase crossings at this voltage level, identified using linear interpolation. All spike waveform metrics were computed from the first action potential evoked during each depolarizing current step and subsequently averaged for each cell.

For voltage-clamp recordings of spontaneous excitatory and inhibitory postsynaptic currents (sEPSCs and sIPSCs), the internal solution contained (in mM): 135 CsMeSO₃, 10 HEPES, 1 EGTA, 3.3 QX-314 (Cl⁻ salt), 4 MgATP, 0.3 NaGTP, and 8 sodium phosphocreatine (pH 7.3, adjusted with CsOH). The membrane potential was held at −70 mV for sEPSC recordings and at 0 mV for sIPSC recordings. The total recording was eight 5s-trails for either sEPSC or sIPSC, and there was a 5s recording gap between adjacent trails. Both amplitude and frequency of sEPSCs and sIPSCs were quantified using Clampfit (Molecular Devices). The E/I ratio was calculated as:

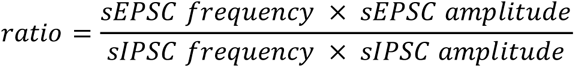

### Statistics

Fisher’s exact test was used for comparing proportions. Pairwise group comparisons were performed using two-sided tests on animal counts, with p-values adjusted for multiple comparisons using the Benjamini–Hochberg false discovery rate (FDR) procedure (FDR = 0.05; MATLAB mafdr, ’BHFDR’, true). Data normality was assessed using the D’Agostino and Pearson omnibus test. For comparison across two treatments in the same group, the paired t test was used for normally distributed datasets while wilcoxon test was used for non-normally distributed datasets. For comparison across two groups, the unpaired t test was used for normally distributed datasets while Mann–Whitney U test was used for non-normally distributed datasets. For comparisons across multiple groups, one-way ANOVA was used when all groups met normality assumptions, while the Kruskal–Wallis test was applied for non-normally distributed data. Post hoc pairwise comparisons were corrected for multiple testing using the two-stage step-up method of Benjamini, Krieger, and Yekutieli to control the FDR. For I–F curves, group differences were assessed using two-way ANOVA and two-way RM ANOVA tests, followed by a two-stage linear step-up procedure of Benjamini, Krieger, and Yekutieli FDR correction (FDR = 0.05). Correlation between variables was evaluated using Pearson’s correlation coefficient. Correlation coefficients (r) and associated p-values were computed using the corr function in MATLAB R2023a. All other analyses were performed using GraphPad Prism version 10.5.0 (673). See Supplementary Table 1 for statistical details.

## Data Availability

Behavior annotations, raw representative histology images, optical absorbance data for urinary protein and corticosterone assays, slice electrophysiology recording data will be deposited in a public database before publication. Behavior videos and additional histology images are available from the corresponding author upon reasonable request.

## Code Availability

All MATLAB code used for analysis will be deposited in a public database and made accessible before publication.

## Author Contributions

D.L. conceived and supervised the project. D.L. and X.D. designed the experiments and co-wrote the manuscript. X.D. performed most behavioral, c-Fos, and slice electrophysiological recording experiments in males and conducted data analysis. Y.W. contributed to behavioral, functional manipulation experiments and slice electrophysiology recording experiments. T.Y. performed behavioral experiments, surgeries, and c-Fos experiments in female mice. M.G. conducted c-Fos, functional manipulation experiments, and related analyses in male mice. E.R. performed electrophysiological recordings in female animals. P.D. contributed to histology experiments. B.D. and J.C. assisted with sample collection and pilot experiments.

## Acknowledgements

We thank Rongzhen Yan for advice on in vitro recording experiments, Long Mei for the drawings of the mouse brain atlas (Fig. 4 and Sup. Fig. 4), and Wenxi Zhou for the helpful discussion. Mouse cartoons in Fig. 4a were from BioRender (https://biorender.com). This research was supported by NIH grants R01MH124927, R01MH101377, U19NS107616 (D.L.), the Vulnerable Brain Project (D.L.), and the Brain and Behavior Research Foundation (D.L.).

## References

1 Lischinsky, J. E. & Lin, D. Neural mechanisms of aggression across species. Nature Neuroscience 23, 1317–1328 (2020). 10.1038/s41593-020-00715-2

2 Baron, R. A. & Richardson, D. R. Human aggression. (Springer Science & Business Media, 1994).

3 Rowell Huesmann, L. & Eron, L. D. Individual differences and the trait of aggression. European Journal of personality 3, 95–106 (1989).

4 Barr, C. S. & Driscoll, C. in Neuroscience of Aggression (eds Klaus A. Miczek & Andreas Meyer-Lindenberg) 45–71 (Springer Berlin Heidelberg, 2014).

5 Ferrari, P. F., Palanza, P., Parmigiani, S. & Rodgers, R. J. Interindividual variability in Swiss male mice: relationship between social factors, aggression, and anxiety. Physiology & behavior 63, 821–827 (1998).

6 Gaskill, B. N. et al. The effect of early life experience, environment, and genetic factors on spontaneous home-cage aggression-related wounding in male C57BL/6 mice. Lab Animal 46, 176–184 (2017).

7 Lathe, R. The individuality of mice. *Genes*, Brain and Behavior 3, 317–327 (2004). 10.1111/j.1601-183X.2004.00083.x

8 Eichelman, B. Variability in rat irritable and predatory aggression. Behavioral & Neural Biology 29, 498–505 (1980). 10.1016/S0163-1047(80)92768-5

9 de Boer, S. F., van der Vegt, B. J. & Koolhaas, J. M. Individual Variation in Aggression of Feral Rodent Strains: A Standard for the Genetics of Aggression and Violence? Behavior genetics 33, 485–501 (2003). 10.1023/A:1025766415159

10 Bishop, A. M., Pomeroy, P. & Twiss, S. D. Variability in individual rates of aggression in wild gray seals: fine-scale analysis reveals importance of social and spatial stability. Behavioral Ecology and Sociobiology 69, 1663–1675 (2015). 10.1007/s00265-015-1978-x

11 Cervantes, M. C. & Delville, Y. Individual differences in offensive aggression in golden hamsters: A model of reactive and impulsive aggression? Neuroscience 150, 511–521 (2007). 10.1016/j.neuroscience.2007.09.034

12 Elizabeth Bolhuis, J., Schouten, W. G. P., Schrama, J. W. & Wiegant, V. M. Individual coping characteristics, aggressiveness and fighting strategies in pigs. Animal Behaviour 69, 1085–1091 (2005). 10.1016/j.anbehav.2004.09.013

13 Svartberg, K. Individual differences in behaviour—dog personality. The behavioural biology of dogs, 182–206 (2007).

14 Mikkola, S., Salonen, M., Hakanen, E., Sulkama, S. & Lohi, H. Reliability and Validity of Seven Feline Behavior and Personality Traits. Animals 11 (2021). 10.3390/ani11071991

15 Yan, R. et al. The multi-stage plasticity in the aggression circuit underlying the winner effect. Cell 187, 6785–6803 e6718 (2024). 10.1016/j.cell.2024.09.030

16 Rose, J., Rillich, J. & Stevenson, P. A. Chronic social defeat induces long-term behavioral depression of aggressive motivation in an invertebrate model system. PLoS One 12, e0184121 (2017). 10.1371/journal.pone.0184121

17 Huhman, K. L. et al. Conditioned defeat in male and female syrian hamsters. Hormones and Behavior 44, 293–299 (2003). 10.1016/j.yhbeh.2003.05.001

18 Veenema, A. H., Blume, A., Niederle, D., Buwalda, B. & Neumann, I. D. Effects of early life stress on adult male aggression and hypothalamic vasopressin and serotonin. Eur J Neurosci 24, 1711–1720 (2006). 10.1111/j.1460-9568.2006.05045.x

19 Nordman, J. C., Bartsch, C. J. & Li, Z. Opposing effects of NMDA receptor antagonists on early life stress-induced aggression in mice. Aggress Behav 48, 365–373 (2022). 10.1002/ab.22022

20 Whitfield, C. L., Anda, R. F., Dube, S. R. & Felitti, V. J. Violent Childhood Experiences and the Risk of Intimate Partner Violence in Adults: Assessment in a Large Health Maintenance Organization. Journal of Interpersonal Violence 18, 166–185 (2003). 10.1177/0886260502238733

21 Brunner, H. G., Nelen, M., Breakefield, X. O., Ropers, H. H. & van Oost, B. A. Abnormal behavior associated with a point mutation in the structural gene for monoamine oxidase A. *Science (New York*, N.Y.) 262, 578–580 (1993). 10.1126/science.8211186

22 Cases, O. et al. Aggressive behavior and altered amounts of brain serotonin and norepinephrine in mice lacking MAOA. *Science (New York*, N.Y.) 268, 1763–1766 (1995). 10.1126/science.7792602

23 Rodriguiz, R. M., Chu, R., Caron, M. G. & Wetsel, W. C. Aberrant responses in social interaction of dopamine transporter knockout mice. Behavioural Brain Research 148, 185–198 (2004). 10.1016/S0166-4328(03)00187-6

24 Tuvblad, C. & Baker, L. A. Human aggression across the lifespan: genetic propensities and environmental moderators. Adv Genet 75, 171–214 (2011). 10.1016/B978-0-12-380858-5.00007-1

25 Le Roy, I. et al. Genetic correlation between steroid sulfatase concentration and initiation of attack behavior in mice. Behavior genetics 29, 131–136 (1999). 10.1023/a:1021664607131

26 Roubertoux, P. L. et al. Attack behaviors in mice: From factorial structure to quantitative trait loci mapping. European Journal of Pharmacology 526, 172–185 (2005). 10.1016/j.ejphar.2005.09.026

27 Brodkin, E. S., Goforth, S. A., Keene, A. H., Fossella, J. A. & Silver, L. M. Identification of quantitative trait Loci that affect aggressive behavior in mice. The Journal of neuroscience : the official journal of the Society for Neuroscience 22, 1165–1170 (2002). 10.1523/jneurosci.22-03-01165.2002

28 Natarajan, D., de Vries, H., Saaltink, D. J., de Boer, S. F. & Koolhaas, J. M. Delineation of violence from functional aggression in mice: an ethological approach. Behavior genetics 39, 73–90 (2009). 10.1007/s10519-008-9230-3

29 Ginsburg, B. & Allee, W. C. Some Effects of Conditioning on Social Dominance and Subordination in Inbred Strains of Mice. Physiological Zoology 15, 485–506 (1942).

30 Scott, J. P. GENETIC DIFFERENCES IN THE SOCIAL BEHAVIOR OF INBRED STRAINS OF MICE. Journal of Heredity 33, 11–15 (1942). 10.1093/oxfordjournals.jhered.a105080 %J Journal of Heredity

31 Osakada, T. et al. A dedicated hypothalamic oxytocin circuit controls aversive social learning. Nature 626, 347–356 (2024). 10.1038/s41586-023-06958-w

32 Golden, S. A. et al. Basal forebrain projections to the lateral habenula modulate aggression reward. Nature 534, 688–692 (2016). 10.1038/nature18601

33 Golden, S. A. et al. Nucleus Accumbens Drd1-Expressing Neurons Control Aggression Self-Administration and Aggression Seeking in Mice. The Journal of Neuroscience 39, 2482 (2019). 10.1523/JNEUROSCI.2409-18.2019

34 Hashikawa, Y., Hashikawa, K., Falkner, A. L. & Lin, D. Ventromedial Hypothalamus and the Generation of Aggression. Front Syst Neurosci 11, 94 (2017). 10.3389/fnsys.2017.00094

35 Lin, D. et al. Functional identification of an aggression locus in the mouse hypothalamus. Nature 470, 221–226 (2011). 10.1038/nature09736

36 Yamaguchi, T. et al. The neural mechanisms supporting the rise and fall of maternal aggression. Nature (2026). 10.1038/s41586-026-10354-5

37 Hashikawa, K. et al. Esr1(+) cells in the ventromedial hypothalamus control female aggression. Nat Neurosci 20, 1580–1590 (2017). 10.1038/nn.4644

38 Meier, S. M. & Deckert, J. Genetics of Anxiety Disorders. Current Psychiatry Reports 21, 16 (2019). 10.1007/s11920-019-1002-7

39 van Gaalen, M. M. & Steckler, T. Behavioural analysis of four mouse strains in an anxiety test battery. Behavioural Brain Research 115, 95–106 (2000). 10.1016/S0166-4328(00)00240-0

40 Griebel, G., Belzung, C., Perrault, G. & Sanger, D. J. Differences in anxiety-related behaviours and in sensitivity to diazepam in inbred and outbred strains of mice. Psychopharmacology 148, 164–170 (2000). 10.1007/s002130050038

41 Milner, L. C. & Crabbe, J. C. Three murine anxiety models: results from multiple inbred strain comparisons. *Genes*, Brain and Behavior 7, 496–505 (2008). 10.1111/j.1601-183X.2007.00385.x

42 Tang, X. & Sanford, L. D. Home cage activity and activity-based measures of anxiety in 129P3/J, 129X1/SvJ and C57BL/6J mice. Physiol Behav 84, 105–115 (2005). 10.1016/j.physbeh.2004.10.017

43 La-Vu, M., Tobias, B. C., Schuette, P. J. & Adhikari, A. To Approach or Avoid: An Introductory Overview of the Study of Anxiety Using Rodent Assays. Frontiers in Behavioral Neuroscience Volume 14 - 2020 (2020). 10.3389/fnbeh.2020.00145

44 Svare, B. B. & Leshner, A. I. Behavioral correlates of intermale aggression and grouping in mice. J Comp Physiol Psychol 85, 203–210 (1973). 10.1037/h0034897

45 Nyberg, J. M., Vekovischeva, O. & Sandnabba, N. K. Anxiety profiles of mice selectively bred for intermale aggression. Behavior genetics 33, 503–511 (2003). 10.1023/a:1025718531997

46 Fujita, O., Annen, Y. & Kitaoka, A. Tsukuba high- and low-emotional strains of rats (Rattus norvegicus): an overview. Behavior genetics 24, 389–415 (1994). 10.1007/BF01067540

47 Guillot, P.-V. & Chapouthier, G. Intermale aggression and dark/light preference in ten inbred mouse strains. Behavioural Brain Research 77, 211–213 (1996). 10.1016/0166-4328(95)00163-8

48 Berton, O., Ramos, A., Chaouloff, F. & Mormède, P. Behavioral Reactivity to Social and Nonsocial Stimulations: A Multivariate Analysis of Six Inbred Rat Strains. Behavior genetics 27, 155–166 (1997). 10.1023/A:1025641509809

49 Zheng, H. & Rinaman, L. Yohimbine anxiogenesis in the elevated plus maze requires hindbrain noradrenergic neurons that target the anterior ventrolateral bed nucleus of the stria terminalis. Eur J Neurosci 37, 1340–1349 (2013). 10.1111/ejn.12123

50 Lee, W., Khan, A. & Curley, J. P. Major urinary protein levels are associated with social status and context in mouse social hierarchies. Proc Biol Sci 284 (2017). 10.1098/rspb.2017.1570

51 Beynon, R. J. & Hurst, J. L. Urinary proteins and the modulation of chemical scents in mice and rats. Peptides 25, 1553–1563 (2004). 10.1016/j.peptides.2003.12.025

52 Hashikawa, K., Hashikawa, Y., Falkner, A. & Lin, D. The neural circuits of mating and fighting in male mice. Curr Opin Neurobiol 38, 27–37 (2016). 10.1016/j.conb.2016.01.006

53 Unger, E. K. et al. Medial amygdalar aromatase neurons regulate aggression in both sexes. Cell Rep 10, 453–462 (2015). 10.1016/j.celrep.2014.12.040

54 Hong, W., Kim, D. W. & Anderson, D. J. Antagonistic control of social versus repetitive self-grooming behaviors by separable amygdala neuronal subsets. Cell 158, 1348–1361 (2014). 10.1016/j.cell.2014.07.049

55 Miller, S. M., Marcotulli, D., Shen, A. & Zweifel, L. S. Divergent medial amygdala projections regulate approach-avoidance conflict behavior. Nat Neurosci 22, 565–575 (2019). 10.1038/s41593-019-0337-z

56 Yang, B., Karigo, T. & Anderson, D. J. Transformations of neural representations in a social behaviour network. Nature 608, 741–749 (2022). 10.1038/s41586-022-05057-6

57 Knoedler, J. R. et al. A functional cellular framework for sex and estrous cycle-dependent gene expression and behavior. Cell 185, 654–671 e622 (2022). 10.1016/j.cell.2021.12.031

58 Yamaguchi, T. et al. Posterior amygdala regulates sexual and aggressive behaviors in male mice. Nat Neurosci 23, 1111–1124 (2020). 10.1038/s41593-020-0675-x

59 Zha, X. et al. VMHvl-Projecting Vglut1+ Neurons in the Posterior Amygdala Gate Territorial Aggression. Cell Rep 31, 107517 (2020). 10.1016/j.celrep.2020.03.081

60 Aubry, A. V. et al. A crucial role for the cortical amygdala in shaping social encounters. Nature 639, 1006–1015 (2025). 10.1038/s41586-024-08540-4

61 Wong, L. C. et al. Effective Modulation of Male Aggression through Lateral Septum to Medial Hypothalamus Projection. Curr Biol 26, 593–604 (2016). 10.1016/j.cub.2015.12.065

62 Leroy, F. et al. A circuit from hippocampal CA2 to lateral septum disinhibits social aggression. Nature 564, 213–218 (2018). 10.1038/s41586-018-0772-0

63 Dai, B. et al. Experience-dependent dopamine modulation of male aggression. Nature 639, 430–437 (2025). 10.1038/s41586-024-08459-w

64 Mahadevia, D. et al. Dopamine promotes aggression in mice via ventral tegmental area to lateral septum projections. Nat Commun 12, 6796 (2021). 10.1038/s41467-021-27092-z

65 Wei, D. et al. A hypothalamic pathway that suppresses aggression toward superior opponents. Nat Neurosci 26, 774–787 (2023). 10.1038/s41593-023-01297-5

66 Goodson, J. L., Kelly, A. M., Kingsbury, M. A. & Thompson, R. R. An aggression-specific cell type in the anterior hypothalamus of finches. Proceedings of the National Academy of Sciences 109, 13847–13852 (2012). 10.1073/pnas.1207995109

67 Ferris, C. F. et al. Vasopressin/serotonin interactions in the anterior hypothalamus control aggressive behavior in golden hamsters. The Journal of neuroscience : the official journal of the Society for Neuroscience 17, 4331–4340 (1997). 10.1523/jneurosci.17-11-04331.1997

68 Lee, H. et al. Scalable control of mounting and attack by Esr1+ neurons in the ventromedial hypothalamus. Nature 509, 627–632 (2014). 10.1038/nature13169

69 Yang, C. F. et al. Sexually Dimorphic Neurons in the Ventromedial Hypothalamus Govern Mating in Both Sexes and Aggression in Males. Cell 153, 896–909 (2013). 10.1016/j.cell.2013.04.017

70 Stagkourakis, S. et al. A neural network for intermale aggression to establish social hierarchy. Nature Neuroscience 21, 834–842 (2018). 10.1038/s41593-018-0153-x

71 Chen, A. X. et al. Specific Hypothalamic Neurons Required for Sensing Conspecific Male Cues Relevant to Inter-male Aggression. Neuron 108, 763–774.e766 (2020). 10.1016/j.neuron.2020.08.025

72 Motta, S. C. et al. Ventral premammillary nucleus as a critical sensory relay to the maternal aggression network. Proceedings of the National Academy of Sciences 110, 14438–14443 (2013). 10.1073/pnas.1305581110

73 Falkner, A. L. et al. Hierarchical Representations of Aggression in a Hypothalamic-Midbrain Circuit. Neuron 106, 637–648.e636 (2020). 10.1016/j.neuron.2020.02.014

74 Wang, L. et al. Hypothalamic Control of Conspecific Self-Defense. Cell Rep 26, 1747–1758.e1745 (2019). 10.1016/j.celrep.2019.01.078

75 Minakuchi, T. et al. Independent inhibitory control mechanisms for aggressive motivation and action. Nat Neurosci 27, 702–715 (2024). 10.1038/s41593-023-01563-6

76 Lo, L. et al. Connectional architecture of a mouse hypothalamic circuit node controlling social behavior. Proc Natl Acad Sci U S A 116, 7503–7512 (2019). 10.1073/pnas.1817503116

77 Yin, L. et al. VMHvll(Cckar) cells dynamically control female sexual behaviors over the reproductive cycle. Neuron 110, 3000–3017.e3008 (2022). 10.1016/j.neuron.2022.06.026

78 Roth, B. L. DREADDs for Neuroscientists. Neuron 89, 683–694 (2016). 10.1016/j.neuron.2016.01.040

79 Yang, T. et al. Social Control of Hypothalamus-Mediated Male Aggression. Neuron 95, 955–970.e954 (2017). 10.1016/j.neuron.2017.06.046

80 Hsieh, L. S., Wen, J. H., Miyares, L., Lombroso, P. J. & Bordey, A. Outbred CD1 mice are as suitable as inbred C57BL/6J mice in performing social tasks. Neuroscience letters 637, 142–147 (2017). 10.1016/j.neulet.2016.11.035

81 Dow, H. C. et al. Genetic dissection of intermale aggressive behavior in BALB/cJ and A/J mice. Genes Brain Behav 10, 57–68 (2011). 10.1111/j.1601-183X.2010.00640.x

82 Ciaranello, R. D., Lipsky, A. & Axelrod, J. Association between fighting behavior and catecholamine biosynthetic enzyme activity in two inbred mouse sublines. Proc Natl Acad Sci U S A 71, 3006–3008 (1974). 10.1073/pnas.71.8.3006

83 Velez, L., Sokoloff, G., Miczek, K. A., Palmer, A. A. & Dulawa, S. C. Differences in aggressive behavior and DNA copy number variants between BALB/cJ and BALB/cByJ substrains. Behavior genetics 40, 201–210 (2010). 10.1007/s10519-009-9325-5

84 Lindenfors, P. & S.Tullberg, B. in Advances in Genetics Vol. 75 (eds Robert Huber, Danika L. Bannasch, & Patricia Brennan) 7–22 (Academic Press, 2011).

85 Maxson, S. C. in Handbook of Behavior Genetics (ed Yong-Kyu Kim) 301–316 (Springer New York, 2009).

86 Mentis, A.-F. A., Dardiotis, E., Katsouni, E. & Chrousos, G. P. From warrior genes to translational solutions: novel insights into monoamine oxidases (MAOs) and aggression. Translational Psychiatry 11, 130 (2021). 10.1038/s41398-021-01257-2

87 Guo, Z. et al. Neural dynamics in the limbic system during male social behaviors. Neuron 111, 3288–3306 e3284 (2023). 10.1016/j.neuron.2023.07.011

88 Mei, L., Yan, R., Yin, L., Sullivan, R. M. & Lin, D. Antagonistic circuits mediating infanticide and maternal care in female mice. Nature 618, 1006–1016 (2023). 10.1038/s41586-023-06147-9

89 Slattery, D. A. & Neumann, I. D. No stress please! Mechanisms of stress hyporesponsiveness of the maternal brain. J Physiol 586, 377–385 (2008). 10.1113/jphysiol.2007.145896

90 Sandnabba, N. K. Selective breeding for isolation-induced intermale aggression in mice: associated responses and environmental influences. Behavior genetics 26, 477–488 (1996). 10.1007/BF02359752

91 Cairns, R. B., MacCombie, D. J. & Hood, K. E. A developmental-genetic analysis of aggressive behavior in mice: I. Behavioral outcomes. J Comp Psychol 97, 69–89 (1983).

92 Nehrenberg, D. L. et al. An anxiety-like phenotype in mice selectively bred for aggression. Behav Brain Res 201, 179–191 (2009). 10.1016/j.bbr.2009.02.010

93 Beiderbeck, D. I. et al. High and abnormal forms of aggression in rats with extremes in trait anxiety – Involvement of the dopamine system in the nucleus accumbens. Psychoneuroendocrinology 37, 1969–1980 (2012). 10.1016/j.psyneuen.2012.04.011

94 Zarrindast, M. R. & Khakpai, F. The Modulatory Role of Dopamine in Anxiety-like Behavior. Arch Iran Med 18, 591–603 (2015).

95 Yamaguchi, T. & Lin, D. Functions of medial hypothalamic and mesolimbic dopamine circuitries in aggression. Current Opinion in Behavioral Sciences 24, 104–112 (2018). 10.1016/j.cobeha.2018.06.011

96 Yan, J. J. et al. A circuit from the ventral subiculum to anterior hypothalamic nucleus GABAergic neurons essential for anxiety-like behavioral avoidance. Nat Commun 13, 7464 (2022). 10.1038/s41467-022-35211-7

97 Hong, C. Y., Din, J. S., Chang, H., Bang, J. Y. & Kim, J. C. Anterior hypothalamic nucleus drives distinct defensive responses through cell-type-specific activity. iScience 28, 112097 (2025). 10.1016/j.isci.2025.112097

98 Davis, M., Walker, D. L., Miles, L. & Grillon, C. Phasic vs sustained fear in rats and humans: role of the extended amygdala in fear vs anxiety. Neuropsychopharmacology 35, 105–135 (2010). 10.1038/npp.2009.109

99 Rietschel, L. et al. Hair Cortisol in Twins: Heritability and Genetic Overlap with Psychological Variables and Stress-System Genes. Scientific Reports 7, 15351 (2017). 10.1038/s41598-017-11852-3

100 Schradin, C. Seasonal changes in testosterone and corticosterone levels in four social classes of a desert dwelling sociable rodent. Horm Behav 53, 573–579 (2008). 10.1016/j.yhbeh.2008.01.003

101 Bailey, S. L. & Heitkemper, M. M. Circadian rhythmicity of cortisol and body temperature: morningness-eveningness effects. Chronobiol Int 18, 249–261 (2001). 10.1081/cbi-100103189

102 Bonier, F., Moore, I. T. & Robertson, R. J. The stress of parenthood? Increased glucocorticoids in birds with experimentally enlarged broods. Biology Letters 7, 944–946 (2011). 10.1098/rsbl.2011.0391

103 Jimeno, B., Hau, M. & Verhulst, S. Corticosterone levels reflect variation in metabolic rate, independent of ‘stress’. Scientific Reports 8, 13020 (2018). 10.1038/s41598-018-31258-z

104 Girard, I. & Garland, T., Jr. Plasma corticosterone response to acute and chronic voluntary exercise in female house mice. J Appl Physiol (1985) 92, 1553–1561 (2002). 10.1152/japplphysiol.00465.2001

105 Summers, C. H. et al. Glucocorticoid interaction with aggression in non-mammalian vertebrates: Reciprocal action. European Journal of Pharmacology 526, 21–35 (2005). 10.1016/j.ejphar.2005.09.059

106 Tokarz, R. R. Effects of corticosterone treatment on male aggressive behavior in a lizard (Anolis sagrei). Hormones and Behavior 21, 358–370 (1987). 10.1016/0018-506X(87)90020-1

107 Meddle, S. L. et al. Steroid Hormone Interrelationships with Territorial Aggression in an Arctic-Breeding Songbird, Gambel’s White-Crowned Sparrow, Zonotrichia leucophrys gambelii. Hormones and Behavior 42, 212–221 (2002). 10.1006/hbeh.2002.1813

108 Haller, J. Glucocorticoids and Aggression: A Tripartite Interaction. Current topics in behavioral neurosciences 54, 209–243 (2022). 10.1007/7854_2022_307

109 Ely, D. L. & Henry, J. P. Neuroendocrine response patterns in dominant and subordinate mice. Horm Behav 10, 156–169 (1978). 10.1016/0018-506x(78)90005-3

110 Williamson, C. M., Lee, W., Romeo, R. D. & Curley, J. P. Social context-dependent relationships between mouse dominance rank and plasma hormone levels. Physiology & behavior 171, 110–119 (2017). 10.1016/j.physbeh.2016.12.038

111 Sherman, G. D. & Mehta, P. H. Stress, cortisol, and social hierarchy. Current Opinion in Psychology 33, 227–232 (2020). 10.1016/j.copsyc.2019.09.013

112 Trigo, S., Saldanha, B. C., Oliveira, P., Silva, P. A. & Soares, M. C. Corticosterone effects on aggression in a passerine species, the common waxbill Estrilda astrild. Physiology & Behavior 308, 115264 (2026). 10.1016/j.physbeh.2026.115264

113 Haller, J., Barna, I. & Baranyi, M. Hormonal and metabolic responses during psychosocial stimulation in aggressive and nonaggressive rats. Psychoneuroendocrinology 20, 65–74 (1995). 10.1016/0306-4530(94)E0042-8

114 Kruk, M. R., Halász, J., Meelis, W. & Haller, J. Fast Positive Feedback Between the Adrenocortical Stress Response and a Brain Mechanism Involved in Aggressive Behavior. Behavioral Neuroscience 118, 1062–1070 (2004). 10.1037/0735-7044.118.5.1062

115 Mikics, É., Kruk, M. R. & Haller, J. Genomic and non-genomic effects of glucocorticoids on aggressive behavior in male rats. Psychoneuroendocrinology 29, 618–635 (2004). 10.1016/S0306-4530(03)00090-8

116 Kim, J. et al. Rapid, biphasic CRF neuronal responses encode positive and negative valence. Nature Neuroscience 22, 576–585 (2019). 10.1038/s41593-019-0342-2

117 Kurischko, A. & Oettel, M. Androgen-dependent fighting behaviour in male mice. Endokrinologie 70, 1–5 (1977).

118 Sagazio, A., Shohreh, R. & Salvatori, R. Effects of GH deficiency and GH replacement on inter-male aggressiveness in mice. Growth hormone & IGF research : official journal of the Growth Hormone Research Society and the International IGF Research Society 21, 76–80 (2011). 10.1016/j.ghir.2011.01.002

119 Takao, K. & Miyakawa, T. Light/dark transition test for mice. J Vis Exp, 104 (2006). 10.3791/104

120 Kraeuter, A. K., Guest, P. C. & Sarnyai, Z. The Open Field Test for Measuring Locomotor Activity and Anxiety-Like Behavior. Methods Mol Biol 1916, 99–103 (2019). 10.1007/978-1-4939-8994-2_9

121 Mathis, A. et al. DeepLabCut: markerless pose estimation of user-defined body parts with deep learning. Nature Neuroscience 21, 1281–1289 (2018). 10.1038/s41593-018-0209-y

122 Meeks, J. P. & Mennerick, S. Action Potential Initiation and Propagation in CA3 Pyramidal Axons. Journal of Neurophysiology 97, 3460–3472 (2007). 10.1152/jn.01288.2006

