## Supplementary figure for "Hypothalamic Representation of Aggressiveness across Mouse Strains"

### Supplementary Figure 1

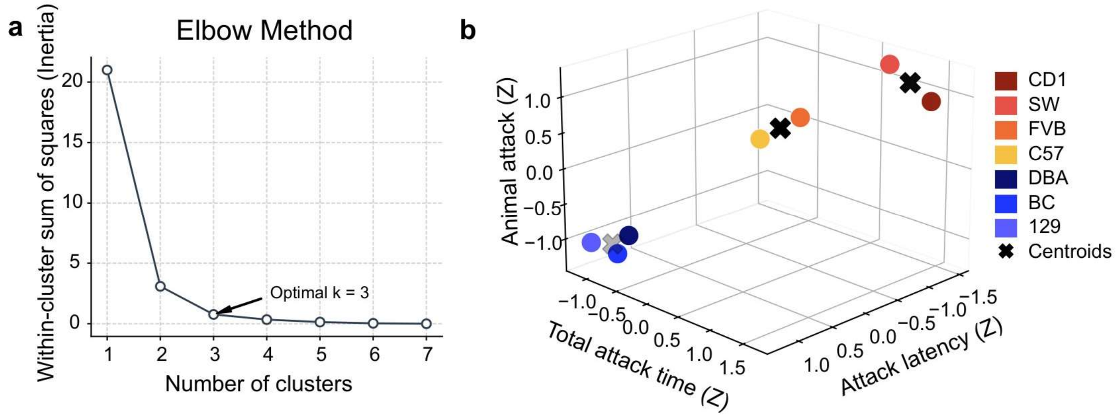


**Supplementary Figure 1. Determine the optimal number of clusters among 7 mouse strains based on their aggressive behaviors.**

**(a)** Elbow analysis of within-cluster sum of squares (WCSS) identifies three clusters as the optimal number. **(b)** Scatter plots showing z-scored aggressive behavioral metrics, including percentage of animals that attacked, attack latency, and total attack duration, across 7 strains. K-mean clustering identifies three clusters. Crosses denote cluster centroids.

Circles in (**b**) represent strain mean. See **Supplementary Table 1** for additional statistical details.

### Supplementary Figure 2

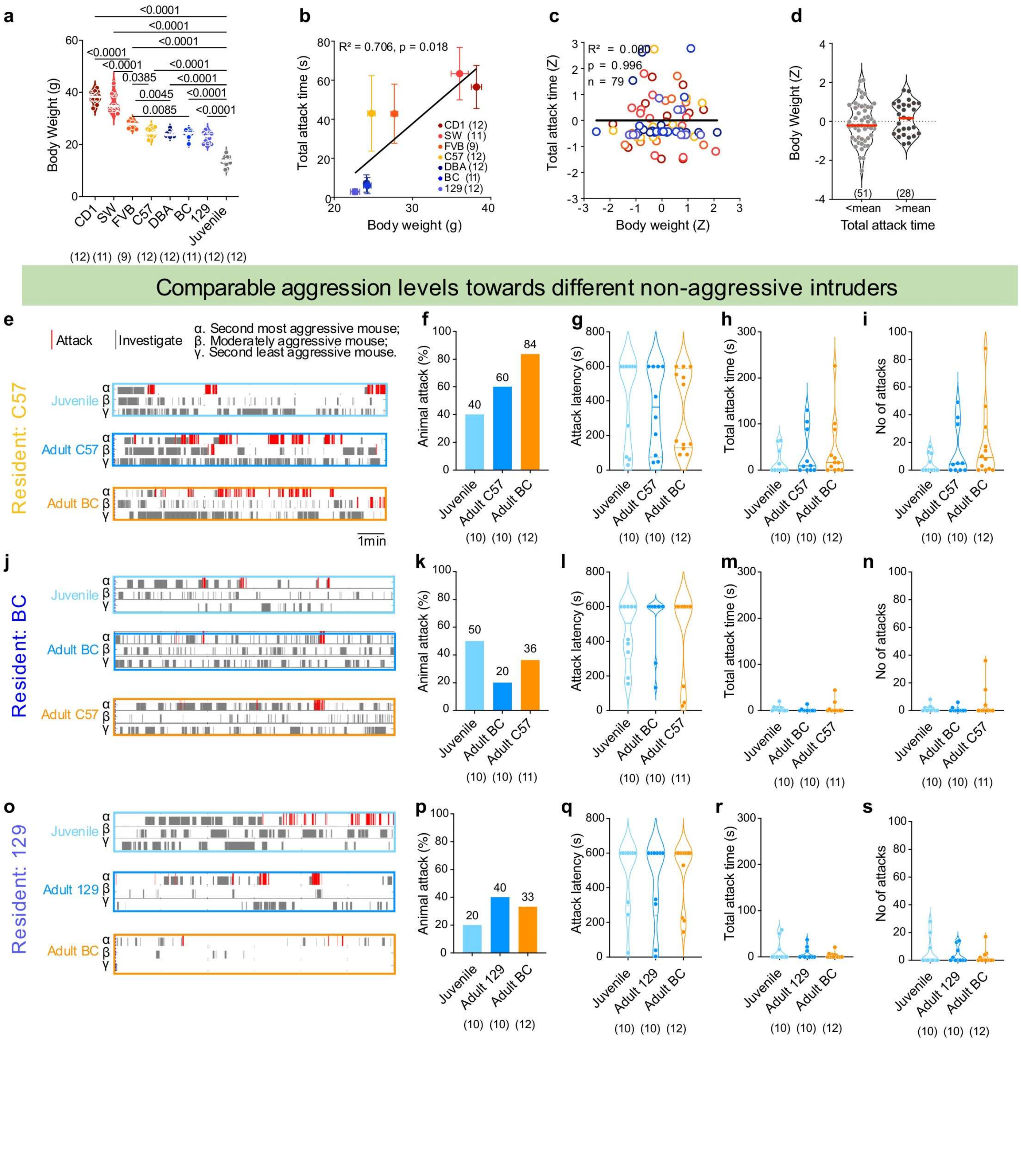

**Supplementary Figure 2. The resident's aggression level does not change based on intruder's body weight.**

**(a)** Body weight across seven strains and juvenile BC intruders.
**(b)** Correlation between average body weight and total attack duration across strains.
**(c)** Correlation between strain-normalized body weight and attack duration across individuals.
**(d)** Comparison of strain-normalized body weight between animals with attack duration below and above the strain mean.
**(e, j, o)** Representative raster plots showing attack (red) and investigation (gray) events for resident C57 **(e)**, BC **(j)**, or 129 **(o)** mice exposed to juvenile, adult same strain intruders, or adult different strain intruders. Three representative individuals are shown per condition: second most aggressive (α), moderately aggressive (β), and second least aggressive (γ).
**(f–i)** Quantification of aggressive behaviors of resident C57 mice against different intruder types, including **(f)** percentage of animals that attacked, **(g)** attack latency, **(h)** total attack time, and **(i)** attack frequency. **(k–n)** Quantification of aggressive behaviors of resident BC mice against different intruder types.
**(p–s)** Quantification of aggressive behaviors of resident 129 mice against different intruder types.

If animal shows no attack, 600s is used as the latency value. Numbers in parentheses indicate animal numbers. Color in (**a-c**) indicates strain identity, while color in (**f–l, k-n, p-s**) indicates different intruders. Circles represent data of individual animals. Solid line in (**a, d, g-i, l-n, q-s**) represents the median for each group, while dashed lines indicate quartiles. **(f, k, p)** Fisher's exact test; **(a, g-i, l-n, q-s)** One-way ANOVA for normally distributed datasets or Kruskal–Wallis test for non-normally distributed datasets with FDR-corrected post hoc comparisons (Benjamini–Krieger–Yekutieli method); **(d)** Unpaired t test. All statistical tests are two-tailed. Exact p- or q-value is shown if ≤ 0.05. Otherwise, p- or q-value is unspecified. See **Supplementary Table 1** for additional statistical details.

### Supplementary Figure 3

#### SW and C57 female mice show different levels of aggression

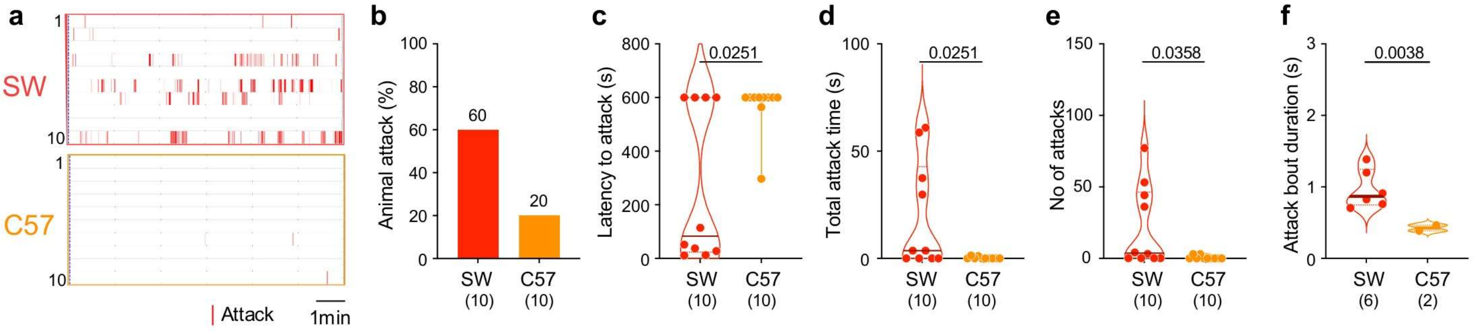


#### SW and C57 female mice show different levels of anxiety

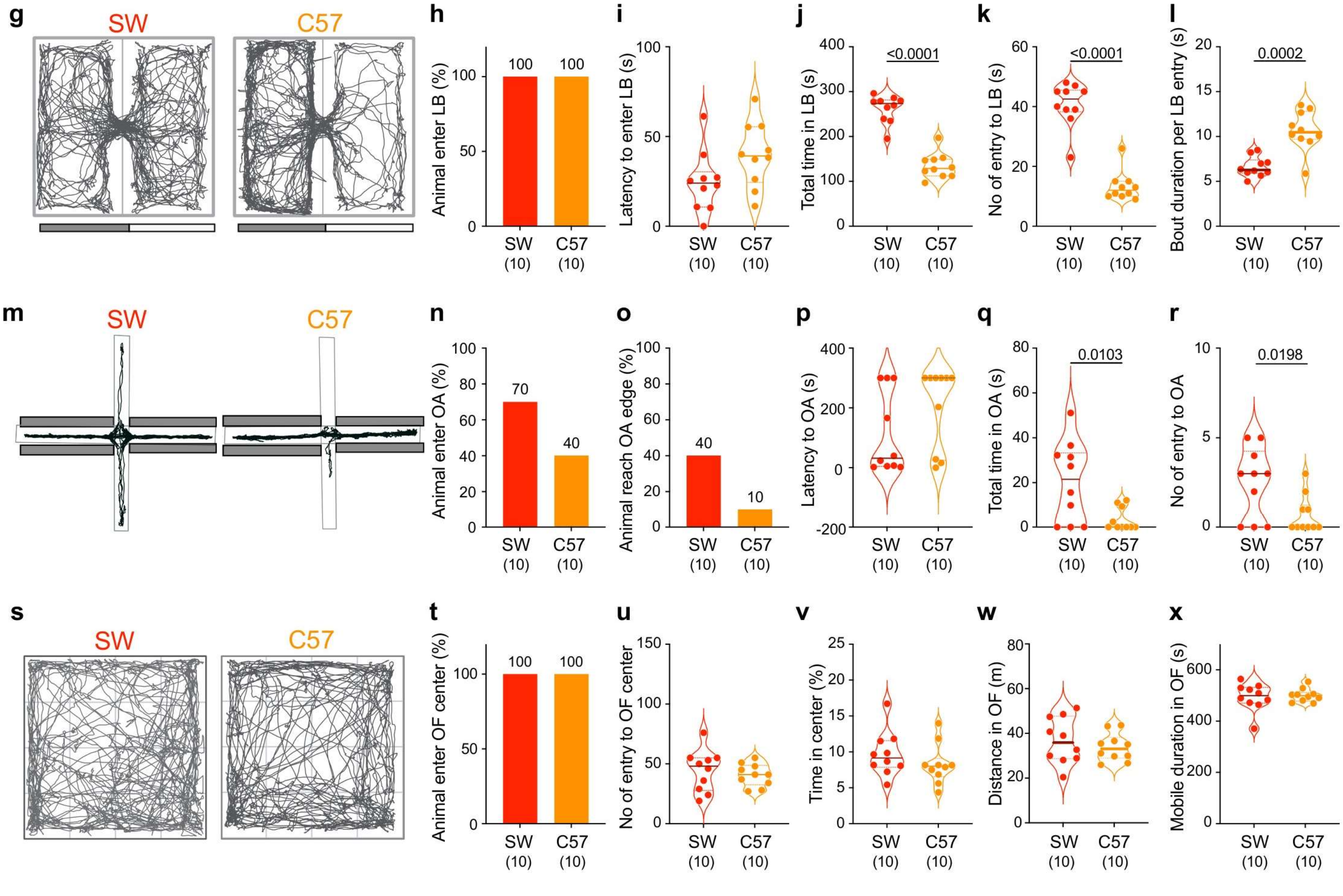


#### Relationship between aggression and anxiety levels in female mice

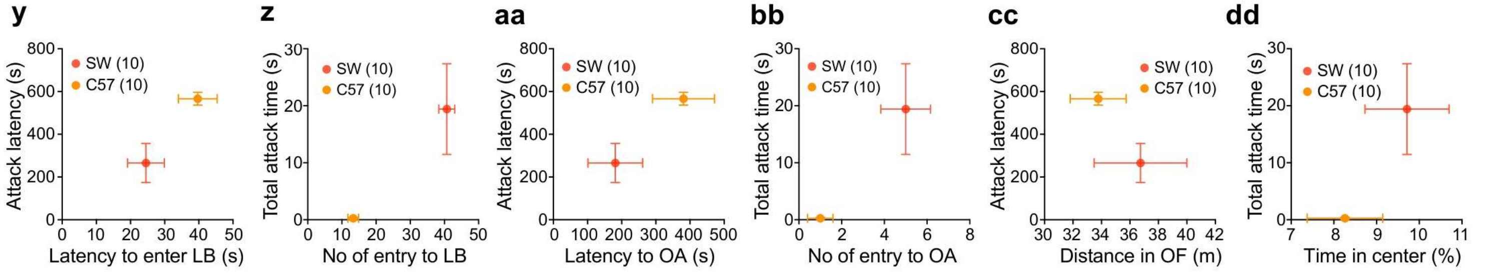

**Supplementary Figure 3. Aggression phenotypes differ between SW and C57 females**

**(a)** Raster plots showing attack events (red) for SW and C57 female resident mice. Each row represents one animal.
**(b–f)** Quantification of aggressive behaviors of SW and C57 females: **(b)** percentage of animals that attacked, **(c)** attack latency, **(d)** attack duration, **(e)** number of attacks, and **(f)** attack bout duration.
**(g)** Representative trajectories of SW and C57 females in the light–dark box (LDB).
**(h–l)** Quantification of LDB performance of SW and C57 females, including **(h)** percentage of animals entering the light box, **(i)** latency to the first entry into light box, **(j)** number of entries into light box, **(k)** total time spent in light box, and **(l)** bout duration per light box entry.
**(m)** Representative trajectories in the elevated plus maze (EPM).
**(n–r)** Quantification of EPM performance of SW and C57 females, including **(n)** percentage of animals entering open arms, **(o)** percentage of animals reaching the distal open-arm region (edge), **(p)** total time spent in open arms, **(q)** mean duration in open arms, and **(r)** number of entries into open arms.
**(s)** Representative trajectories in the open field test (OFT).
**(t–x)** Quantification of OFT performance of SW and 57 females, including **(t)** percentage of animals entering the center, **(u)** number of center entries, **(v)** percentage of time spent in the center, **(w)** distance travelled in the open field, and **(x)** mobile duration in the open field.
**(y–dd)** Plots showing aggression- and anxiety-related measures of SW and C57 females across LDB, OF and EPM tests. Each point represents the mean ± SEM across all individuals for each strain.

If animal shows no attack, 600s is used as the latency value. If animal does not enter OA, 300s is used as the latency value. Numbers in parentheses indicate animal numbers. Color indicates strain identity. Circles represent data of individual animals. Solid line in (**c-f, i-l, p-r, u-x**) represents the median for each group, while dashed lines indicate quartiles. Solid circles represent the strain mean, and error bars represent ± SEM in (**y-dd**). **(b, h, n, o, t)** Fisher's exact test; **(c-f, i-l, p-r, u-x)** Unpaired t test for normally distributed datasets or Mann–Whitney U test for non-normally distributed datasets. All statistical tests are two-tailed. Exact p- or q-value is shown if ≤ 0.05. Otherwise, p- or q-value is unspecified. See **Supplementary Table 1** for additional statistical details.

### Supplementary Figure 4

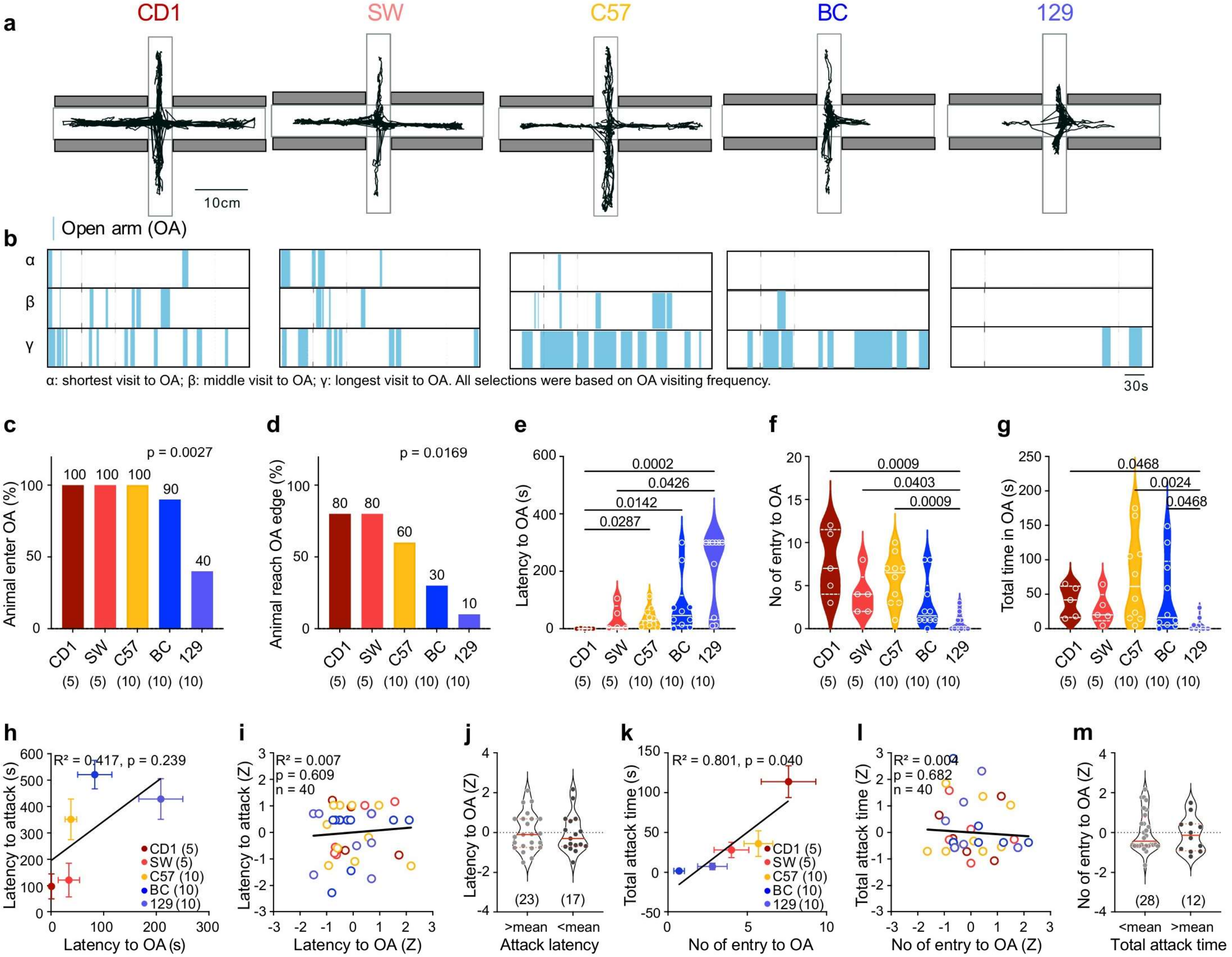


**Supplementary Figure 4. Elevated plus maze behavior differs across mouse strains correlates with aggression.**

**(a)** Representative trajectories of mice from five strains (CD1, SW, C57, BC, 129) in the elevated plus maze. Closed arms are indicated by dark gray bars.
**(b)** Raster plots showing time spent in open arms (blue) for representative animals from each strain. OA: open arm.
**(c)** Percentage of animals that entered the open arms.
**(d)** Percentage of animals that reached the far end of open-arms.
**(e–g)** Quantification of open-arm behaviors, including **(e)** latency to first open-arm entry, **(f)** number of open-arm entries, and **(g)** total time spent in open arms.
**(h)** Correlation between latency to attack and latency to enter open arm at the strain level.
**(i)** Correlation between latency to attack and latency to enter open arm at the individual level normalized by the strain mean.
**(j)** Comparison of latency to enter open arm between animals with below-mean and above-mean attack latency.
**(k)** Positive correlation between total attack duration and number of open-arm entries at the strain level.
**(l)** No significant correlation between total attack duration and number of open-arm entries at the individual level.
**(m)** Comparison of open-arm entry counts between animals with below-mean and above-mean attack duration.

If animal shows no attack, 600s is used as the latency value. If animal does not enter OA, 300s is used as the latency value. Numbers in parentheses indicate animal numbers. Color in (**c-i and k-l**) indicates strain identity. Circles in **(e-g, i-j, l-m**) represent data of individual animals. Solid line in (**e-g, j, m**) represents the median for each group, while dashed lines indicate quartiles. Solid circles represent the strain mean, and error bars represent ± SEM in (**h, k**). **(c, d)** Fisher's exact test; **(e-g)** One-way ANOVA for normally distributed datasets or Kruskal–Wallis test for non-normally distributed datasets with FDR-corrected post hoc comparisons (Benjamini–Krieger–Yekutieli method); **(h-i, k-l)** Linear regression and Pearson correlation with reported $R^2$ and *p*-values; **(j, m)** Unpaired t test for normally distributed datasets or Mann–Whitney U test for non-normally distributed datasets. All statistical tests are two-tailed. Exact p- or q-value is shown if ≤ 0.05. Otherwise, p- or q-value is unspecified. See **Supplementary Table 1** for additional statistical details.

### Supplementary Figure 5

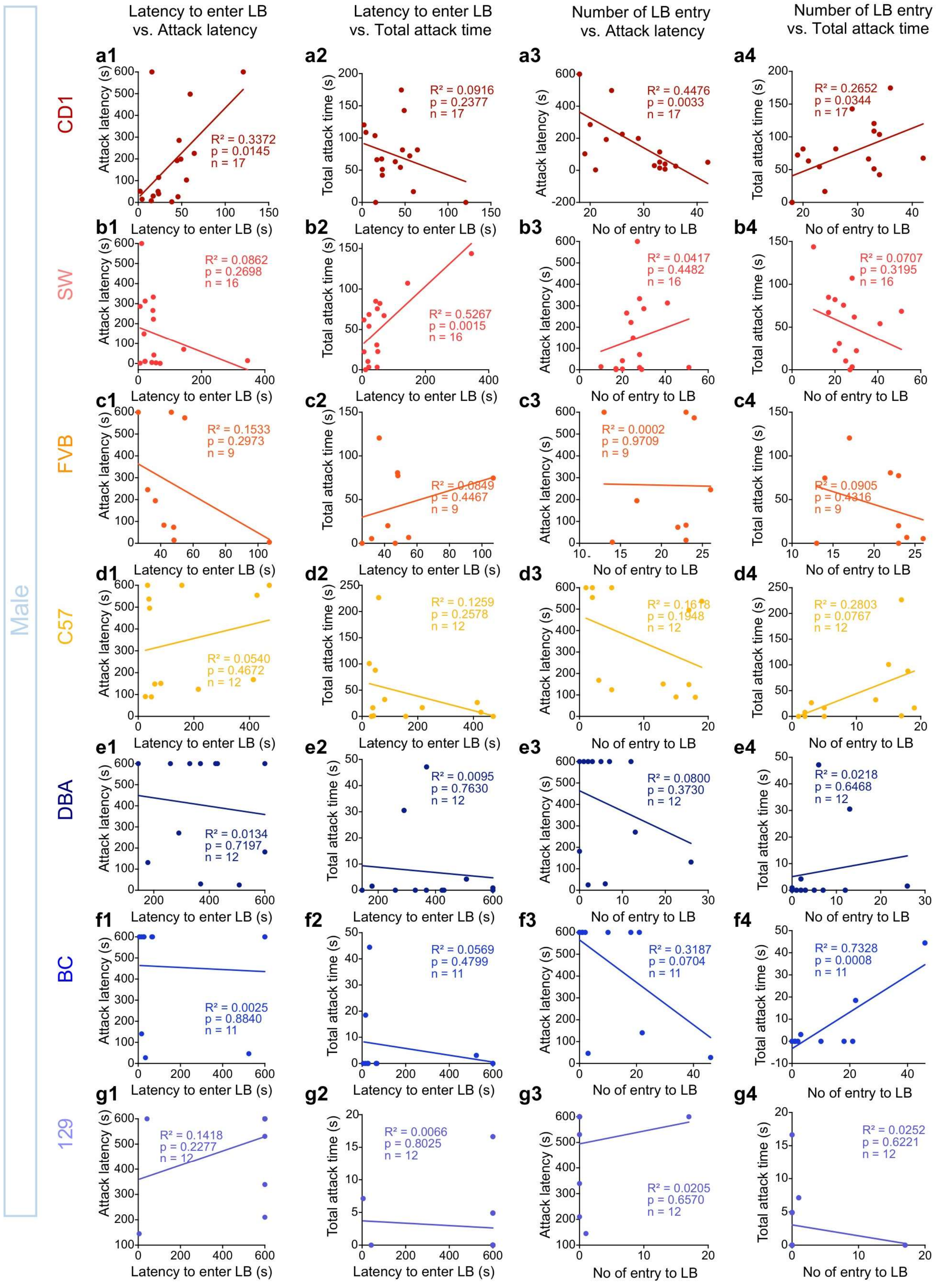

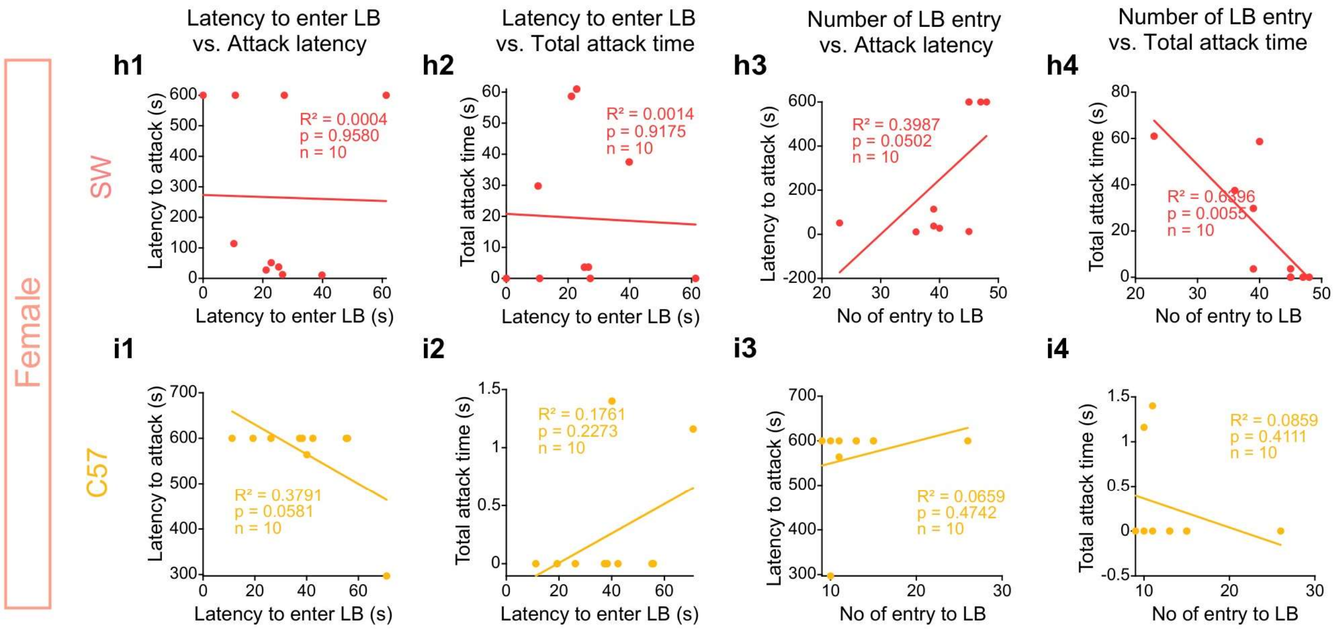


**Supplementary Figure 5. Correlation between light-dark box performance and innate aggressiveness across individuals of each strain.**

**(a1–i1)** Correlations between latency to enter the light compartment in the light–dark box test and attack latency. Data are shown for male mice of the following strains: **(a1)** CD1, **(b1)** SW, **(c1)** FVB, **(d1)** C57, **(e1)** DBA, **(f1)** BC, and **(g1)** 129; and for female mice: **(h1)** SW and **(i1)** C57. LB, light box.
**(a2-i2)** Correlations between latency to enter the light box and total attack time. Data are shown for male mice of the following strains: **(a2)** CD1, **(b2)** SW, **(c2)** FVB, **(d2)** C57, **(e2)** DBA, **(f2)** BC, and **(g2)** 129; and for female mice: **(h2)** SW and **(i2)** C57.
**(a3-i3)** Correlations between number of light box entries and attack latency. Data are shown for male mice of the following strains: **(a3)** CD1, **(b3)** SW, **(c3)** FVB, **(d3)** C57, **(e3)** DBA, **(f3)** BC, and **(g3)** 129; and for female mice: **(h3)** SW and **(i3)** C57.
**(a4-i4)** Correlations between number of light box entries and total attack time and. Data are shown for male mice of the following strains: **(a4)** CD1, **(b4)** SW, **(c4)** FVB, **(d4)** C57, **(e4)** DBA, **(f4)** BC, and **(g4)** 129; and for female mice: **(h4)** SW and **(i4)** C57.

If animal shows no attack or does not enter LB, 600s is used as the latency value. Circles represent data of individual animals and color indicates strain identity. **(a-i)** Pearson correlation with reported $R^2$, *p*-values, and animal numbers. Colored lines represent the linear regression lines. See **Supplementary Table 1** for additional statistical details.

### Supplementary Figure 6

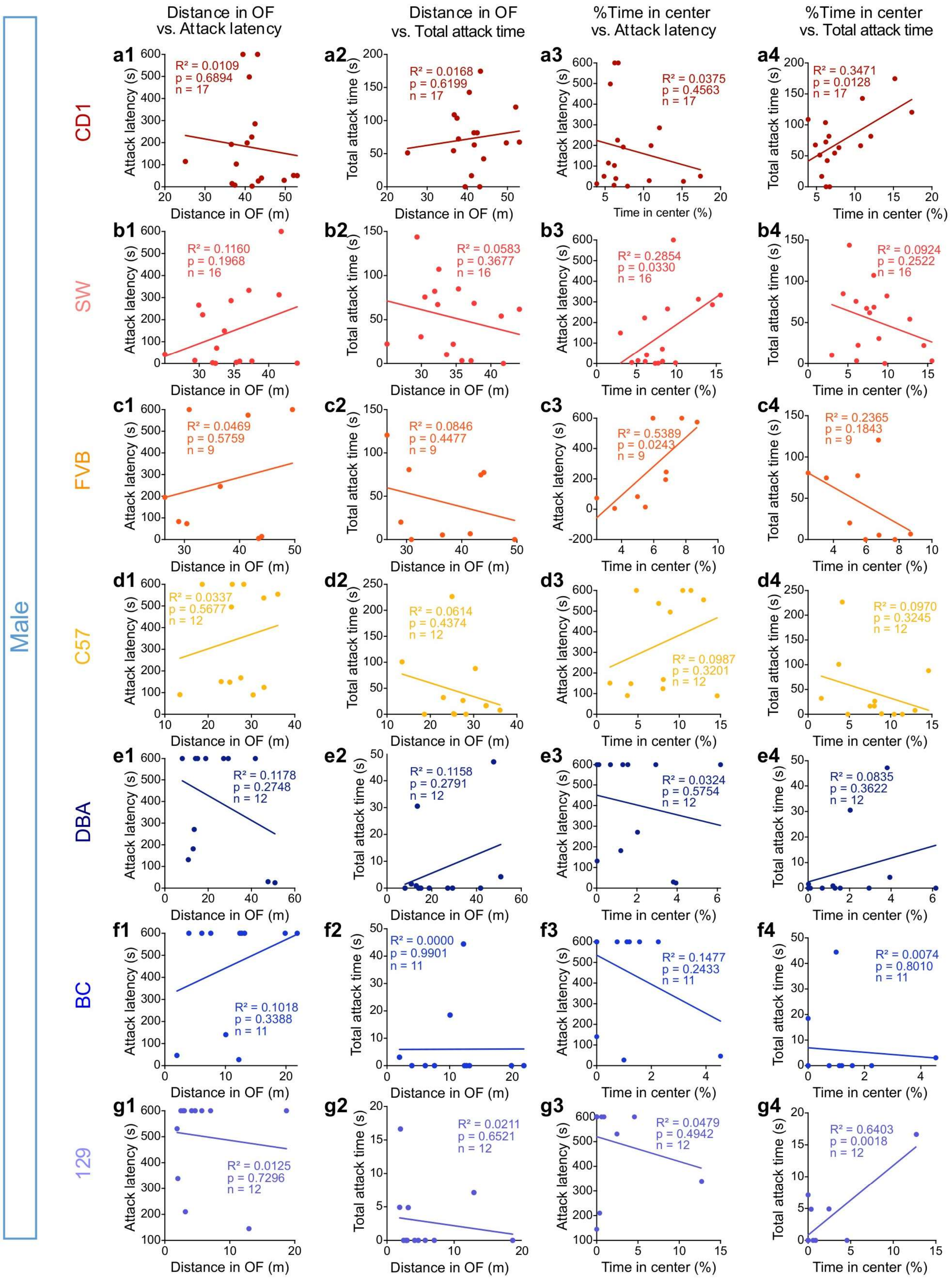

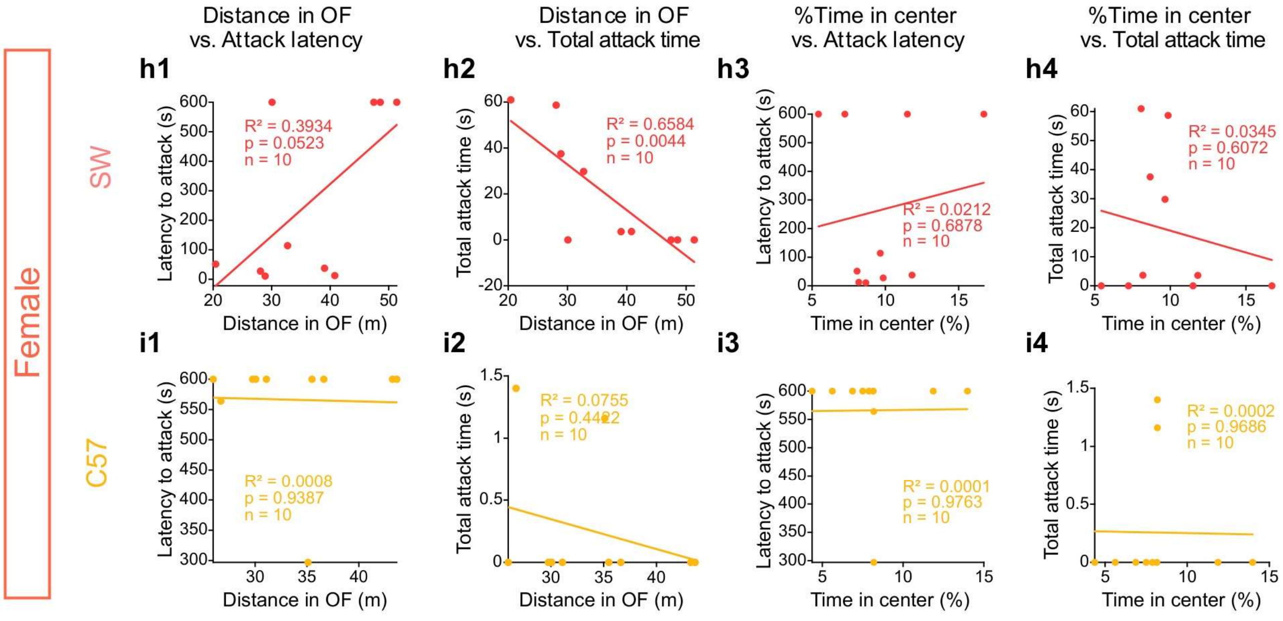


**Supplementary Figure 6. Correlation between open field test performance and innate aggressiveness across individuals of each strain.**

**(a1-i1)** Correlations between distance traveled in the in open field during a 10-minute test and attack latency. Data are shown for male mice of the following strains: **(a1)** CD1, **(b1)** SW, **(c1)** FVB, **(d1)** C57, **(e1)** DBA, **(f1)** BC, and **(g1)** 129; and for female mice: **(h1)** SW and **(i1)** C57.
**(a2-i2)** Correlations between and distance traveled in the in open field and total attack time. Data are shown for male mice of the following strains: **(a2)** CD1, **(b2)** SW, **(c2)** FVB, **(d2)** C57, **(e2)** DBA, **(f2)** BC, and **(g2)** 129; and for female mice: **(h2)** SW and **(i2)** C57.
**(a3-i3)** Correlations between percentage of time in the open field center and attack latency. Data are shown for male mice of the following strains: **(a3)** CD1, **(b3)** SW, **(c3)** FVB, **(d3)** C57, **(e3)** DBA, **(f3)** BC, and **(g3)** 129; and for female mice: **(h3)** SW and **(i3)** C57.
**(a4-i4)** Correlations between percentage of time in the open field center and total attack time. Data are shown for male mice of the following strains: **(a4)** CD1, **(b4)** SW, **(c4)** FVB, **(d4)** C57, **(e4)** DBA, **(f4)** BC, and **(g4)** 129; and for female mice: **(h4)** SW and **(i4)** C57.

If animal shows no attack, 600s is used as the latency value. Circles represent data of individual animals and color indicates strain identity. **(a-i)** Pearson correlation with reported $R^2$, *p*-values and animal numbers. Colored lines represent the linear regression lines. See **Supplementary Table 1** for additional statistical details.

Supplementary Figure 7

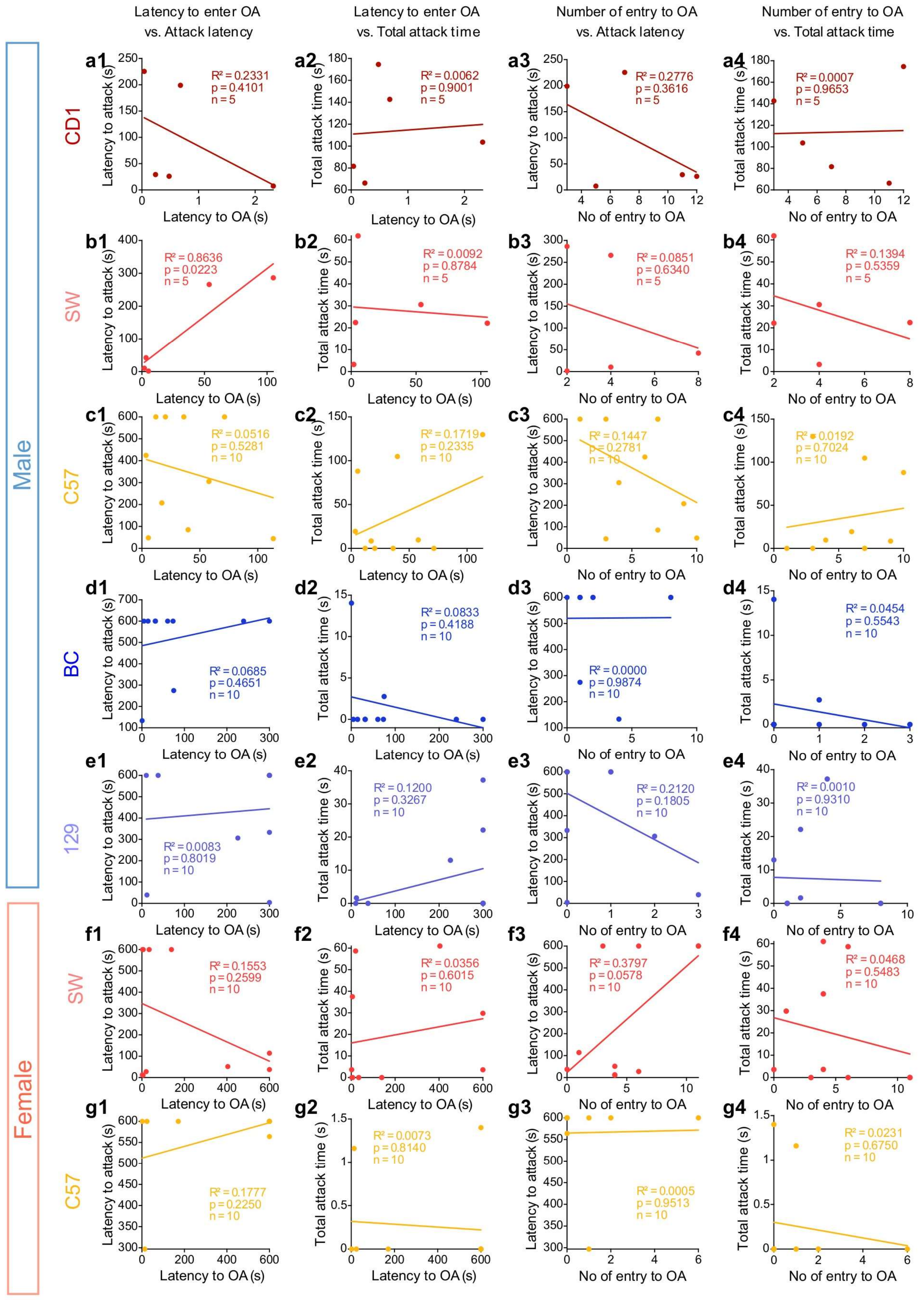

**Supplementary Figure 7. Correlation between elevated-plus maze performance and innate aggressiveness across individuals of each strain.**

**(a1-g1)** Correlations between latency to enter the open arm during a 10-minute elevated-plus maze test and attack latency. Data are shown for male mice of the following strains: **(a1)** CD1, **(b1)** SW, **(c1)** C57, **(d1)** BC, **(e1)** 129; and for female mice: **(f1)** SW and **(g1)** C57. OA: open arm.
**(a2-g2)** Correlations between latency to enter the open arm and total attack time and. Data are shown for male mice of the following strains: **(a2)** CD1, **(b2)** SW, **(c2)** C57, **(d2)** BC, **(e2)** 129; and for female mice: **(f2)** SW and **(g2)** C57.
**(a3-g3)** Correlations between number of entries into the open arm and attack latency. Data are shown for male mice of the following strains: **(a3)** CD1, **(b3)** SW, **(c3)** C57, **(d3)** BC, **(e3)** 129; and for female mice: **(f3)** SW and **(g3)** C57.
**(a4-g4)** Correlations between number of entries into the open arm and total attack time. Data are shown for male mice of the following strains: **(a4)** CD1, **(b4)** SW, **(c4)** C57, **(d4)** BC, **(e4)** 129; and for female mice: **(f4)** SW and **(g4)** C57.

If animal shows no attack or does not enter to the OA, 600s is used as the latency value. Circles represent data of individual animals and color indicates strain identity. **(a-g)** Pearson correlation with reported $R^2$, *p*-values and animal numbers. Colored lines represent the linear regression lines. See **Supplementary Table 1** for additional statistical details.

### Supplementary Figure 8

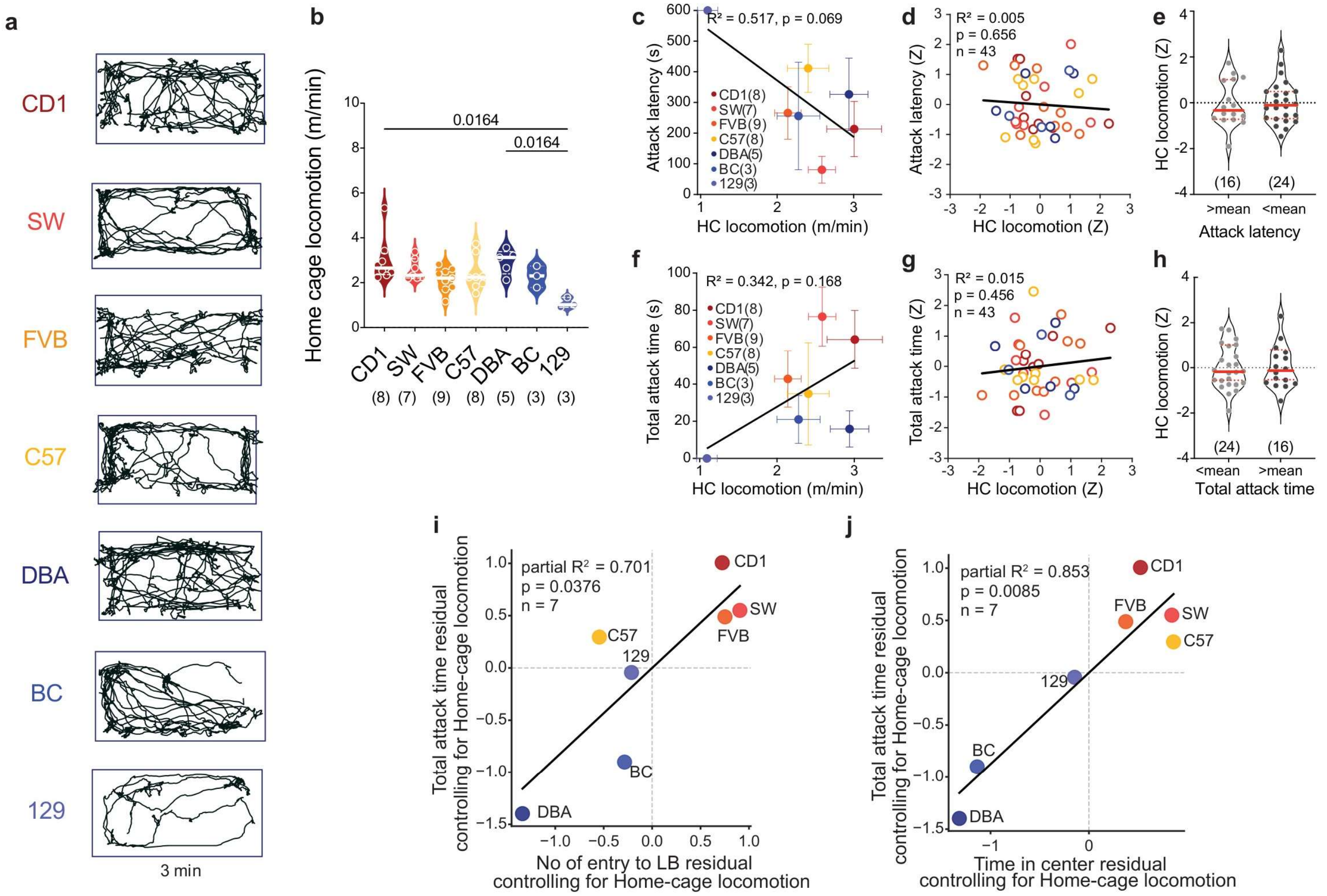


**Supplementary Figure 8. Separate the contribution of locomotion and anxiety to aggression.**

**(a)** 3-mintues home cage tracking results from representative animals of various strain.
**(b)** Average home-cage locomotion speed across strains.
**(c)** Correlation between home-cage locomotion speed and mean attack latency across strains.
**(d)** Correlation between strain-normalized home-cage locomotion speed and attack latency across individuals.
**(e)** Comparison of home-cage locomotion speed between individuals with attack latency above or below strain average.
**(f)** Correlation between home-cage locomotion speed and total attack time across strains.
**(g)** Correlation between strain-normalized and home-cage locomotion speed and total attack time across individuals.
**(h)** Comparison of home-cage locomotion speed between individuals with total attack time below or above strain average.
**(i)** Relationship between total attack duration and number of entries into the light compartment in the light–dark box, after controlling for home-cage velocity. Each point represents the mean value for one strain (n = 7 strains). Residuals were obtained by regressing each variable against home-cage velocity.
**(j)** Relationship between total attack duration and percentage of time in the open-field center, after controlling for home-cage velocity. Each point represents the mean value for one strain. Residuals were computed as in (**j**).

Numbers in parentheses indicate animal numbers. Circles represent data of individual animals (**b, d, e, g, h**) or strains (**c, f, i, j**). Solid line in (**b, e, h**) represents the median for each group, while dashed lines indicate quartiles. Color in **(b-d, f, g, i, j)** indicates strain identity. Error bars in (**c, f**) represent ± SEM. Kruskal–Wallis test with FDR correction **(b)**, unpaired t test **(e, h)**, Pearson correlation **(c, d, f, g, I, j)** with reported *R²* and *p*-values. Lines in **(c, d, f, g, I, j)** represent the linear regression lines. All statistical tests are two-tailed. Exact p- or q-values are shown if ≤ 0.05; If not indicated, p or q values > 0.05. See **Supplementary Table 1** for additional statistical details.

### Supplementary Figure 9

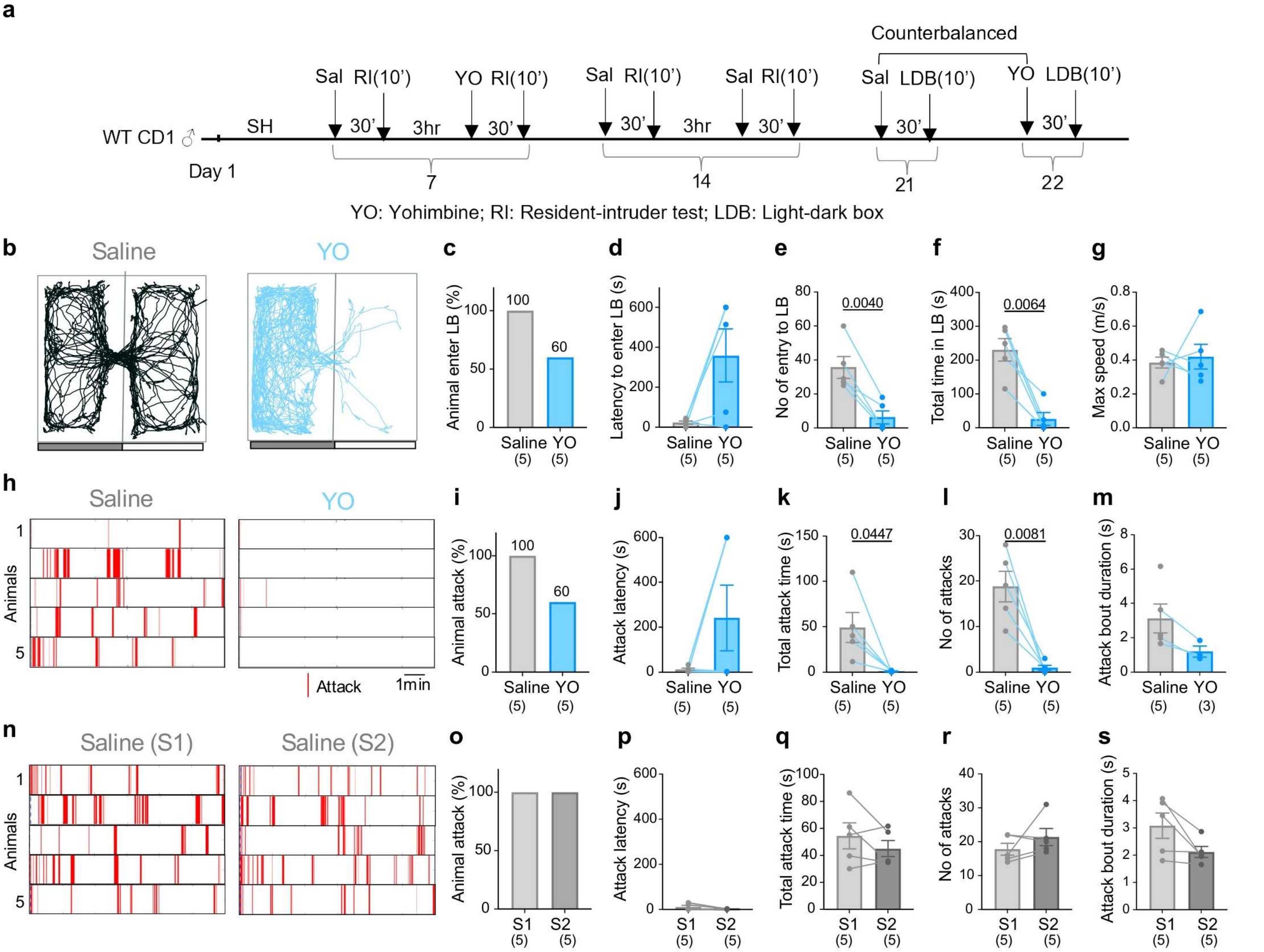


**Supplementary Figure 9. Effect of Yohimbine (YO) treatment on anxiety-like and aggression-related behaviors.**

**(a)** Experimental paradigm.
**(b)** Representative trajectories of CD1 mice in the light–dark box under saline (control) and YO treatment conditions.
**(c–g)** Quantification of LDB behavior after saline and YO injections, including **(c)** percentage of animals entering the light box, **(d)** latency to first entry into the light box, **(e)** number of entries into the light box, **(f)** total time spent in the light box, and **(g)** maximum locomotor speed. LB: light box.
**(h)** Raster plots showing attack for saline- and YO-treated CD1 mice in a 10-min test.
**(i–m)** Quantification of aggression after saline and YO injections, including **(i)** percentage of animals exhibiting attack, **(j)** attack latency, **(k)** total attack duration, **(l)** number of attacks, and **(m)** attack bout duration. If no attack, 600s is used as the latency value.
**(n)** Raster plots showing attack for saline- and YO-treated CD1 mice in a 10-min test.
**(o–s)** Quantification of aggression after saline injections (S1: session 1, S2: session 2), including **(o)** percentage of animals exhibiting attack, **(p)** attack latency, **(q)** total attack time, **(r)** number of attacks, and **(s)** attack bout duration.

Numbers in parentheses indicate animal numbers. Circles represent data of individual animals and lines represent paired measurements. Bars represent the group mean and error bars represent ± SEM in (**d-g, j-m, p-s**). **(c, i, o)** Fisher's exact test; **(d-g, j-m, p-s)** Paired t test for normally distributed datasets or Wilcoxon test for non-normally distributed datasets. All statistical tests are two-tailed. Exact p- or q-value is shown if ≤ 0.05. Otherwise, p- or q-value is unspecified. See **Supplementary Table 1** for additional statistical details.

### Supplementary Figure 10

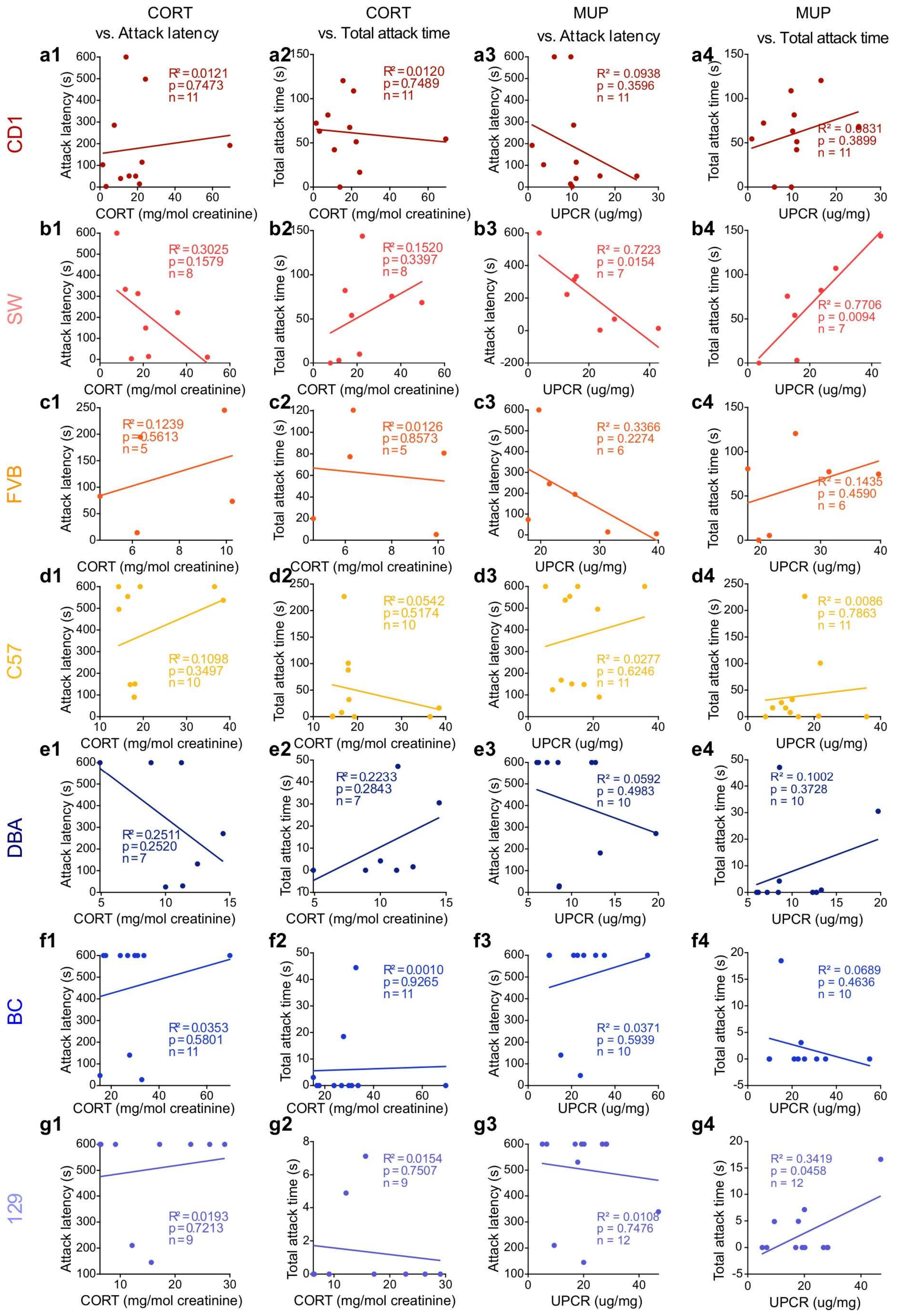

**Supplementary Figure 10. Correlation between CORT, urinary protein and aggression across individual of each strain.**

**(a1-g1)** Correlations between corticosterone concentration and attack latency of (**a1**) CD1, (**b1**) SW, (**c1**) FVB, (**d1**) C57, (**e1**) DBA, (**f1**) BC, and (**g1**) 129 male mice.
**(a2-g2)** Correlations between corticosterone concentration and total attack time of (**a2**) CD1, (**b2**) SW, (**c2**) FVB, (**d2**) C57, (**e2**) DBA, (**f2**) BC, and (**g2**) 129 male mice.
**(a3-g3)** Correlations between urinary protein concentration and attack latency of (**a3**) CD1, (**b3**) SW, (**c3**) FVB, (**d3**) C57, (**e3**) DBA, (**f3**) BC, and (**g3**) 129 male mice. UPCR: urine protein-to-creatinine ratio.
**(a4-g4)** Correlations between urinary protein concentration and total attack time of (**a4**) CD1, (**b4**) SW, (**c4**) FVB, (**d4**) C57, (**e4**) DBA, (**f4**) BC, and (**g4**) 129 male mice.

If animal shows no attack, 600s is used as the latency value. Circles represent data of individual animals and color indicates strain identity. **(a-g)** Pearson correlation with reported $R^2$, *p*-values and animal numbers. Colored lines represent the linear regression lines. See **Supplementary Table 1** for additional statistical details.

### Supplementary Figure 11

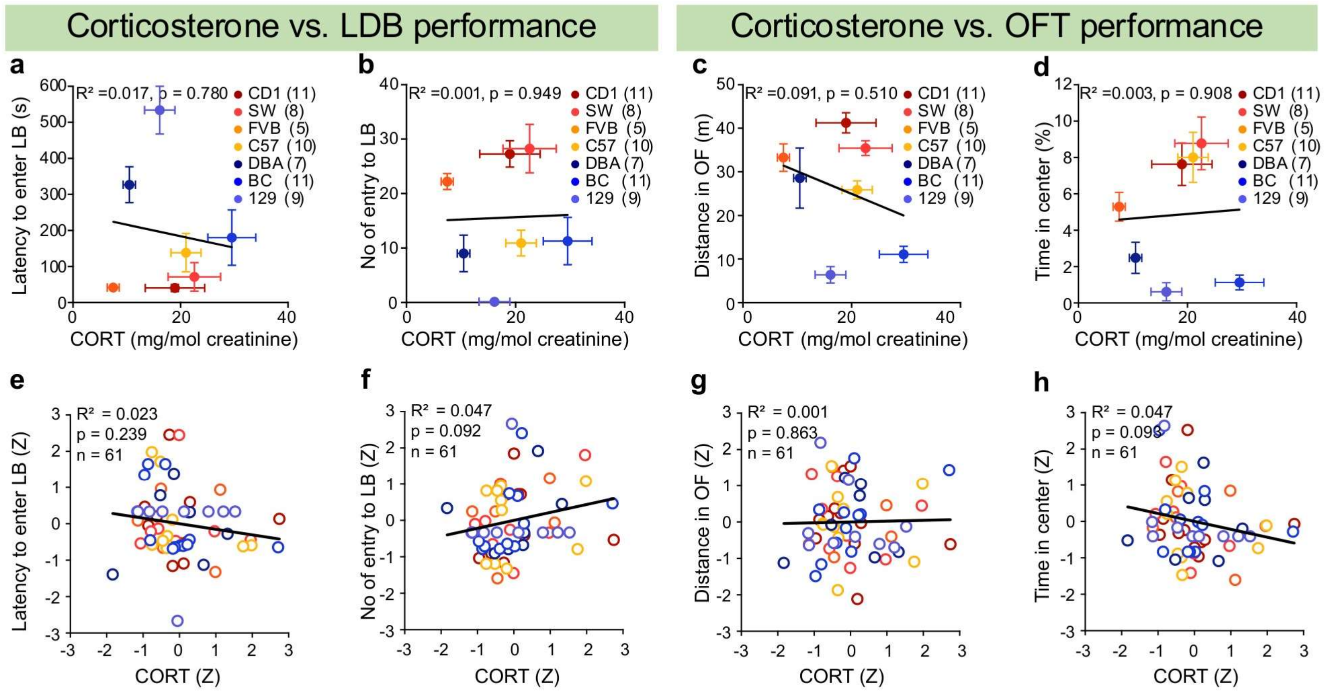


**Supplementary Figure 11. Relationship between corticosterone level and anxiety.**

**(a-b)** Correlation between average corticosterone concentration and **(a)** latency to enter light box and **(b)** number of entries to light box across strains. LB: light box.
**(c-d)** Correlation between average corticosterone concentration and **(c)** distance traveled in the open field and **(d)** percentage of time spent in the center. OF: open field.
**(e-f)** Correlation between strain-normalized corticosterone concentration and **(e)** latency to enter light box and **(f)** number of entries to light box across individuals.
**(g-h)** Correlation between strain-normalized corticosterone concentration and **(g)** distance traveled in the open field and **(h)** percentage of time spent in the center.

Color indicates strain identity. Circles and error bars in **(a-d)** represent strain mean and ±SEM. Circles in **(e-h)** represent data of individual animals. **(a-h)** Pearson correlation with reported *R²*, *p*-values and animal numbers. Lines represent the linear regression lines. See **Supplementary Table 1** for additional statistical details.

### Supplementary Figure 12

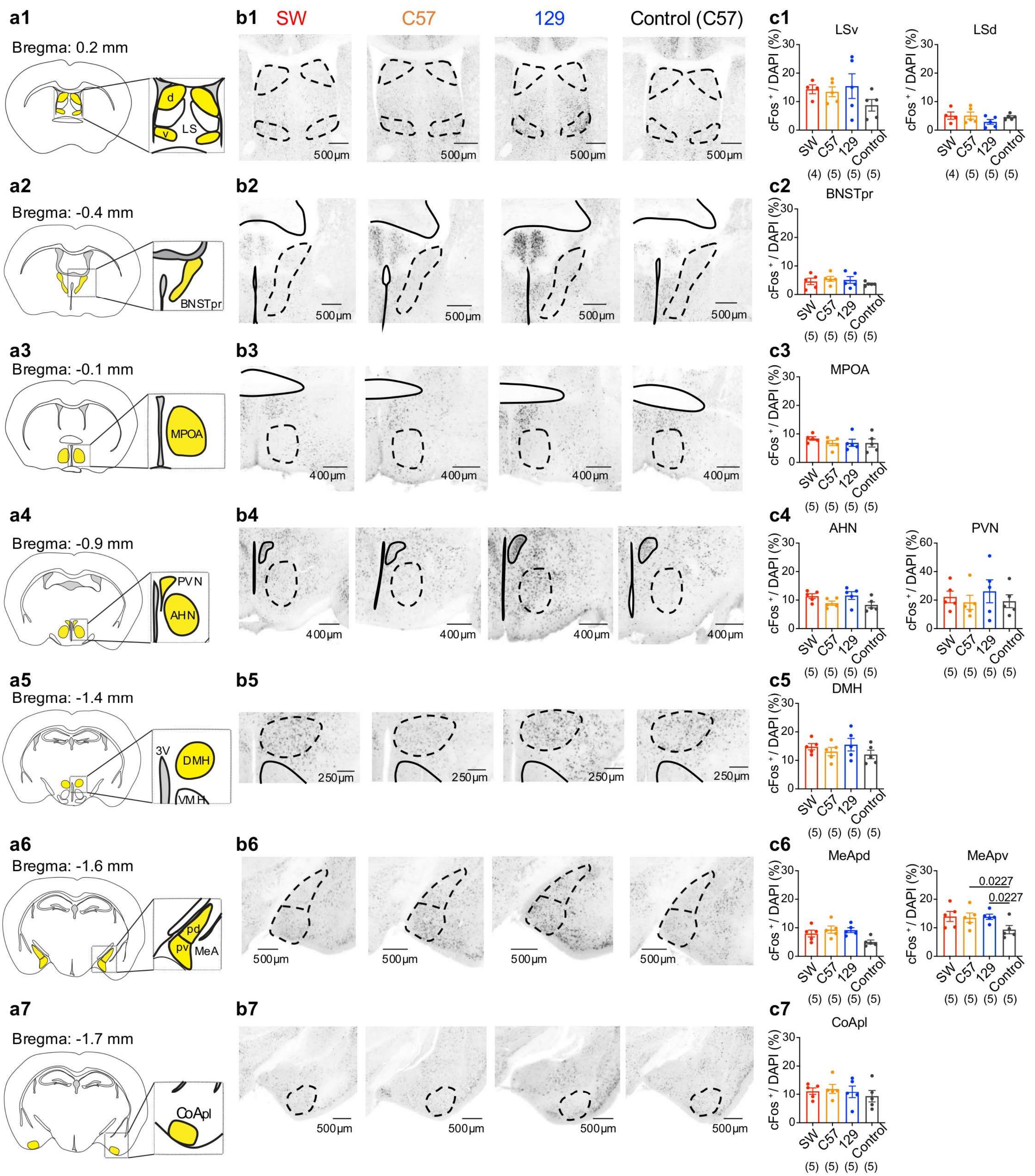

### Supplementary Figure 12

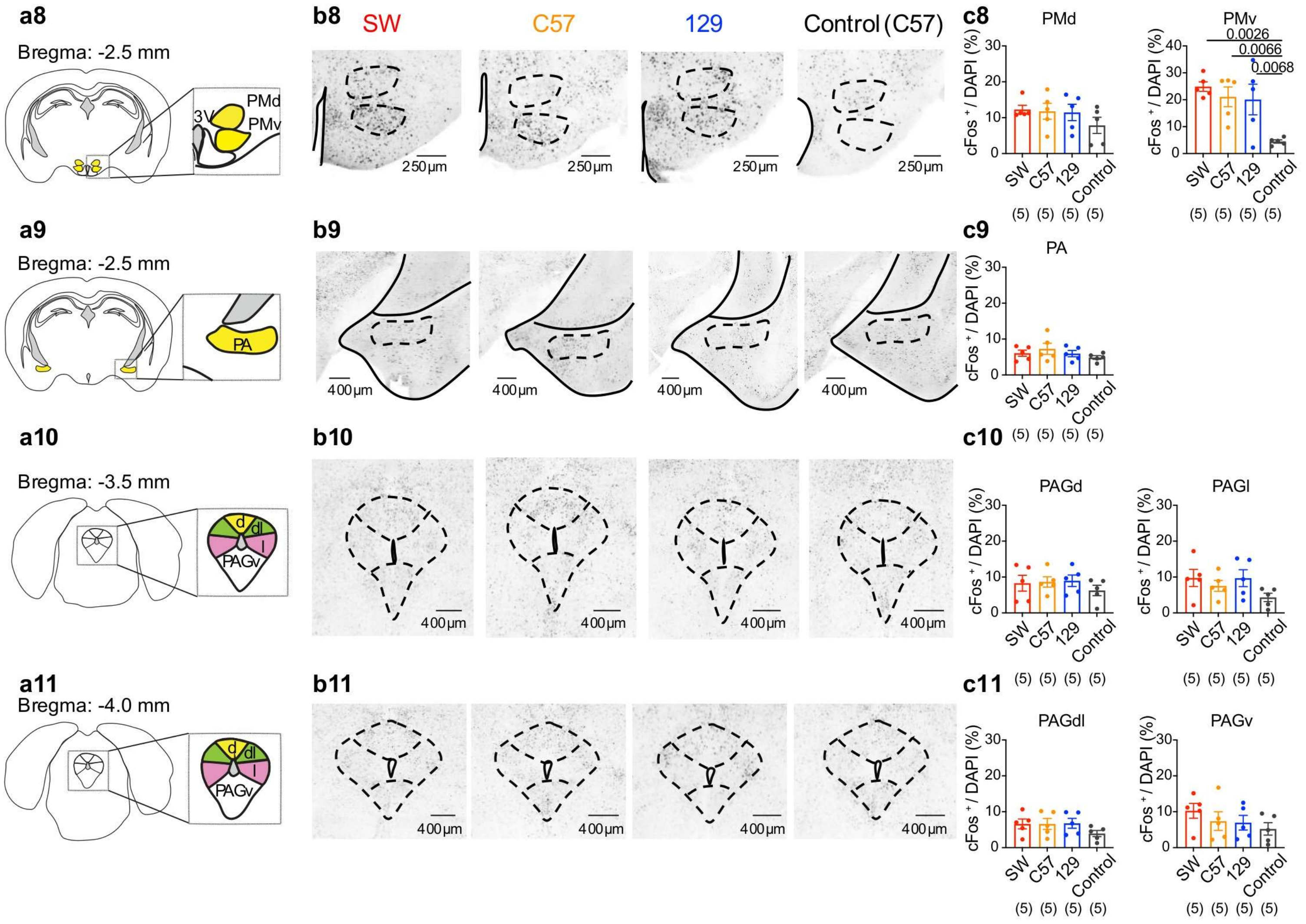


**Supplementary Figure 12. Intruder-induced c-Fos expression in the limbic system across strains.**

**(a1–a11)** Schematics of the mouse brain atlas showing analyzed regions, including **(a1)** dorsal and ventral lateral septum (LSd and LSv), **(a2)** principal nucleus of the bed nucleus of the stria terminalis (BNSTpr), **(a3)** medial preoptic area (MPOA), **(a4)** paraventricular nucleus of the hypothalamus (PVN) and anterior hypothalamic nucleus (AHN), **(a5)** dorsomedial hypothalamus (DMH), **(a6)** posterodorsal and posteroventral medial amygdala (MeApd and MeApv), **(a7)** posterolateral cortical amygdala (CoApl), **(a8)** dorsal and ventral premammillary nucleus (PMd and PMv), **(a9)** posterior amygdala (PA), **(a10-a11)** dorsal part of periaqueductal gray (PAGd), lateral part of periaqueductal gray (PAGl), dorsal-lateral part of periaqueductal gray (PAGdl), and ventral part of periaqueductal gray (PAGv).
**(b1–b11)** Representative coronal sections showing intruder-induced c-Fos expression at Bregma levels: **(b1)** 0.2 mm, **(b2)** –0.4 mm, **(b3)** –0.1 mm, **(b4)** –0.9 mm, **(b5)** –1.4 mm, **(b6)** –1.6 mm, **(b7)** –1.7 mm, **(b8)** –2.5 mm, **(b9)** –2.5 mm, **(b10)** –3.5 mm, **(b11)** –4.0 mm from SW, C57, and 129 males, and empty-cup control from a C57 male. c-Fos–positive cells appear black. Dashed lines mark the boundaries of the regions of interest as shown in **(a).**
**(c1–c11)** Quantification of c-Fos–positive cells in corresponding regions in **(a)**. n=5 animals per group.

Numbers in parentheses indicate animal numbers. Circles represent data of individual animals. Solid circles represent the strain mean, and error bars represent ± SEM. **(c1-c11)** One-way ANOVA for normally distributed datasets or Kruskal–Wallis test for non-normally distributed datasets with FDR-corrected post hoc comparisons (Benjamini–Krieger–Yekutieli method). All statistical tests are two-tailed. Exact p- or q-value is shown if ≤ 0.05. Otherwise, p- or q-value is unspecified. See **Supplementary Table 1** for additional statistical details.

### Supplementary Figure 13

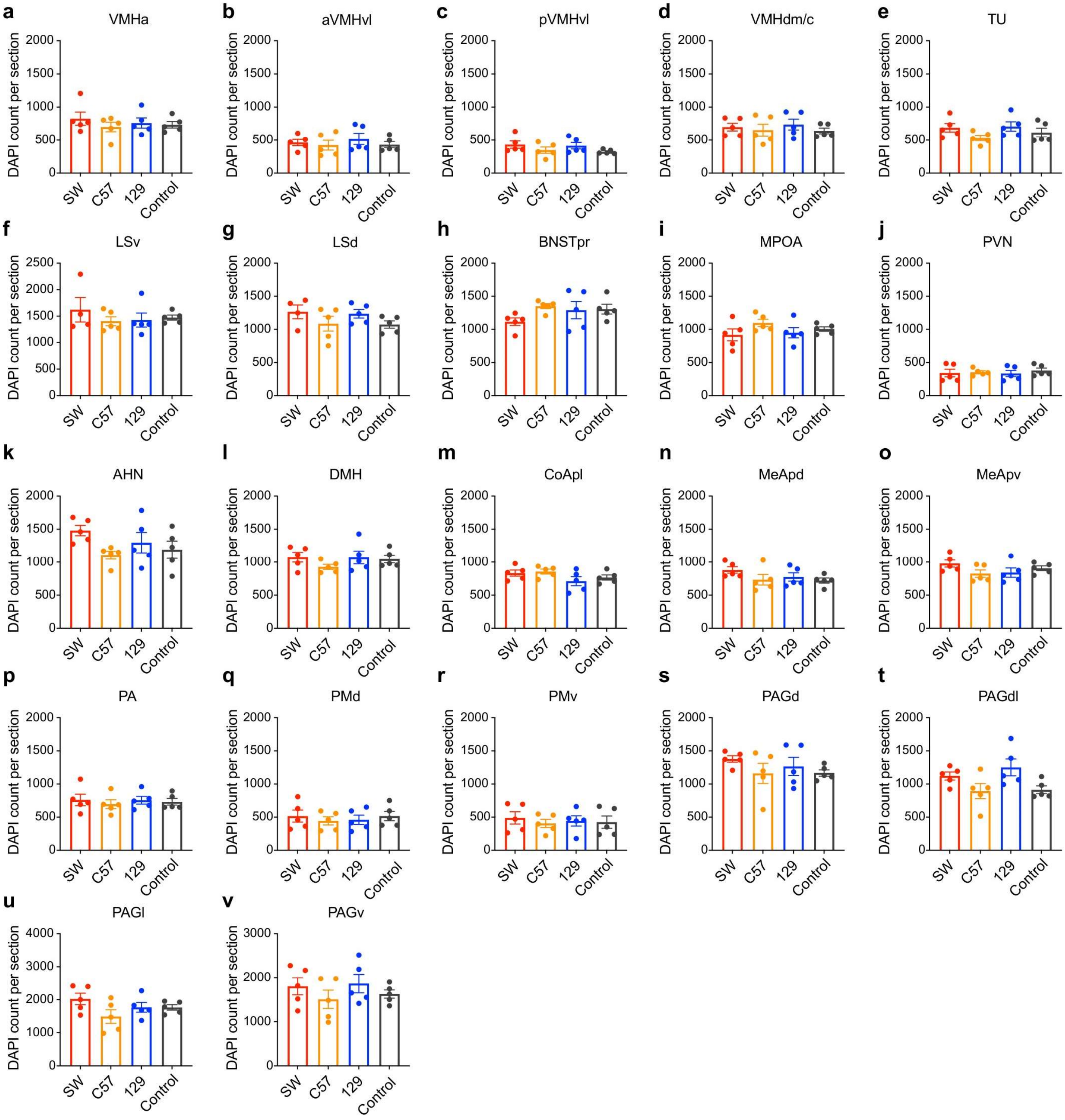


**Supplementary Figure 13. Quantification of total cells in analyzed brain regions across strains.**

**(a–v)** Quantification of DAPI-positive cells (average DAPI counts per section) across brain regions, including **(a)** anterior ventromedial hypothalamus (aVMH), **(b)** anterior ventrolateral part of the ventromedial hypothalamus (aVMHvl), (**c**) posterior VMHvl (pVMHvl), **(d)** dorsomedial/central VMH (VMHdm/c), **(e)** tuberal nucleus (TU), **(f and g)** lateral septum ventral and dorsal divisions (LSv, LSd), **(h)** principal nucleus of the bed nucleus of the stria terminalis (BNSTpr), (**i**), medial preoptic area (MPOA), **(j)** paraventricular nucleus of the hypothalamus (PVN), **(k)** anterior hypothalamic nucleus (AHN), **(l)** dorsomedial hypothalamus (DMH), **(m)** cortical amygdala posterolateral part (CoApl), **(n-o)** medial amygdala posterior dorsal and ventral divisions (MeApd, MeApv), **(p)** posterior amygdala (PA), **(q-r)** dorsal and ventral premammillary nuclei (PMd, PMv), **(s-v)** periaqueductal gray subdivisions including dorsal (PAGd; **s**), dorsolateral (PAGdl; **t**), lateral (PAGl; **u**), and ventral (PAGv; **v**).

Circles represent data of individual animals. Solid circles represent the strain mean, and error bars represent ± SEM. One-way ANOVA for normally distributed datasets or Kruskal–Wallis test for non-normally distributed datasets with FDR-corrected post hoc comparisons (Benjamini–Krieger–Yekutieli method). All statistical tests are two-tailed. Exact p- or q-value is shown if ≤ 0.05. Otherwise, p- or q-value is unspecified. See **Supplementary Table 1** for additional

### Supplementary Figure 14

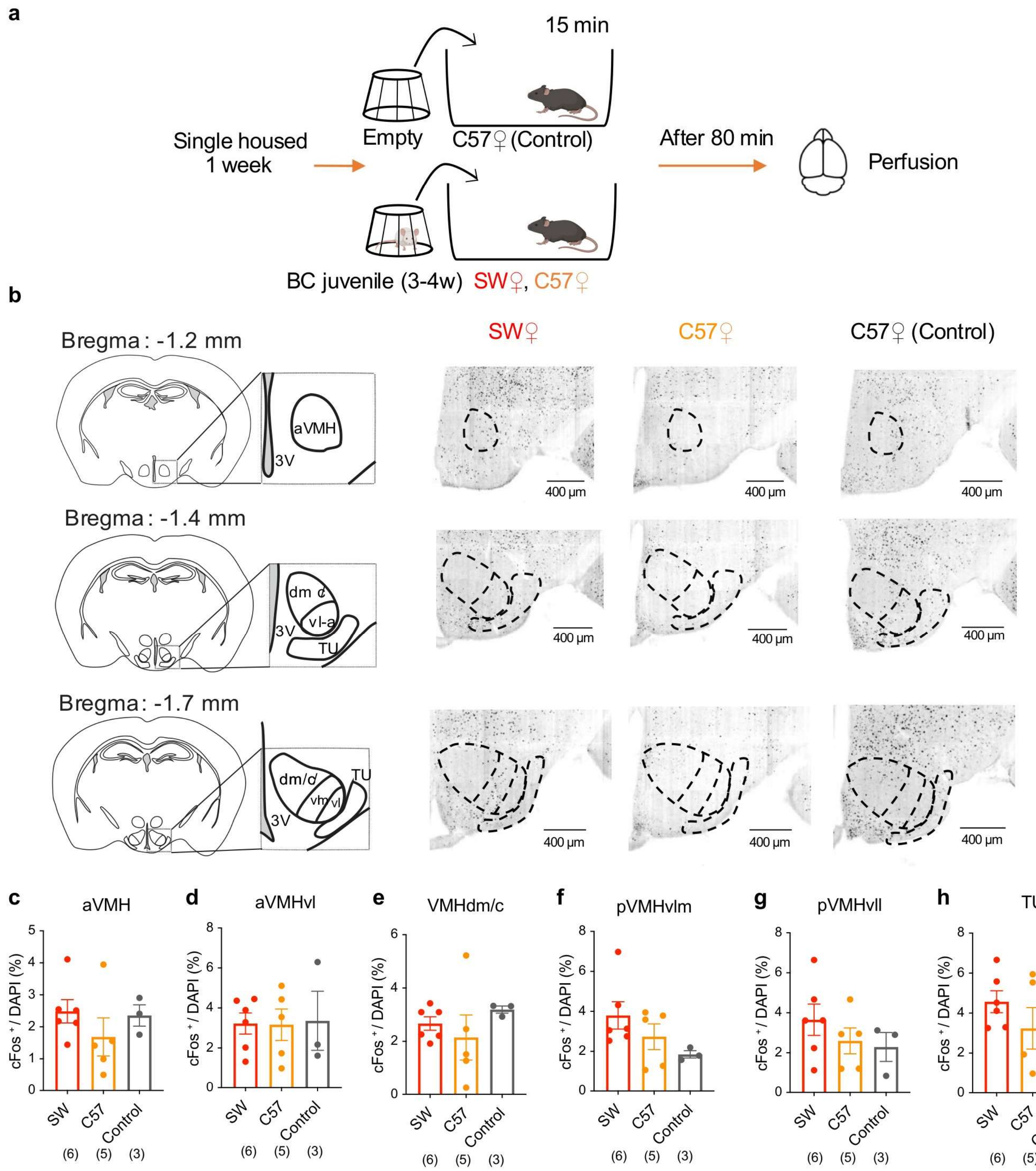


**Supplementary Figure 14. c-Fos expression in females after exposing to a cupped intruder.**

**(a)** Experimental timeline for c-Fos induction. Female mice were single-housed for one week, exposed to a 15-minute cupped male BC juvenile intruder or an empty cup, and perfused 80 minutes later.
**(b)** Left: mouse brain atlas illustrating ventromedial hypothalamus (VMH) subdivisions and tuberal nucleus (TU). Right: coronal brain sections showing intruder-induced c-Fos expression at Bregma levels −1.2, −1.4, and −1.7 mm in representative SW and C57 female mice, and empty cup–induced c-Fos expression in a C57 female mouse. c-Fos–positive cells appear black.
**(c–h)** The percentage of c-Fos–positive cells in **(c)** anterior VMH (aVMH), **(d)** anterior VMHvl (aVMHvl), **(e)** posterior VMH dorsomedial subdivision (pVMHdm), **(f)** posterior VMH ventrolateral medial subdivision (pVMHvlm), **(g)** posterior VMH ventrolateral lateral subdivision (pVMHvll), and **(h)** tuberal nucleus (TU).

Numbers in parentheses indicate animal numbers. Circles represent data of individual animals. Solid circles represent the strain mean, and error bars represent ± SEM. **(c-h)** One-way ANOVA for normally distributed datasets or Kruskal–Wallis test for non-normally distributed datasets with FDR-corrected post hoc comparisons (Benjamini–Krieger–Yekutieli method). All statistical tests are two-tailed. Exact p- or q-value is shown if ≤ 0.05. Otherwise, p- or q-value is unspecified. See **Supplementary Table 1** for additional statistical details.

### Supplementary Figure 15

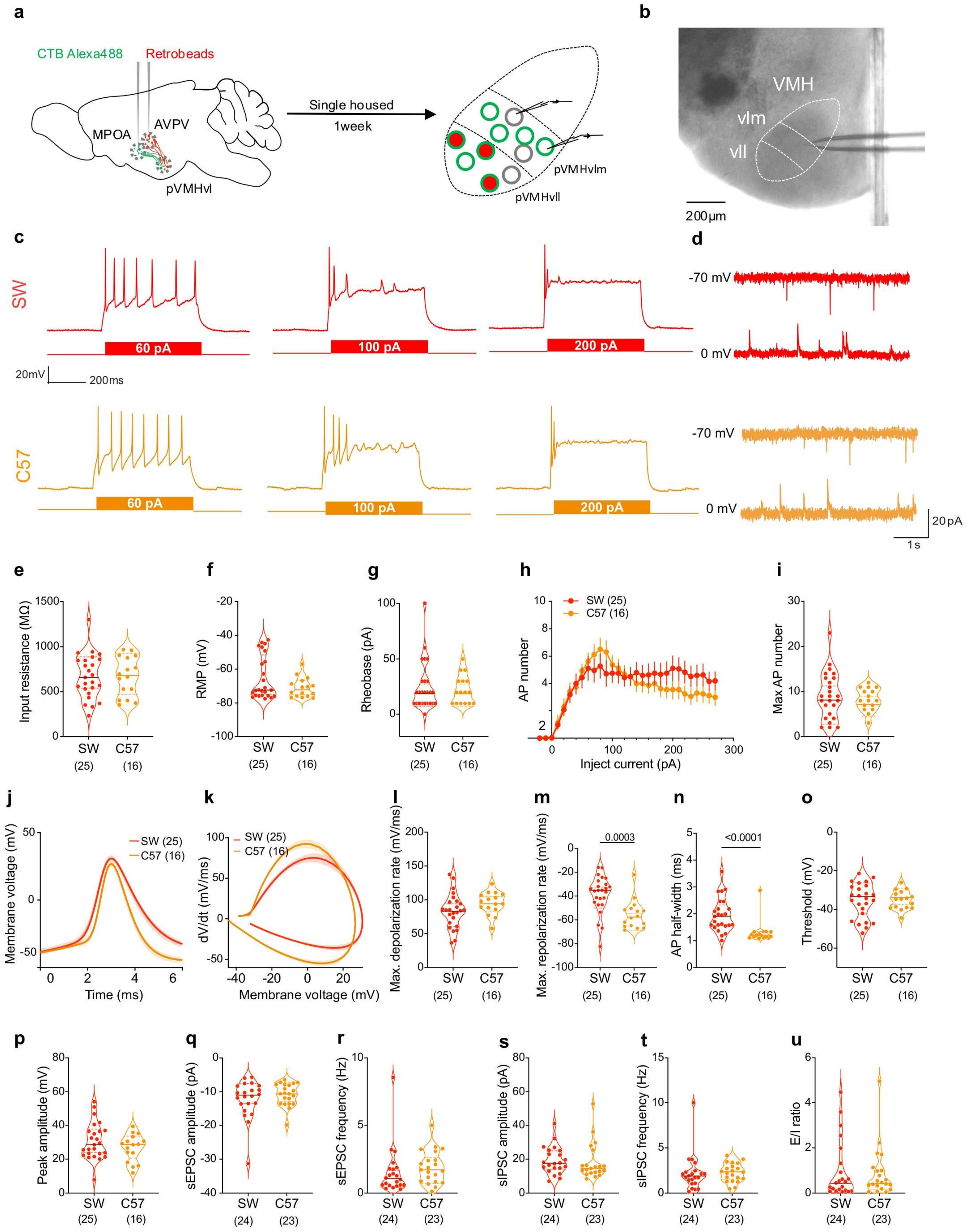

**Supplementary Figure 15. Physiological properties of pVMHvlm cells in female mice**

**(a)** Experimental design for patch-clamp recordings. CTB Alexa-488 was injected into the anteroventral periventricular nucleus (AVPV), and red retrobeads were injected into the medial preoptic area (MPOA) to label neurons projecting to the posterior ventrolateral ventromedial hypothalamus medial subdivision (pVMHvlm). Following post-surgical recovery and one week of single housing, brain slices were prepared, and both Alexa 488-positive and -negative cells were recorded in the green labeling positive and red labeling negative zone.
**(b)** Representative image showing the patched cell location under DIC microscopy.
**(c)** Representative recording traces from pVMHvlm cells in SW (red) and C57 (orange) female mice in response to 60, 100, and 200 pA current injections.
**(d)** Representative traces of sEPSCs (top) and sIPSCs (bottom).
**(e-g)** Intrinsic properties of pVMHvlm cells in C57 and SW females, including **(e)** input resistance, **(f)** resting membrane potential (RMP), and **(g)** rheobase.
**(h)** Spike counts across –20 pA to 270 pA current steps, each for 500 ms.
**(i)** Maximum spike number across SW, C57, and 129 pVMHvlm cells.
**(j)** Average waveform of the first action potential across different current steps in pVMHvl neurons from SW and C57 female mice.
**(k)** Phase plots (dV/dt vs. V) for first spikes across all current steps. Solid lines indicate strain means, shaded areas denote SEM.
**(l-p)** First spike properties, including **(l)** maximum depolarization rate, **(m)** maximum repolarization rate, **(n)** action potential half-width, **(o)** spike threshold, and **(p)** spike peak amplitude.
**(q-u)** Quantification of synaptic activities, including **(q)** sEPSC amplitude, **(r)** sEPSC frequency, **(s)** sIPSC amplitude, **(t)** sIPSC frequency, and **(u)** excitation/inhibition (E/I) ratio.

Numbers in parentheses indicate cell numbers. Circles represent individual cells. Solid lines and shades in **(j, k)** represent means ± SEM. Error bars in **(h)** represent ± SEM. Solid line in (**e-g, i, l-u**) represents the median for each group, while dashed lines indicate quartiles. Cells in **(e-p)** were from 5 SW and 3 C57 female mice, while cells in **(q-u)** were from 4 SW and 4 C57 female mice. **(e-g, i, l-u**) Unpaired t test for normally distributed datasets or Mann–Whitney U test for non-normally distributed datasets. **(h)** Two-way ANOVA with FDR correction. All statistical tests are two-tailed. Exact p- or q-value is shown if ≤ 0.05. Otherwise, p- or q-value is unspecified. See **Supplementary Table 1** for additional statistical details.

### Supplementary Figure 16

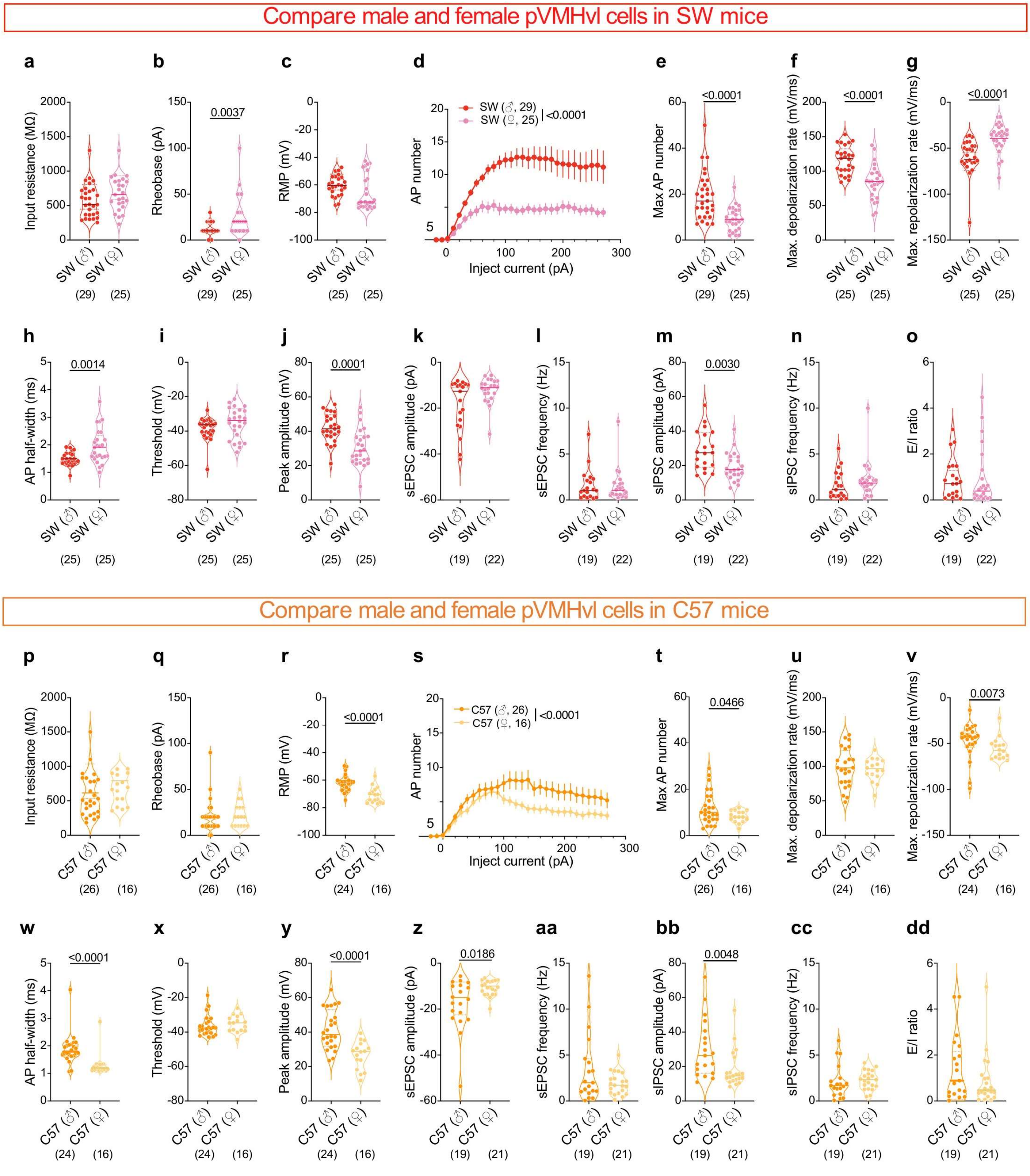

**Supplementary Figure 16. Comparison of electrophysiological properties of VMHvl cells between male and female mice.**

**(a–c)** Intrinsic membrane properties of pVMHvl neurons in male and female SW mice, including **(a)** input resistance, **(b)** rheobase, and **(c)** resting membrane potential (RMP).
**(d)** Spike number as a function of injected current (−20 to 270 pA, 500 ms steps) in male and female SW mice.
**(e)** Maximum spike number evoked across current steps in male and female SW mice.
**(f–j)** First action potential properties in male and female SW mice, including **(f)** maximum depolarization rate, **(g)** maximum repolarization rate, **(h)** action potential half-width, **(i)** spike threshold, and **(j)** peak amplitude.
**(k–o)** Synaptic activity properties in male and female SW mice, including **(k)** sEPSC amplitude, **(l)** sEPSC frequency, **(m)** sIPSC amplitude, **(n)** sIPSC frequency, and **(o)** excitation/inhibition (E/I) ratio.
**(p–r)** Intrinsic membrane properties of pVMHvl neurons in male and female C57 mice, including **(p)** input resistance, **(r)** rheobase, and **(q)** resting membrane potential.
**(s)** Spike number as a function of injected current (0–270 pA, 500 ms steps) in male and female C57 mice.
**(t)** Maximum spike number evoked across current steps in male and female C57 mice.
**(u–y)** First action potential properties for male and female C57 mice, including **(u)** maximum depolarization rate, **(v)** maximum repolarization rate, **(w)** action potential half-width, **(x)** spike threshold, and **(y)** peak amplitude.
**(z–dd)** Synaptic properties, including **(z)** sEPSC amplitude, **(aa)** sEPSC frequency, **(bb)** sIPSC amplitude, **(cc)** sIPSC frequency, and **(dd)** excitation/inhibition ratio.

Numbers in parentheses indicate cell numbers. Circles represent individual cells. Error bars in **(d, s)** represent ± SEM. Solid line in (**a-c, e-o, p-r, t-dd**) represents the median for each group, while dashed lines indicate quartiles. Cells in **(a-o)** were from 5 SW male and 5 SW female mice, while cells in **(p-dd)** were from 5 C57 male and 3 C57 female mice. **(a-c, e-o, p-r, t-dd**) Unpaired t test for normally distributed datasets or Mann–Whitney U test for non-normally distributed datasets. **(h, s)** Two-way ANOVA with FDR correction. All statistical tests are two-tailed. Exact p- or q-value is shown if ≤ 0.05. Otherwise, p- or q-value is unspecified. See **Supplementary Table 1** for additional statistical details.
